# Quantifying the Resource Landscape That Shapes the Human Gut Microbiome

**DOI:** 10.64898/2026.10.01.756135

**Authors:** Jonas Cremer

## Abstract

The microbial ecosystem in the human gut is fundamentally shaped by the resources it receives. Yet, while quantitative information is available for many individual resource fluxes, a comprehensive description of the overall resource landscape is lacking. Here, I systematically integrate physiological measurements and food composition data to map diet and digestion to the fluxes of more than 30 major resources entering the human large intestine, including carbohydrates, proteins, mucus, bile acids, ions, and electron acceptors. Applying this reconstruction across dietary variation in the U.S. population and analyzing the resulting influxes of biomass-forming elements, I find that microbial growth is carbon-limited for most individuals, whereas phosphorus is limiting for a subset; nitrogen, by contrast, is not limiting once host-derived urea recycling is accounted for. Resource supply also determines whether microbial auxotrophies impose physiological constraints. While amino acid dependencies can often be satisfied by protein-derived influx, characterized dietary cobamide influx is insufficient to account for the fecal cobamide pool, pointing to a much greater contribution from microbial production. Together, these results provide a quantitative resource landscape for the human gut, defining physiological constraints on microbial growth and providing a basis for predictive models of microbiome function and its impact on the host.

## 1 Introduction

Elucidating how the dense microbial ecosystem of the human intestine functions and impacts the host requires quantifying the resources available to it. Resources, which I define here broadly as chemical compounds entering the gut, shape microbial physiology, growth, and ecological interactions. As in other ecosystems, their impact depends not only on which resources are present, but on how much of each resource is available [1–3]. Systematically quantifying the influxes of different resources into the large intestine is therefore essential for determining the physiological and ecological constraints under which the microbiome operates.

Resource influx also shapes the impact of the microbiome on the host, as the magnitude and composition of chemical fluxes from the microbiome to the host tightly depend on the resources available to it. For example, the production of fermentation products such as butyrate, lactate, and acetate depends strongly on the influx of carbohydrates and other fermentable substrates into the large intestine [4–6]. Resource availability similarly affects the production of other microbial metabolites exchanged with the host, including potentially harmful metabolites [7, 8]. More generally, as recently demonstrated, how molecular fluxes from the microbiome to the host depend on microbial abundance, biomass, community composition, and per-cell metabolic activity [9]. Here, I focus on how resource influx into the gut shapes microbial growth, activity, and community composition.

Much of the resource influx into the gut originates from diet, with dietary inputs transformed by digestion, absorption, and transport along the digestive tract, while additional resources enter the gut through secretion by the host epithelium. Extensive experimental work has established how different nutrient conditions, particularly the availability of carbohydrates and proteins, affect gut microbial growth, metabolism, and community composition, both in vitro and in animal models [3, 10–16]. In contrast, the influxes of these resources into the human large intestine have received far less systematic quantitative analysis [17, 18].

Because the vast majority of gut microbial biomass resides in the large intestine [19, 20], I focus on resource influxes into this part of the gut. By integrating diverse datasets on diet composition, digestion, absorption, and host physiology, I establish a *quantitative resource landscape* of the healthy adult large intestine, systematically reconstructing the influxes of more than 30 major resources, including different types of carbohydrates, proteins, lipids, mucus, bile acids, electron acceptors, and major ions.

For each resource, I assess the quantitative confidence of both its influx estimate and its dependence on diet, identifying where estimates are well constrained by available measurements and where major uncertainties remain. I then apply the quantitative mapping between food intake and resource influx to dietary intake data from NHANES to demonstrate how the resource landscape varies across the U.S. population.

With this quantitative resource landscape, I then analyze which biomass-forming elements limit microbial growth in the gut. Comparing the influxes of carbon, nitrogen, phosphorus, and sulfur with their requirements for biomass production identifies the conditions under which each element becomes limiting and reveals how host-derived urea recycling alters these constraints. I further use the reconstructed resource influxes to examine microbial auxotrophies in their quantitative environmental context, investigating whether essential metabolites can be supplied by diet and the host or instead require microbial production and exchange. I apply this supply–demand perspective to amino acids and cobamides, two widespread classes of microbial dependencies with distinct patterns of environmental supply [21–23].

## 2 Results

### 2.1 The resource landscape of the large intestine

The reconstructed influxes of more than 30 resources are shown in Figure 1A and Table S1. Resources were selected based on their quantitative contribution to the overall influx into the large intestine and their relevance to microbial growth and ecology, with lower-flux resources such as cobalamin and oxygen included because of their specific physiological importance. The major types of physiological evidence supporting each estimate are summarized in Figure 1AB. The full analyses underlying each influx reconstruction, including the integrated measurements, assumptions, and supporting literature, are provided in the supplementary text (SI Text 2–7).

**Figure 1:**
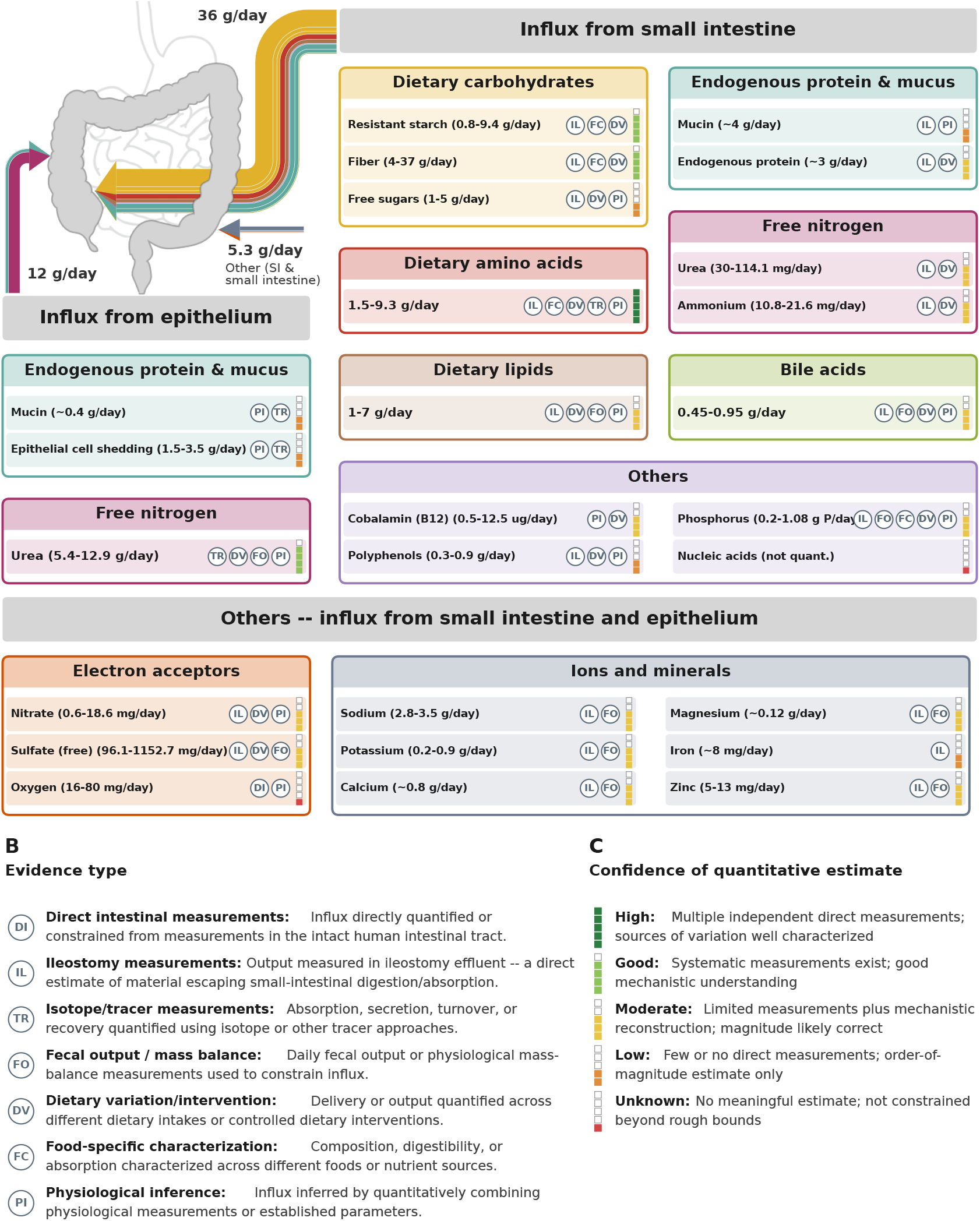
Quantitative reconstruction of the resource landscape encountered by the human gut microbiome. (A) Estimated daily influxes of more than 30 major resources into the healthy adult large intestine, including dietary and host-derived nutrients, major ions, important trace metals, and electron acceptors. Resources enter from the small intestine or are supplied directly by the host epithelium. Ranges represent estimated physiological variation, and symbols indicate both the major type(s) of evidence underlying each estimate and its assigned confidence. (B) Evidence types underlying each influx estimate, as badged in (A). (C) Assigned confidence with which the magnitude and variation of each resource influx can currently be quantified, based on the available evidence. The choice of analyzed resources and details on influx estimation, including the available evidence for each resource, are provided in SI Text 2–7. Resource influxes are summarized in Table S1. Evidence types and confidence levels are defined in Table S2.

To assess the reliability of each resource influx estimate and its dependence on diet, I assigned five confidence levels (unknown, low, moderate, good, and high; Figure 1C and Table S2; Methods). These levels reflect the availability and quality of quantitative measurements rather than the ecological importance of a resource. Dietary macronutrient influxes are generally estimated with the highest confidence, reflecting extensive measurements of intestinal digestion and absorption in humans. In contrast, estimates for host-derived secretions and electron acceptors tend to rely more strongly on indirect physiological reconstructions and are generally estimated with lower confidence. This classification highlights which parts of the resource landscape are well constrained and where additional measurements would be most informative.

Most resources available to the microbiome in the large intestine enter from the small intestine, including dietary carbohydrates, proteins, lipids, bile acids, most major ions, and trace metals (Fig. 1A, SI Text 2–5). For many of these resources, multiple independent physiological estimates are available with good agreement. Carbohydrate influx, estimated at 8–49 g/day (combining resistant starch, fiber, and free sugars), is supported by estimates from dietary intake and digestibility, ileostomy measurements, and fecal mass-balance studies. Protein influx, estimated at 1.5–9.3 g/day, is supported by food-specific ileal digestibility measurements and direct ileal effluent analyses [24, 25] (SI Text 2.2). Lipid influx is estimated at 1–7 g/day with comparatively less known about the molecular composition of lipids reaching the large intestine (SI Text 2.3). Bile acid influx, estimated at 1–2 mmol/day, can be reconstructed from physiological turnover and ileal-loss measurements (SI Text 3.3).

In addition to dietary and endogenous material entering from the small intestine, the host supplies resources directly to the large intestine through secretion and epithelial turnover. Urea influx is estimated at 90–215 mmol/day, based largely on isotope-labeling measurements of urea turnover and hydrolysis. Mucus secretion and epithelial-cell turnover provide additional host-derived organic material, with estimated influxes of approximately 0.4 g/day and 1.5–3.5 g/day, respectively (SI Text 6).

Electron acceptors constitute a distinct class of resources as they enable respiration and thereby shape microbial metabolism [26] (SI Text 7). Nitrate and sulfate enter from the small intestine at approximately 0.01–0.3 and 1–12 mmol/day, respectively, based on direct physiological measurements. Oxygen influx is more difficult to quantify but available concentration measurements and diffusion calculations give an upper bound estimation of 0.5–2.5 mmol/day (SI Text 7.1).

### 2.2 Diet-dependent variation in resource influx

Having reconstructed the typical influx ranges of the different resources, I next investigated how the influxes of major resources vary with diet and across individuals. To quantify this variation, I developed a mapping from dietary intake to large-intestinal resource influx, ranging from simple fractional estimates to detailed food-level models depending on the resource and data available, integrating food composition with resource- and food-specific estimates of digestion and absorption (Figure 2A, SI Text 8). I applied this mapping to dietary records from NHANES [27] to reconstruct the distribution of major resource influxes across the U.S. population.

**Figure 2:**
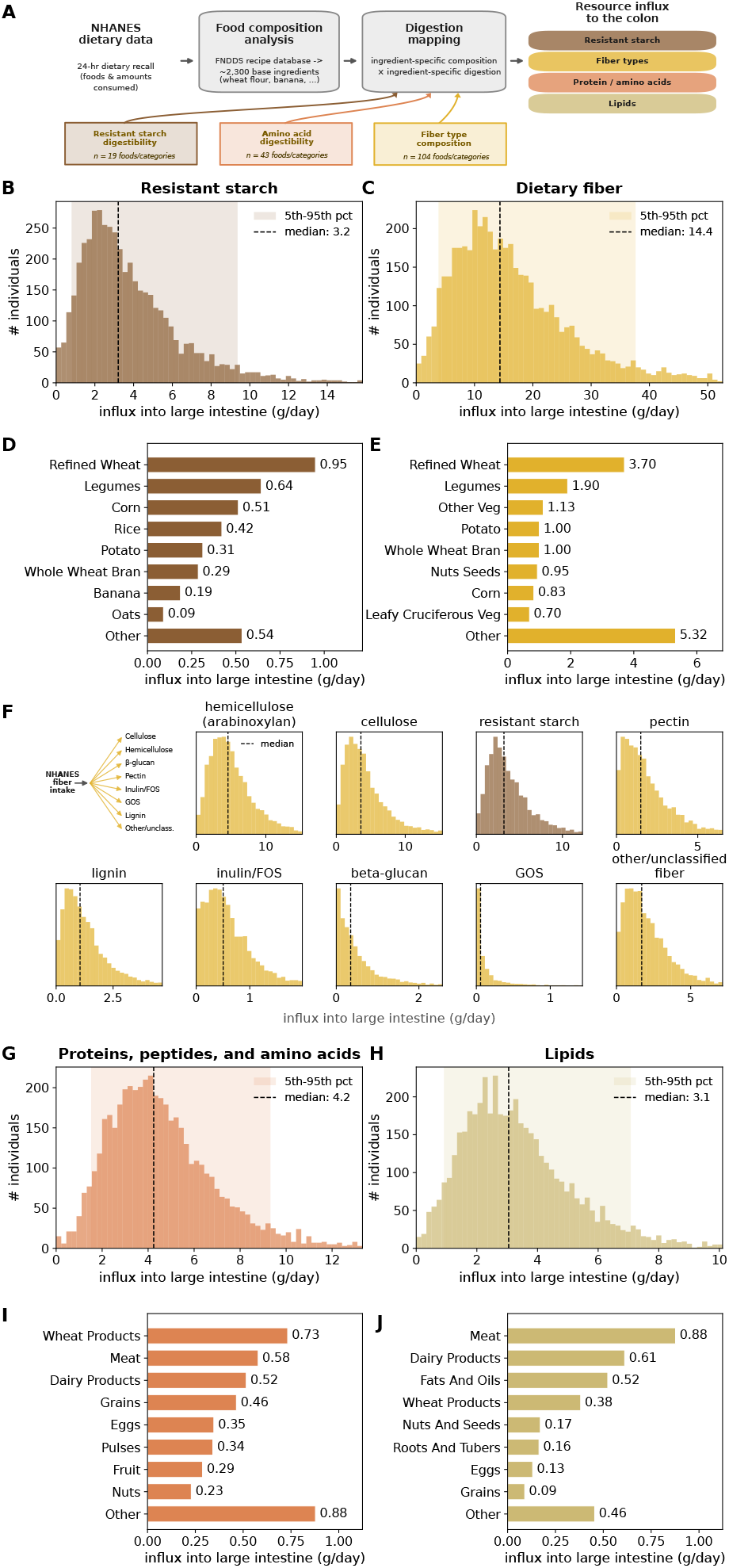
Diet-dependent variation of the resource landscape. (A) Implemented mapping of dietary intake to large-intestinal resource influx, integrating NHANES dietary records for the US population and food composition with resource- and food-specific estimates of small-intestinal digestion and absorption, extending from simple flat-fraction to detailed food-level models depending on the resource (SI Text 8). (B,C) Population distributions of the estimated daily influx of resistant starch (B) and dietary fiber (C). (D,E) Contributions of major food groups to resistant starch (D) and dietary fiber (E) influx. (F) Composition of the carbohydrate influx reaching the large intestine, including the contributions and population distributions of major dietary fiber types. (G,H) Population distributions of total amino acid (G) and lipid (H) influx. Amino acid influx includes free amino acids, peptides, and proteins. (I,J) Contributions of major food groups to total amino acid (I) and lipid (J) influx into the large intestine. Amino acid and carbohydrate flux estimates accounted for food-specific digestion and composition values. Lipid influx was estimated assuming a constant fractional escape from small-intestinal absorption. The analyses shown use all NHANES dietary records without survey weighting. Applying NHANES survey weights to better represent the U.S. population resulted in only modest changes in the reconstructed distributions of dietary intake and large-intestinal resource influx (Figure S2).

Complex carbohydrate influx varies substantially across individuals, with resistant starch ranging from 1–8 g/day and dietary fiber from 5–31 g/day between the 10th and 90th percentiles (Figure 2B,C). To determine the dietary origins of this variation, I extended a recent reconstruction of carbohydrate influx [6] to account explicitly for food-specific fiber and resistant starch contents (Figure S1A,B, SI Text 2.1.1–2.1.2).

Resistant starch and fiber show distinct contributions from major food groups (Figure 2D,E). The composition of the fiber influx also varies across individuals, with typical (median) daily intake across individual fiber types spanning roughly an 80-fold range from well under 0.1 g/day to, in the case of hemicellulose, more than 4 g/day (Figure 2F). Because different fiber types differ in their microbial utilization, this variation changes the composition of carbohydrate resources available to the microbiome.

Dietary protein-derived amino acid influx varies from 2–8 g/day between the 10th and 90th percentiles, with a median of 4 g/day (Figure 2G). To capture differences in amino acid composition and ileal digestibility among dietary proteins such as animal and protein derived proteins, the mapping integrates food-specific digestibility measurements with nutrient composition databases (Figure S1C). This influx originates from a broad range of food sources, with wheat products, meat, and dairy providing the largest contributions at the population level (Figure 2I).

Lipid influx ranges from 1–6 g/day across the central 80% of the population, with a median of approximately 3 g/day (Figure 2H). I estimate total dietary lipid influx assuming a constant fractional escape from small-intestinal absorption rather than resolving individual lipid classes, since considerably less is known about food-specific differences in lipid absorption or the molecular composition of lipids reaching the large intestine, including the relative contributions of triglycerides, phospholipids, and individual fatty acids. The contributions of different food sources are shown in Figure 2J.

### 2.3 Growth limitation

I next asked which biomass-forming elements constrain potential microbial biomass production in the healthy human large intestine, focusing on the supply of carbon, nitrogen, phosphorus, and sulfur relative to their requirements for biomass synthesis. Carbon and nitrogen have received particular attention as potential limiting resources in the large intestine. Carbohydrates provide both carbon for biomass synthesis and energy for growth and can therefore readily become limiting in predominantly anaerobic heterotrophic ecosystems [28, 29]. Nitrogen availability can likewise shape microbial community composition [30]. Comparisons of carbon-to-nitrogen ratios in diet and feces together with protein-supplementation experiments in mice have further suggested that microbial growth in the gut can be nitrogen limited [31]. Whether carbon or nitrogen limits microbial growth in the human large intestine, however, remains debated [32].

To address this question, I compared the influxes of carbon (C), nitrogen (N), phosphorus (P), and sulfur (S) with their requirements for biomass synthesis, calculating for each element the amount of microbial biomass that could be supported by its influx into the large intestine. Although microbial growth follows highly dynamic spatiotemporal patterns shaped by varying local resource availability, these influx-based bounds can reveal which elemental supplies fundamentally constrain biomass production.

To do so, I first converted the typical values of all reconstructed resource influxes into their elemental components (SI Text 9 and Table S3) and complemented these estimates with total P measurements (Figure S3 and SI Text 4.3), yielding daily influx estimates of C, N, P, and S into the large intestine (Figure 3A). For the further analysis, I only considered resources that can be utilized by the microbiome (Figure 3A, filled bars), excluding lipids, poorly utilized fiber fractions, and other resources with limited availability for microbial growth (SI Text 9.5 and Table S3). The contribution of individual resources to the growth-accessible influx of each element, based on NHANES dietary records, is shown in Figure 3B.

**Figure 3:**
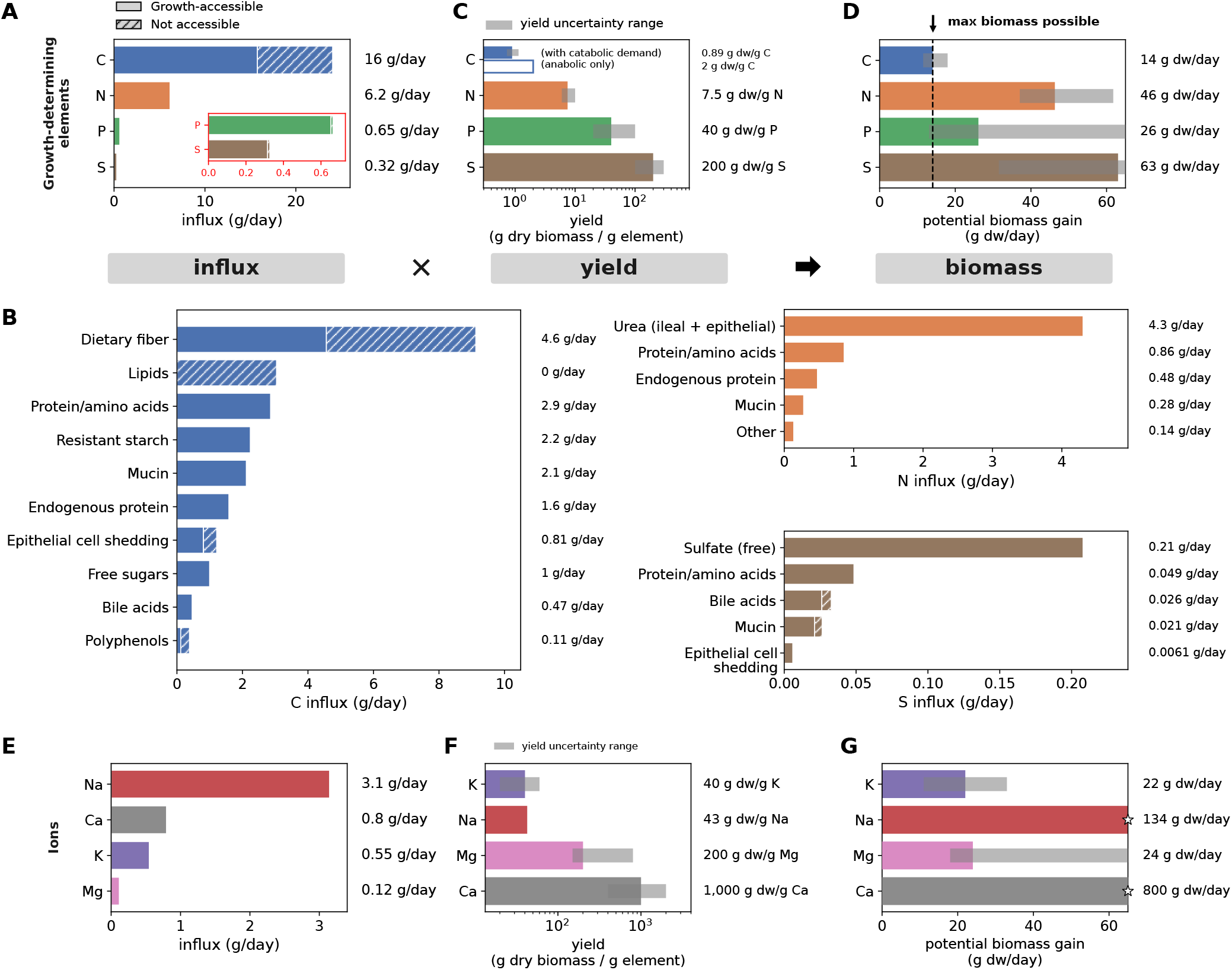
Elemental resource constraints on microbial biomass production. (A) Estimated daily influxes of the major biomass-forming elements carbon (C), nitrogen (N), phosphorus (P), and sulfur (S) into the healthy adult human large intestine. Hatched fractions indicate influxes considered poorly accessible or unavailable for microbial growth and are excluded from subsequent growth estimations. Inset in red shows an expanded view of P and S. (B) Contributions of individual dietary and host-derived resources to the growth-accessible influxes of C, N, and S. (C) Element-specific biomass yields for C, N, P, and S, assuming that other biomass requirements are met. Gray ranges indicate plausible variation based on available measurements and physiological conditions. For carbon, the anabolic biomass requirement is distinguished from the total carbon requirement, which additionally includes catabolic carbon consumption. (D) Potential microbial biomass production supported by the growth-accessible influx of each element. Gray ranges reflect variation based on variation in biomass yields in C. Black dashed line indicates the biomass supported by carbon, the most constraining element. Phosphorus is the next most constraining element. (E) Estimated daily influxes of the ions magnesium (Mg), calcium (Ca), sodium (Na), and potassium (K). (F) Representative cellular ion quotas expressed as apparent biomass yields, with gray ranges indicating variation across measurements and physiological conditions. (G) Biomass corresponding to the estimated growth-accessible ion influxes and representative cellular ion quotas. Unlike in (D), these values should not be interpreted as strict growth-limiting capacities because cellular requirements for these ions are flexible and depend on extracellular conditions.

To convert elemental influxes into potential microbial biomass synthesis, I next compiled available measurements of biomass yields *Y*_*X*/*E*_, defined as the potential dry biomass *X* that can be synthesized per mass of element *E* supplied when *E* alone limits growth (SI Text 10.1 and Table S4). Typical values (colored bars) and reported ranges (gray bars) are shown in Figure 3C. For N, P, and S, yields are largely determined by anabolic demands to make new biomass and the elemental composition of cells (SI Text 10.1) [33, 34]. Carbon is different because, beyond satisfying anabolic demands, carbon substrates also provide energy for growth, making biomass yield dependent on catabolic metabolism [28, 29]. Given the comparatively small influx of respiratory electron acceptors relative to fermentable substrates, energy generation from carbohydrate catabolism in the large intestine is dominated by fermentation (SI Text 10.1). To obtain yields appropriate for the gut microbiome, I used recent systematic measurements of carbohydrate consumption across major bacterial species in the human gut [6], while accounting for the contribution of amino acids to anabolic biomass synthesis under the measured growth conditions (SI Text 10.1). Consistent with substantial catabolic carbon consumption, the resulting biomass yields are considerably lower than those inferred from anabolic carbon demand alone (Figure 3C, blue bars).

To obtain the potential microbial biomass production supported by each element, I then multiplied each element’s influx by the corresponding yield (Figure 3D). For the representative resource influxes and yield values considered here, carbon supports the least biomass, approximately 14 g dry biomass day^*−*1^, making it the most constraining element. This value closely matches estimates of daily microbial biomass loss in feces across different human populations, which range from approximately 12–20 g dry biomass day^*−*1^, with the lower values observed for low-fiber diets typical of the U.S. population [6].

Phosphorus is the next most constraining element, supporting approximately 26 g dry biomass day^*−*1^, whereas nitrogen and sulfur can support substantially greater biomass production. Carbon remains the most constraining element across much of the reported variation in biomass yields (Figure 3D, gray bars). I also compared sodium, potassium, magnesium, and calcium influxes with representative cellular quotas (Figure 3E–G), indicating that these ions are unlikely to limit biomass production. Moreover, unlike the elemental requirements considered above, cellular ion content is more variable, with flexible ion homeostasis placing weaker constraints on growth [35–37].

The apparent excess of nitrogen relative to carbon is largely due to nitrogen recycling between the host and gut microbiome: In addition to nitrogen entering the large intestine directly through dietary and endogenous proteins and amino acids that escape small-intestinal absorption, the microbiome receives substantial amounts of host-derived nitrogen through urea secreted across the intestinal epithelium (Figure 4A, SI Text 6.2) [38–40], corresponding to approximately 180–430 mmol N day^*−*1^. Microbial ureases hydrolyze urea to ammonium, making this nitrogen available for microbial amino acid synthesis and growth [40, 41].

**Figure 4:**
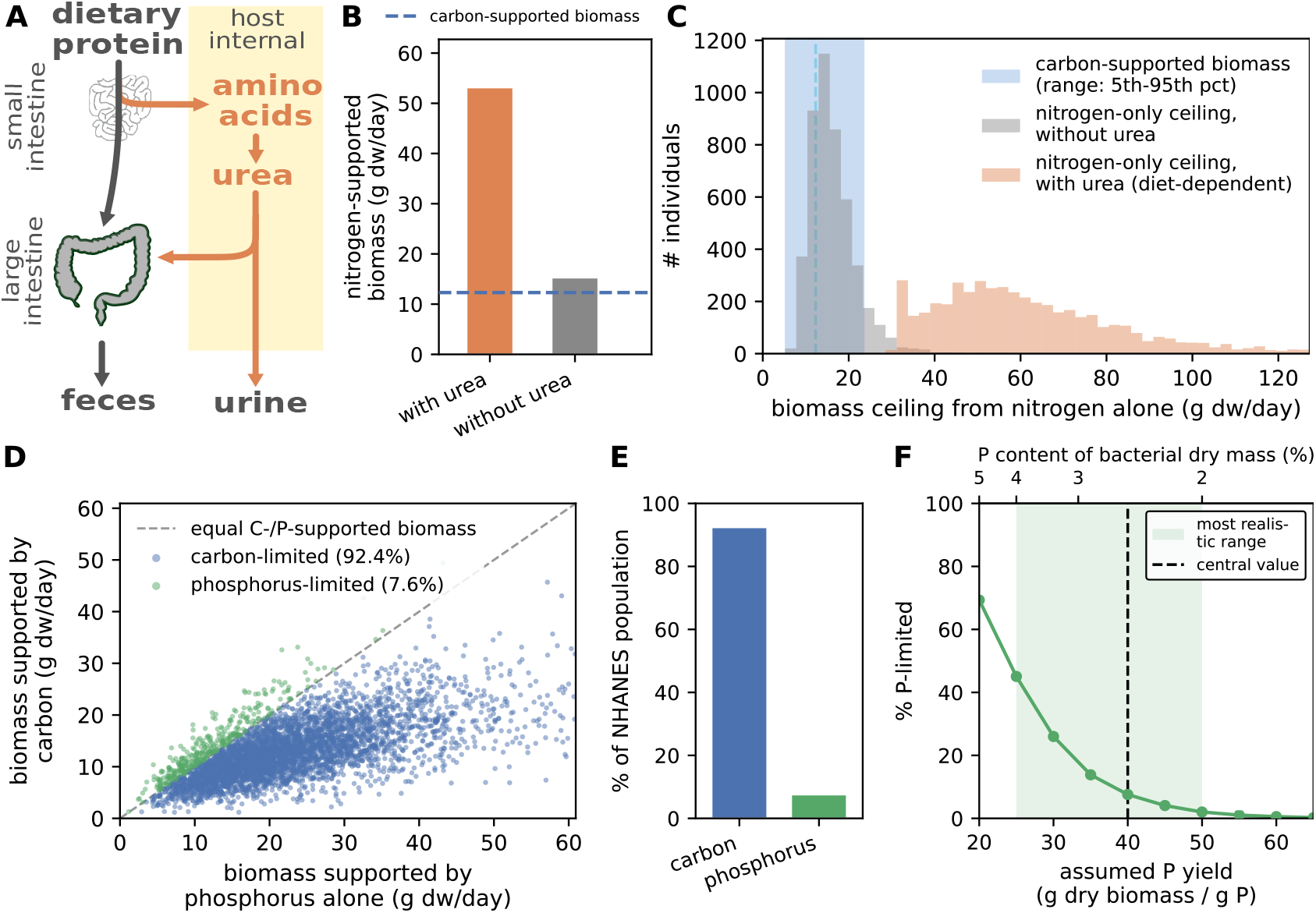
Host urea recycling relieves nitrogen limitation and diet determines carbon versus phosphorus limitation. (A) Schematic of host nitrogen fluxes contributing to urea production and excretion. Dietary amino acids contribute to host urea production, with urea subsequently excreted in urine or released into the gastrointestinal tract, where microbial urease activity converts urea into ammonium available for microbial assimilation. (B) Predicted microbial biomass that can be supported by nitrogen influx into the large intestine, with and without host-derived urea influx. In the absence of urea influx, nitrogen-supported biomass is comparable to carbon-supported biomass (blue dashed line), whereas inclusion of urea substantially increases nitrogen-supported biomass. (C) Distribution of nitrogen-supported microbial biomass across individuals in the U.S. population. Individual estimates account for variation in dietary protein intake and protein digestibility; host-derived urea influx is estimated using a mapping in which urea delivery to the large intestine depends on dietary protein intake (SI Text 6.2). Without urea influx (gray), nitrogen-supported biomass overlaps substantially with the range of carbon-supported biomass (blue shaded region), whereas inclusion of urea shifts the distribution toward higher values. (D) Comparison of microbial biomass that can be supported by carbon and phosphorus influx across individuals in the U.S. population. The diagonal indicates equal C- and P-supported biomass and separates individuals predicted to be carbon- and phosphorus-limited. (E) Fraction of individuals in D predicted to be carbon- or phosphorus-limited. (F) Sensitivity of the fraction of individuals predicted to be phosphorus-limited to the assumed phosphorus biomass yield. Individual resource influxes in (C–E) were reconstructed from NHANES dietary data together with estimates of food-specific digestion and host-derived resource influxes.

When considering only nitrogen entering directly through proteins and amino acids, the potential biomass supported by nitrogen is comparable to that supported by carbon (Figure 4B, dashed line). Including host-derived urea substantially increases nitrogen availability and the corresponding nitrogen-supported biomass (Figure 4B). Extending this comparison across individuals, I combined NHANES dietary variation with food-specific protein digestibility and carbohydrate supply. Without urea influx, the distributions of nitrogen- and carbon-supported biomass overlap substantially, and nitrogen is the more constraining element for approximately 19% of individuals. With urea influx included, nitrogen-supported biomass exceeds carbon-supported biomass across essentially all individuals (Figures 4C and S4A–D). Host urea recycling therefore prevents nitrogen from becoming the more constraining elemental supply across individuals.

Finding that nitrogen is not limiting does not, however, imply that microbial growth is universally carbon limited. Across individuals, variation in resource influx changes the relative biomass supported by carbon and phosphorus (Figure 4D). For the representative yield values, carbon is the more constraining element for approximately 92% of individuals, whereas phosphorus is more constraining for the remaining 8% (Figure 4D,E).

In these calculations, I used a fixed phosphorus yield. However, cellular phosphorus content varies physiologically with growth rate, in large part because RNA content increases in rapidly growing cells to synthesize more ribosomes, thereby increasing phosphorus demand [42, 43]. Consequently, the fraction of individuals predicted to be phosphorus-limited depends strongly on the assumed phosphorus yield. Varying the phosphorus biomass yield across the range observed in available measurements changes the predicted fraction of phosphorus-limited individuals from effectively 0% to approximately 69% (Figures 4F and S4E). The prevalence of phosphorus limitation in the human gut therefore remains poorly constrained. Because phosphorus yield depends on growth rate, its systematic measurements across gut species and culture conditions will be particularly important for determining whether phosphorus limitation is rare or widespread.

### 2.4 Resource supply and microbial auxotrophies

I next asked how resource influx shapes microbial auxotrophies, the inability to synthesize one or more metabolites required for growth. Auxotrophies for amino acids, vitamins, and other cellular components are widespread among gut bacteria [44, 45] and are commonly associated with metabolic exchange between community members [44, 46]. In the gut, however, an auxotrophic requirement can also be met directly by metabolites supplied through diet or by the host. Whether an auxotrophy creates a dependence on microbial cross-feeding therefore depends on the balance between cellular demand, external resource influx, and microbial production. Quantifying this balance is also important for understanding the physiological and ecological consequences of auxotrophy and its evolutionary origin.

I first applied this supply–demand perspective to amino acids. While protein synthesis is an essential component of microbial growth, with proteins constituting a large fraction of cellular biomass and accounting for a major fraction of cellular nitrogen [43], many gut bacterial strains lack the ability to synthesize one or more amino acids [44]. However, substantial amounts of protein, peptides, and amino acids enter the large intestine and can provide amino acids directly for protein synthesis, thereby reducing the amount that must be synthesized de novo by the microbial community (Figure 5A).

**Figure 5:**
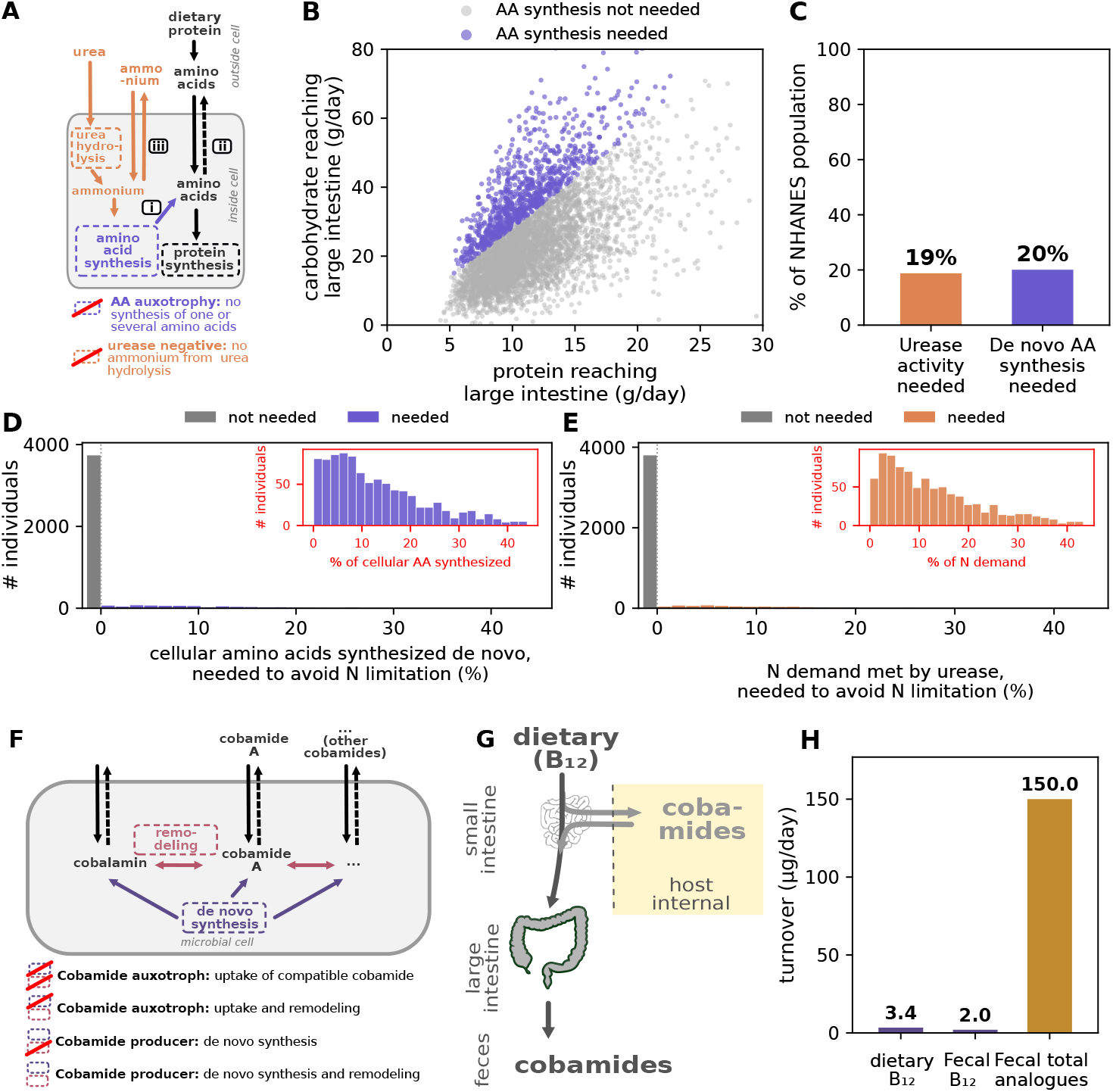
Resource influx shapes the physiological consequences of microbial auxotrophies. (A) Alternative routes supplying amino acids and nitrogen for microbial growth. Protein synthesis creates a cellular demand for amino acids (i), which can be met through uptake of amino acids entering the large intestine or de novo amino acid synthesis (ii). Host-derived urea provides an additional nitrogen source for de novo amino acid synthesis following urease-mediated hydrolysis to ammonium (iii). (B) Requirement for de novo amino acid synthesis across individuals in the U.S. population, determined by comparing the total anabolic amino acid demand of carbon-supported microbial biomass production to the estimated amino acid influx into the large intestine. (C) Fraction of individuals for whom external amino acid supply is insufficient to meet total anabolic amino acid demand or for whom urea-derived nitrogen is required to meet total anabolic nitrogen demand. (D) Minimum fraction of total anabolic amino acid demand requiring de novo synthesis across the NHANES population. Gray indicates individuals for whom no de novo synthesis is required; the colored distribution indicates individuals with a nonzero requirement. The inset enlarges the colored distribution. (E) Minimum fraction of total anabolic nitrogen demand requiring urea-derived nitrogen following urease-mediated hydrolysis across the NHANES population. Gray indicates individuals for whom urea-derived nitrogen is not required; the colored distribution indicates individuals with a nonzero requirement. The inset enlarges the colored distribution. The calculations in (B–E) compare resource supply and anabolic demand at the community level and assume that the total amino acid influx reaching the large intestine, including amino acids contained in proteins and peptides, is accessible for microbial utilization. They therefore provide lower-bound estimates of the required de novo amino acid synthesis and urea-derived nitrogen. Individual resource influxes underlying (B–E) were reconstructed as in Figure 2. (F) Cobamide acquisition and production strategies. Cobamide requirements can be met through uptake of compatible extracellular cobamides, remodeling of imported cobamides, or de novo synthesis. (G) Schematic of dietary cobalamin (vitamin B_12_) intake, host absorption and enterohepatic recycling, and entry into the large intestine. (H) Comparison of the estimated cobalamin influx into the large intestine with fecal cobalamin and total corrinoid output. Dietary influxes of corrinoids other than cobalamin are poorly characterized, but their low levels in available food-composition data suggest that they cannot account for the large fecal corrinoid output.

To quantify the extent to which amino acid supply from the host can meet anabolic demand, I compared amino acid influx into the large intestine with the demand for microbial biomass synthesis. For each individual in the NHANES population, I calculated the anabolic nitrogen demand of the carbon-supported microbial biomass estimated above and compared it with the corresponding amino acid influx. External amino acid supply is insufficient to meet total anabolic demand in approximately 20% of individuals, requiring the remaining demand to be met by de novo amino acid synthesis within the microbial community (Figure 5B,C). Among these individuals, community-level de novo synthesis typically accounts for approximately 10% of total anabolic amino acid demand (Figure 5D).

The same supply–demand analysis can be extended to host-derived urea nitrogen. Urease hydrolyzes urea to ammonia/ammonium, providing assimilable nitrogen for de novo amino acid synthesis (Figure 5A). Because urease is encoded by only a subset of gut bacteria [40, 47], many community members cannot directly access this nitrogen source. I therefore recalculated the biomass ceiling for each individual without urea-derived nitrogen to determine when urea-derived nitrogen is required to meet anabolic nitrogen demand. Across the NHANES population, urea-derived nitrogen is required in approximately 19% of individuals (Figure 5C) and, among these individuals, typically accounts for less than 20% of total anabolic nitrogen demand (Figure 5E).

My analysis assumes that the entire amino acid influx reaching the large intestine, including amino acids contained in proteins and peptides, can ultimately be utilized by the microbial community. However, while many gut bacteria exhibit extracellular protease activity [48], peptide hydrolysis and subsequent amino acid uptake vary strongly among taxa [49]. My estimates therefore provide a lower bound on the requirement for de novo amino acid synthesis and access to urea-derived nitrogen. Furthermore, the analysis considers total amino acid supply and demand at the community level and does not resolve individual amino acids or microbial taxa. Resolving these dependencies will require amino-acid-specific supply and demand together with taxon-specific biosynthetic capabilities, uptake, abundance, and metabolic exchange. Nevertheless, the substantial influx of protein-derived amino acids shows that external amino acid supply can satisfy a large fraction of microbial anabolic amino acid demand, potentially reducing the need for de novo synthesis and facilitating the widespread occurrence of amino acid auxotrophies in the gut.

The supply–demand perspective extends to vitamins and cofactors. Cobamides provide a useful contrast to amino acids. Cobalamin (vitamin B_12_) is one member of this broader family of structurally related cofactors. While cobamide-dependent metabolism is widespread among gut bacteria, complete de novo cobamide synthesis is restricted to a smaller subset of organisms [50] (SI Text 13.2). Cobamide requirements can instead be met through uptake of a compatible extracellular cobamide or remodeling of a different cobamide (Figure 5F) [22, 51]. These different strategies for cobamide acquisition and utilization shape microbial communities: cobamide identity affects competitive fitness in vitro, individual corrinoid transporters determine fitness in the mammalian gut, and microbial production and cross-feeding can sustain cobamide-dependent community members [22, 23, 52].

The consequences of cobamide auxotrophy should depend strongly on external cobamide supply. Although a substantial fraction of dietary cobalamin is absorbed in the small intestine, cobalamin is also continuously secreted by the host into bile (Figure 5G; SI Text 2.5.1). Over sufficiently long timescales, average dietary cobalamin intake thus balances cobalamin loss from the body. Because enterohepatic cycling redistributes cobalamin without providing a net source, the average cobalamin flux reaching the large intestine is ultimately constrained by dietary intake and is therefore expected to be of similar magnitude.

Comparing this estimated cobalamin influx with corrinoid output in feces [53, 54] reveals a substantial imbalance: the estimated cobalamin influx is approximately 3 *µ*g/day, compared with 2 *µ*g/day of cobalamin and 150 *µ*g/day of total corrinoids in feces (Figure 5H). Thus, fecal cobalamin output alone is comparable to the estimated influx, whereas total fecal corrinoid output is approximately 44-fold higher. Although dietary influxes of non-canonical cobamides remain insufficiently characterized, available food-composition data suggest that they are unlikely to account for this excess. The flux imbalance therefore points to substantial net microbial corrinoid production at the community level.

Cobamides thus contrast with amino acids: whereas protein-derived amino acid influx can satisfy a large fraction of community anabolic demand, external cobamide influx is small relative to fecal corrinoid output, pointing to substantial microbial production.

## 3 Discussion

Understanding microbial growth in the human large intestine requires knowing which resources reach the microbiome and in what quantities. Here, I reconstructed these resource fluxes by integrating physiological measurements across nutrient digestion, intestinal transport, and whole-body metabolism. Many of these measurements were originally obtained to address physiological and nutritional questions independent of microbial ecology, such as protein digestion in malnutrition or intestinal water and electrolyte homeostasis. Together, they provide a quantitative mapping between diet and gastrointestinal physiology and the resources available to the microbiome. This reconstruction identifies carbon as the dominant constraint on microbial biomass production under representative conditions, although phosphorus can also become limiting. In contrast, host urea recycling provides substantial additional nitrogen and prevents nitrogen from becoming limiting.

Resource influx also shapes the physiological consequences of microbial auxotrophies. Protein-derived amino acid influx can satisfy total community-level anabolic demand for most individuals if these resources are accessible to the microbiome. In contrast, external cobalamin influx is small relative to microbial cobamide requirements, highlighting the greater dependence of cobamide auxotrophs on microbial production and exchange. These influx analyses extend naturally to other common biosynthetic dependencies, including heme, additional vitamins and cofactors, and nucleotides, wherever environmental supply can be quantitatively compared with cellular demand. Such supply–demand comparisons could identify which dependencies can be met by environmental supply and which instead require microbial production or exchange, while highlighting where key quantitative measurements are still lacking.

More broadly, the physiological resource landscape can be understood as a quantitative boundary condition connecting diet and host physiology to microbial ecology. As in all biological systems, the behavior of a microbial ecosystem depends not only on its internal components and processes but also on the fluxes of matter and energy across its boundaries. Sequencing, transcriptomics, and metabolomics provide increasingly detailed views of community composition, metabolic capabilities, and activity: which species and genes are present, and which metabolic transformations and pathways are possible [55]. These properties alone, however, do not determine which metabolic processes are realized or the magnitude of microbial growth. Both also depend on the resources entering the system. Composition, metabolic capability, and resource influxes should therefore not be viewed as competing descriptions of the gut microbial ecosystem, but as complementary components for understanding the same biophysical system.

A quantitatively reconstructed resource environment also provides a way to interpret microbial phenotypes. Cellular traits that appear puzzling in isolation, such as the presence of particular transporters, patterns of global proteome regulation, or the strong expression of proteins with no apparent function under standard laboratory conditions, can instead be interpreted in the context of the resources encountered in the native environment. In a separate study, we apply this ecophysiological perspective to *Escherichia coli*, showing how the resource environment in its primary growth habitat, the ileocecal region, provides a physiological context for phenotypes that are difficult to understand without considering the environment in which they evolved [56].

The analysis here considers resource influxes into the large intestine as a whole and therefore establishes system-level supply and growth constraints rather than local resource concentrations or limiting conditions. Once resources enter the large intestine, their availability is dynamically reshaped by microbial uptake, secretion, and metabolic transformation, as well as by spatial variation in host secretion, mixing, and transit. Local resource environments and growth constraints can therefore differ substantially across the colon and between luminal and mucus-associated niches even when they are subject to the same overall influxes [57]. The resource influxes reconstructed here provide quantitative boundary conditions for extending this framework toward such spatially resolved descriptions of the gut environment.

The resource landscape reconstructed here also focuses on the healthy adult large intestine. An important next step will be to determine how this landscape changes with disease, inflammation, malnutrition, medication, and more extreme dietary conditions. Oxygen illustrates this particularly well. Increased epithelial oxygenation and oxygen availability during inflammation can favor facultative anaerobes such as *Escherichia coli* over obligate anaerobes [58, 59]. Luminal oxygen availability reflects a balance between diffusion from host tissue, oxygen consumption by the epithelium, and microbial uptake, all of which can change during inflammation. A mechanistic account of how inflammation reshapes the gut microbiome therefore requires quantifying how oxygen influx changes with the underlying physiological conditions rather than considering luminal oxygen concentration alone.

Ultimately, integrating quantitative resource influxes across physiological states with spatially resolved microbial transformations and microbiome-to-host molecular fluxes will be essential for mechanistically connecting diet and host physiology to microbial metabolism, community dynamics, and host exposure.

## Acknowledgments

I thank Markus Arnoldini, Théo Gervais, KC Huang, Richa Sharma, Orkun Soyer, Alfred Spormann, Andrew Verdegaal and all members of the Cremer and Spormann Labs for extensive discussion and critical feed-back on the manuscript. I also thank colleagues at the KITP workshop “Coarse-graining Microbial Ecology: from Genes to Physiological Strategies to Communities across Environments” for fruitful discussions. This research was funded in part by the National Institute of General Medical Sciences, National Institutes of Health, grant number 1R01GM149611. J.C. also acknowledges support via Bio-X Food@Stanford grant and a Terman Fellowship from Stanford University. This research was also supported by grant NSF PHY-2309135 and the Gordon and Betty Moore Foundation Grant No. 2919.02 to the Kavli Institute for Theoretical Physics (KITP).

## Notes

### Competing Interest Statement

The authors have declared no competing interest.

