## Supplementary Text for "Quantifying the Resource Landscape That Shapes the Human Gut Microbiome"

Jonas Cremer[1,2,*]

[1]Department of Biology, Stanford University, Stanford, CA, USA
[2]Bio-X, Stanford University, Stanford, CA, USA

**14 Summary of robust and assumption-dependent conclusions** **80**

**15 Supplementary Figures** **81**

**16 Supplementary Tables** **85**

**References** **99**

### 1 Overview of the quantitative resource reconstruction

In this Supplementary Information I provide the quantitative analysis underlying the resource influx estimates summarized in the main text. For each resource discussed in the main text I reconstruct the daily influx into the large intestine primarily from physiological measurements in humans, with a focus on healthy individuals. For each resource, I integrate available measurements to determine a representative influx range, assess its dependence on diet and host physiology where possible, and evaluate the confidence with which the influx can currently be estimated.

In the following, I present this analysis grouped by the type of resources entering, distinguishing dietary resources, host-derived compounds supplied from the ileum and across the gut epithelium, terminal electron acceptors, and ions. For each resource I also discuss potential dependencies on diet and assemble a mapping to reconstruct the entire resource landscape by estimating all fluxes together. I also convert the relevant molecular resource fluxes into elemental influxes, account for the potential microbial utilization of resources, and compare growth-accessible supply with microbial anabolic and catabolic requirements to determine potential growth limitations. Finally, I use the same quantitative supply–demand framework to investigate the ecological context of amino acid and corrinoid auxotrophies.

Throughout this analysis I focus on the gross chemical influx of a compound into the large intestine. These quantities are not identical to the flux of resources utilized by bacteria within the large intestine. For example some compounds entering the lumen are only partially accessible to bacteria or are taken up by the host again. I therefore first reconstruct physical resource influxes independently of their microbial fate and only subsequently comment on microbial accessibility.

### 2 Dietary resources entering from the ileum

As the first major category of resources available to the microbiome in the large intestine, I consider dietary resources entering from the ileum. The flux of different nutrient classes from food intake passing the terminal ileum and reaching the cecum can generally be estimated well. The passage of nutrients through the terminal ileum has been quantified systematically in human ileal effluent, often in controlled dietary intervention studies. Furthermore, the digestion and subsequent absorption of many macronutrients along the upper digestive tract and the small intestine is mechanistically well understood thanks to extensive research on human digestive physiology. Based on these findings, I can estimate mappings from diet to inflow into the ileum. Nevertheless, the reliability of different numbers varies considerably depending on study availability and the complexity of the biology at play.

Among the very large number of chemically distinct compounds present in the diet, I focus on those that are quantitatively substantial contributors to ileal outflow under typical diets and for which a tractable evidence base exists, at minimum dietary intake data combined with an estimable or measurable small-intestinal escape fraction. This selection is necessarily incomplete: numerous dietary compounds, including many individual micronutrients, phytochemicals, and food additives, are not treated separately here. A more complete reconstruction of the dietary resource landscape remains an important direction for future work; here, I provide a starting point by focusing on the most abundant and best-characterized dietary resources. In the following, I summarize the quantitative knowledge on the flux of carbohydrates, proteins, fats, polyphenols, and vitamins, with a focus on cobalamin (vitamin $B_{12}$), an instructive example of a low-abundance resource with important ecological relevance.

#### 2.1 Dietary carbohydrates

The amounts of dietary carbohydrates reaching the large intestine differ markedly from the amounts consumed in the diet because absorption along the small intestine selectively removes readily digestible substrates. Consequently, the carbon available to the gut microbiota is enriched in complex carbohydrates. Here I provide influx estimates for three major classes, resistant starch, dietary fibers, and soluble sugars, substantially extending our previous analysis of available studies to estimate the carbohydrates reaching the large intestine and its dependence on diet.

##### 2.1.1 Resistant starch

**Digestible and resistant starch** Starch is the major storage polysaccharide of plants and consists of glucose monomers linked by $\alpha$-glycosidic bonds in two principal components, the mostly linear polymer amylose ($\alpha$-1,4 linkages) and the highly branched polymer amylopectin ($\alpha$-1,4 and $\alpha$-1,6 linkages). Humans possess the enzymatic machinery to digest dietary starch efficiently through the combined action of salivary and pancreatic amylases together with brush-border enzymes in the small intestine. However, some fraction of consumed starch escapes digestion in the small intestine and reaches the large intestine. This fraction is operationally defined as resistant starch. Unlike fiber, resistant starch is not due to a distinct chemical composition – both digestible and resistant starch consist entirely of glucose. Instead, resistance to digestion arises from physical structure and accessibility. Different types of resistance have been well established through standardized measurements integrating biochemical assays and digestion in controlled dietary studies [1]. Following this work, starch is categorized into rapidly digestible starch (RDS), slowly digestible starch (SDS), and RS, with RS further divided by the mechanism of resistance: RS1, physical inaccessibility within intact plant cell walls or a food matrix (whole grains, seeds, legumes), destroyed by milling or thorough chewing; RS2, native starch granules with a resistant crystalline structure (raw potato, green banana, high-amylose maize starch), destroyed by cooking/gelatinization; RS3, retrograded starch formed when gelatinized starch is subsequently cooled and its amylose chains re-associate into a resistant structure;

RS4, chemically modified starch (cross-linked, esterified, or etherified), found only in industrially processed foods; and RS5, resistant amylose–lipid complexes. This classification means RS content is not a fixed property of a food the way fiber composition is; it depends strongly on both the composition of consumed food and how food is prepared. For a broad range of foods, the fraction of starch reaching the large intestine has been systematically measured for individual food items, varying between close to 0% for pure soluble starch and 96% for banana flour [2]. Cooking and cooling substantially alter this picture in both directions [3, 2, 4].

**Estimating resistant starch** To estimate RS fractions in entire meals, systematic measurements are available for foods representative of Western diets, including major staple foods such as boiled potato and white bread. Across these measurements, the fraction of dietary starch reaching the large intestine typically falls in the range of approximately 10–15% [1]. Building on these measurements, we previously adopted a working value of 13% of total dietary starch reaching the large intestine [5]. While this is a good starting point with the ranges well confirmed by different studies from the 1980s in the UK, it should be noted that recent RS intake could be substantially different due to changes in food composition and preparation habits.

**Variation in current Western diets** At the population level, Murphy, Douglass & Birkett [6] provide the most directly relevant estimate for extending the mapping above to more current diets, using the 1999–2002 NHANES dietary recall data set assessing the dietary behavior of the US population. Specifically, they built a food-specific RS concentration database from the published literature, linked it to each food reported, and estimated mean US resistant starch intake at roughly 3–8 g/day depending on demographic subgroup, with breads (∼21%), cooked cereals/pasta (∼19%), non-legume vegetables (∼19%), bananas/plantains (∼14%), and legumes (∼9%) as the largest contributing categories. Of course these numbers also come with uncertainty. Specifically, unlike many of the earlier studies from the UK, these numbers are not based on direct measurements of starch amounts escaping small-intestinal digestion for specific meals and specific meal preparations, missing for example some preparation-dependent RS3 contribution from how test meals were prepared.

**Modeling diet-dependent resistant starch delivery** I implement two models to estimate the amount of resistant starch (RS) entering the large intestine.

Simple mapping: I first estimate available starch as total carbohydrate intake minus simple sugars and fiber,

$$S_i = C_i - \mathrm{Fib}_i - U_i, \tag{1}$$

with $C_i$, $\mathrm{Fib}_i$, $U_i$ the NHANES-reported total carbohydrate, fiber, and sugar intake (`DR1ICARB`, `DR1IFIBE`, `DR1ISUGR`). Following Arnoldini et al. [5] I then assume a fixed population-wide fraction $\rho = 0.13$ of available starch escapes small-intestinal digestion as resistant starch,

$$\mathrm{RS}_i^{\mathrm{simple}} = \rho\, S_i. \tag{2}$$

**Detailed mapping.** As for fiber, I resolve each person's reported foods into base ingredients and assign each ingredient a starch content $\sigma_k$ (g starch / 100 g ingredient) from FNDDS, giving per-category starch intake $S_{i,c} = \sum_{k:c(k)=c} g_{i,k}\sigma_k/100$. Extending previous food-composition studies of RS content [6, 7], each category $c$ is assigned its own resistant-starch fraction of starch, $\rho_c$ – essentially zero for starch-poor categories (citrus, leafy/cruciferous vegetables, dairy, meat/egg/fish, fats/sugars/beverages), $\rho_c \approx 0.02\text{–}0.03$ for the starch-dominant staples refined wheat and rice, up to $\rho_c \approx 0.18$ for legumes – so that

$$\mathrm{RS}_i^{\mathrm{detailed}} = \sum_c S_{i,c}\,\rho_c. \tag{3}$$

I deliberately do not attempt to model RS3 formation from food preparation (cooking followed by cooling), as indicators of preparation state (e.g. distinguishing potato salad or reheated rice from freshly cooked equivalents) are largely absent from NHANES and in any case still leave substantial residual variation. My mapping should therefore be read as an approximate population-level estimate of variation in RS delivery, not a predictive model of specific individuals' intake, with a preparation-state-aware extension left for future work.

**Comparison of simple and detailed mapping.** The two RS mappings disagree substantially when using $\rho = 0.13$ as flat escape fraction, a value we calibrated previously for diets common in Britain of the 1970s [5]. This suggests that for modern diets, resistant starches are substantially less abundant than for the British reference diet. Indeed, the population-weighted average fraction implied by the detailed, food-category-resolved mapping on the present-day NHANES population is

$$\bar{\rho} = \frac{\sum_i \mathrm{RS}_i^{\mathrm{detailed}}}{\sum_i S_i} = 0.032, \tag{4}$$

roughly four-fold lower – consistent with the modern US diet's carbohydrate intake being dominated by refined wheat and rice, both low-RS categories, rather than the more RS-rich staples (legumes, less-processed grains) that contributed more to the reference diet. Applying $\rho = 0.13$ population-wide accordingly overestimates RS delivery on this population by a similar factor (median 14.68 g/day, 5th–95th percentile 4.74–32.47, versus the detailed mapping's 3.21 g/day, 0.80–9.37). Applying instead the population-weighted average fraction $\bar{\rho} = 0.032$ as a constant, $\mathrm{RS}_i^{\bar{\rho}} = \bar{\rho}\,S_i$, gives 3.57 g/day (1.15–7.90), in much closer agreement with the detailed mapping's 3.21 g/day (0.80–9.37) (Figure S1) – confirming that the two mappings are structurally consistent, and that the earlier discrepancy is attributable entirely to the flat fraction's calibration to a different population/diet, not to a difference in modeling approach. I adopt $J_{\mathrm{RS},i}^{\mathrm{LI}} = \mathrm{RS}_i^{\mathrm{detailed}}$ as the resistant-starch influx carried forward into the elemental-conversion step (Section 9).

**Summary and confidence classification** Overall, starch entering the large intestine can be estimated with good confidence. The digestion of starch is well understood and the total amount of resistant starch reaching the large intestine is well constrained, anchored by a consistent body of measurements from more than 20 years ago, spanning specific staple foods, dependencies on meal preparation, and population-level estimates. For the US population, my detailed, food-category-resolved mapping gives a range between approximately 0.8 and 9.4 g/day (5th–95th percentile), with a median of around 3.2 g/day – broadly consistent with the independent population-level estimate of Murphy et al. [6] (mean 3–8 g/day across demographic subgroups, using an earlier NHANES cycle and a different food-to-RS mapping). The principal uncertainty does not arise from a lack of digestion understanding and measurements, but from the complex dependence of starch digestibility on food composition, processing, cooking, and retrogradation. Future work systematically relating dietary intake and food preparation to resistant starch delivery, together with improved quantification of the individual RS classes, would substantially improve quantitative predictions of the carbohydrate amount entering the large intestine.

##### 2.1.2 Dietary fiber

**Definition and overall flux** Dietary fibers comprise a diverse group of largely non-digestible polysaccharides, including cellulose, hemicellulose, pectins, arabinoxylans, inulin, resistant oligosaccharides, and $\beta$-glucans. Humans possess little enzymatic capacity to degrade these polymers, resulting in negligible digestion and absorption within the small intestine. Consequently, and in contrast to starch, the vast majority of consumed fiber reaches the cecum. This is confirmed by direct ileostomy measurements, which find non-starch polysaccharides such as the water-soluble beta-glucan in oats almost completely recovered in ileal effluent regardless of food source [3]. This makes estimating the amount of fiber reaching the large intestine comparatively easy, as it is essentially set by the amount of fiber consumed [5], which itself can be estimated well as the fiber content of food, routinely measured using a standardized enzymatic-gravimetric method [8]. While measurement specifics changed slightly over time to better capture RS4-type fibers and oligosaccharides, food surveys reported comparable ranges of fiber consumption. For example, for the United States population in the 1980s, a fiber consumption range between 2 and 27 g per day has been reported, with an average fiber consumption of approximately 12 g/day [9]. A similar analysis with the recent NHANES dataset from 2017/18 shows that intake varies between 4 and 37 g/day, with a median of 14 g/day [5].

**Different fiber types** Beyond total fiber content, the relative proportions of individual fiber classes such as cellulose, hemicellulose, pectin, arabinoxylans, lignin, $\beta$-glucan, and fructan-type oligosaccharides like inulin are important to differentiate, since different fiber types are utilized with different efficiency by the microbiome (Section 9.5), with strong variation between different bacterial species. The split between soluble and insoluble fiber, and the compositional split between different fiber types, have been determined for a substantial set of individual food ingredients, directly measuring, for example, cellulose, hemicellulose, pectin, and lignin content [10, 11]. Summary statistics for particular food groups are available [12], such as the characteristic cellulose content of legumes or the pectin content of fruit. While the coverage of food ingredients is not complete, it often covers a substantial fraction of consumed fiber. In my quantitative mapping relating diet composition to fiber intake below, I build on these available data to approximate the amounts of different fiber types consumed and their variation in the US population.

**Modeling fiber reaching the large intestine** Given the limited digestion and absorption of fiber in the small intestine, I model total fiber delivery to the large intestine directly as the fiber content of the diet, using NHANES-reported fiber intake. For person $i$, a simple mapping is

$$F_i^{\text{simple}} = \mathrm{Fib}_i, \tag{5}$$

where $\mathrm{Fib}_i$ is NHANES-reported total dietary fiber intake (variable DR1IFIBE), summed across all foods and beverages reported in that person's 24-hour recall.

To resolve this total into individual fiber types, I consider a finer, food-level mapping. I start with every food item $f$ and its consumed mass $m_{i,f}$ (g) that person $i$ reported eating in the NHANES dietary recall. Each food is resolved recursively into its constituent base ingredients using the USDA FNDDS 2017–2018 recipe database, yielding a set of resolution weights $w_{f,k}$ (the mass fraction of food $f$ attributable to base ingredient $k$, $\sum_k w_{f,k} = 1$), so that the total mass of ingredient $k$ consumed by person $i$ is

$$g_{i,k} = \sum_f m_{i,f}\, w_{f,k}. \tag{6}$$

Each ingredient $k$ carries an FNDDS-reported fiber content $\phi_k$ (g fiber / 100 g ingredient), giving person $i$'s total fiber intake from ingredient $k$ as $g_{i,k}\phi_k/100$.

I then assign every ingredient to one of ∼20 broad food-type categories $c(k)$ (e.g. whole grains, legumes, citrus fruit, root vegetables) by keyword matching on the ingredient description, and aggregate to the category level:

$$F_{i,c} = \sum_{k:\, c(k)=c} g_{i,k}\, \phi_k/100. \tag{7}$$

Each category $c$ is assigned a default fiber-type composition $\theta_{c,t}$, i.e. the fraction of category $c$'s fiber mass present as fiber type $t \in \{$cellulose, hemicellulose/arabinoxylan, $\beta$-glucan, pectin, inulin/FOS, GOS, lignin, other$\}$, with $\sum_t \theta_{c,t} = 1$. For grain, legume, and fruit categories, these defaults are grounded in group-level fiber-composition statistics from Marlett [12] (e.g. soluble fiber ∼23% of total fiber in refined grains, cellulose ∼50% of fiber in legumes, pectin 15–30% of fiber in fruit; only the paper's reported summary statistics were used, as its full per-food table was not accessible). The remaining categories (roots, most vegetables, nuts/seeds, dairy, and the non-$\beta$-glucan portion of oats) are instead approximated from general plant cell-wall composition patterns (e.g. leafy-vegetable fiber skewing more cellulose-dominant than fruit fiber), without a specific per-category literature source. Rather than relying on these category-level defaults uniformly, I override them wherever an ingredient is directly matched to published, ingredient-level fiber-fractionation data [10, 11], replacing $\theta_{c(k),t}$ with the ingredient's own measured composition $\theta_{k,t}$ for that ingredient's contribution specifically. Marlett & Cheung's four-way cellulose/hemicellulose/pectin/lignin split takes precedence where both sources cover the same ingredient, with the remaining four fiber types ($\beta$-glucan, inulin/FOS, GOS, other) then filled in proportionally to their category-default shares so that $\sum_t \theta_{k,t} = 1$ is preserved; Li et al.'s soluble/insoluble split, which rescales all eight category-default fractions by a common soluble or insoluble factor, is used otherwise. The detailed-model total for fiber type $t$ is then

$$F^{\text{detailed}}_{i,t} = \sum_c \Bigg[ \underbrace{F^{\text{unmatched}}_{i,c}\, \theta_{c,t}}_{\text{category default}} + \underbrace{\textstyle\sum_{k:\, c(k)=c,\, k\in\mathcal{M}} g_{i,k}\phi_k/100\; \theta_{k,t}}_{\text{directly measured, matched ingredients } \mathcal{M}} \Bigg], \tag{8}$$

where $F^{\text{unmatched}}_{i,c}$ is category $c$'s fiber mass excluding the directly-matched ingredient set $\mathcal{M}$. By construction, summing across fiber types recovers the same person-level total as the food-level resolution, $\sum_t F^{\text{detailed}}_{i,t} = \sum_c F_{i,c}$, independent of the type-split. With this two-step approach, roughly half of population fiber mass is currently grounded in directly measured fiber-type composition (matched ingredients), with the remainder resting on literature-informed category defaults; the dietarily most important ingredients are disproportionately well covered, since matching prioritized high-consumption items. As such it should be noted that the distribution of fiber type consumptions are only first approximations to describe the variability across individuals and diets.

**Comparison of simple and detailed mapping.** The two mappings answer different questions – the simple mapping gives a population-wide total, while the detailed mapping resolves that total into fiber types and their food origins. However, since both are anchored to the same underlying NHANES fiber-intake data, the population-level total they imply should agree closely, and doing so is a useful consistency check on the food-level resolution pipeline itself. This is indeed the case: median total fiber reaching the colon is 14.50 g/day (5th–95th percentile 3.90–38.10) under the simple mapping versus 14.36 g/day (3.83–37.63) under the detailed mapping ($n = 4741$) with a near-total overlap in the population distributions (Figure S1). This agreement confirms that the FNDDS ingredient-level fiber values, once resolved through the recipe hierarchy and aggregated per person, reproduce NHANES's own reported fiber totals almost exactly. I adopt $J^{\mathrm{LI}}_{\text{fiber},i} = F_i^{\text{simple}} = \mathrm{Fib}_i$ as the total fiber influx carried forward into the elemental-conversion step (Section 9), since the two mappings agree closely on total mass. The model applies a single, population-wide utilization fraction to this total (50%, Table S3), reflecting the fact that mixed fiber types ferment very differently; the detailed, per-type breakdown $F_{i,t}^{\text{detailed}}$ is not itself run through a per-individual accessibility-weighted average in the model, but its type-specific fermentability values (also listed in Table S3) informed the choice of this single working value, since individual fiber types differ substantially in microbial accessibility.

**Summary and confidence classification** Overall, the amount of fiber reaching the large intestine can be estimated with good confidence, given comparably simple digestion physiology and systematic food-composition data, including standardized fiber measurements. For the US population, a broad range between 4 and 37 g/day is observed, with a median of around 14 g/day. The relative amounts of major fiber types reaching the large intestine can also be estimated, although with higher uncertainty, as the direct quantification of fiber types in different food ingredients remains less systematic.

##### 2.1.3 Free glucose and other soluble sugars

**Challenging influx estimations** Part of the consumed carbohydrates also reaches the human large intestine as free mono- and oligosaccharides. Quantifying this influx is, however, more challenging, because concentrations are low and can easily be shifted either by bacterial activity in the collected sample or by sample treatments that release additional sugars from polysaccharides before measurement. Accordingly, the values reported by different studies vary substantially, and each carries its own specific risk of bias. For example, Stephen et al. [13], via multilumen ileal intubation applied to single test meals, found mean glucose rising from 186.9 mg (20 g starch meal, $\sim$0.9% of ingested starch) to 1261.9 mg (60 g starch meal, $\sim$2.1%), with total unabsorbed carbohydrate (glucose + amyloglucosidase-releasable material) averaging 6–9% of ingested starch across meal sizes. However, this value could be a substantial overestimate, given the heat and acid pretreatment used, which could itself release additional sugar from oligosaccharides before measurement. In contrast, Berghouse et al. [14], in ileostomy subjects on refined vs. unrefined carbohydrate diets, measured combined free glucose + oligosaccharide at $82.1 \pm 25.8$ vs. $169.0 \pm 40.8$ mg over a 9 h postprandial period – approximately 0.18% and 0.36% of the starch dose, respectively, with free glucose consistently about one-third of the oligosaccharide fraction. Here, bacterial growth in the ileostomy effluent during the collection period could substantially shape this value, most plausibly biasing it toward an underestimate of the true flux. Flourié et al. [15], via a similar ileal intubation approach to Stephen et al., but without a heat pretreatment step, found unabsorbed glucose rose from $1.1 \pm 0.3$ to $2.7 \pm 0.8$ g/24h (approximately 1.1% and 0.9% of ingested starch, respectively) as dietary starch intake increased from 100 to 300 g/day. More recently, van Trijp et al. [16] applied high-performance anion-exchange chromatography (HPAEC-PAD) directly to samples from the distal ileum and proximal colon. They detected a glucose and galactose signal but could not confidently attribute it to genuine luminal free sugar, noting it could instead reflect breakdown of galacto-oligosaccharides during sample handling, or interference from host-derived mucus saccharides. This emphasizes that fundamental measurement ambiguities persist even with modern instrumentation.

**Strong dependence on diet** Irrespective of the challenges in measuring absolute sugar influx numbers, all studies consistently report free sugars in ileal effluent to increase strongly with the amount of carbohydrates consumed. This suggests that soluble sugars reaching the large intestine come mostly from the partial digestion of complex carbohydrates, not from free sugars in the diet. This view is consistent with measured intestinal uptake capacities that comfortably exceed typical intake rates: glucose uptake rates as high as 550 mmol/h have been measured in humans [17, 18], and hydrolysis and uptake of sucrose, the main dietary sugar, proceeds at around 100 mmol/h [19] – both far in excess of typical dietary sugar consumption (e.g. $\sim$59 g or 172 mmol sucrose/day for a British reference diet [5]). Fructose is a notable exception to this pattern, as its intestinal transporters have lower capacity and saturate more readily, so free dietary fructose taken up in high quantities is comparatively more likely to escape complete small-intestinal absorption [20].

**Mapping diet to free-sugar flux** To approximate the diet-dependent influx of sugars into the large intestine, I use a simple two-parameter model for the flux of free glucose, maltose, and short oligosaccharides derived from starch reaching the colon, reusing the available-starch quantity $S_i$ already defined for resistant starch (Section 2.1.1, Eq. 2).

$$J^{\mathrm{LI}}_{\text{free-sugar},i} \approx S_i \times f_{\text{escape}} \times f_{\text{monomeric}} \tag{9}$$

Here, $S_i$ is person $i$'s available dietary starch (g/day; total carbohydrate intake minus fiber and sugar, as defined in Section 2.1.1), $f_{\text{escape}}$ is the fraction of ingested starch escaping small-intestinal absorption in any form (glucose, maltose, oligosaccharide, or intact resistant starch), and $f_{\text{monomeric}}$ is the fraction of that escaped material present as free glucose, maltose, or short oligosaccharides rather than higher-degree-of-polymerization resistant starch. The literature reviewed above supports a food-dependent escape fraction ($f_{\text{escape}}$) of roughly 1%. Furthermore, the only study offering a within-food split of escaped starch by chemical form is Englyst & Cummings [3], who found oat starch escaping digestion appeared as $<$0.6% intact starch plus a further 1.2% as dextrins and maltose – i.e., roughly two-thirds of the escaped material in short-chain/monomeric form. I therefore adopt $f_{\text{monomeric}} \approx 0.6$–$0.7$ as a provisional default, explicitly flagging that this needs additional data to generalize beyond a single cereal food. Applying Equation 9 with the suggested central values ($f_{\text{escape}} = 0.01$, $f_{\text{monomeric}} = 0.65$) to a typical starch intake of 150–250 g/day yields an estimated free glucose/maltose/short-oligosaccharide flux of roughly 1.0–1.6 g/day – in good agreement with the 1–3 g/day range directly measured by Stephen et al. and Flourié et al.

**Beyond starch: fructose and other free sugars** The mapping above captures only free sugar arising from starch digestion (glucose, maltose, and short oligosaccharides). Free fructose, primarily from dietary sucrose, high-fructose corn syrup, and fruit, is a separate and potentially non-trivial contribution that this equation does not include. Unlike glucose, fructose crosses the apical membrane via GLUT5, a facilitative transporter with a genuinely saturable capacity: dose-response studies using hydrogen breath testing found healthy adults absorb single doses of 15 g fructose essentially completely, show incipient malabsorption in a minority (10%) at 25 g, and malabsorption in the majority (80%) at 50 g [21]. This capacity is substantially higher when fructose is co-ingested with glucose, as it almost always is in the diet (sucrose is 50:50 glucose:fructose; high-fructose corn syrup is approximately 45:55) – the same study noted that a 50 g fructose dose reliably causing malabsorption in isolation often does not do so when taken with normal mixed food. US adult dietary fructose intake has been estimated at roughly 37–55 g/day across recent decades, with higher-consuming groups reaching 85–100 g/day [22]. Spread across typical eating occasions, this corresponds to per-meal fructose exposure mostly at or below the single-dose malabsorption threshold, suggesting a modest average escape fraction (order of a few percent of total intake) for most individuals under typical, food-matrix-embedded consumption – corresponding to a rough estimate of 1–3 g/day reaching the colon as free fructose, with substantial individual variation (higher in fructose malabsorption or concentrated high-dose consumption, e.g. sugar-sweetened beverages).

Other free monosaccharides – galactose, xylose, and arabinose – are expected to contribute comparatively little, for distinct reasons. Galactose is transported with similar efficiency to glucose once liberated from lactose by lactase; where lactase activity is limiting, the sugar reaching the colon is predominantly intact lactose rather than free galactose. Xylose and arabinose are structural components of hemicellulose that humans lack the enzymatic machinery to release in the small intestine at all, so these pentoses should be essentially absent as free monosaccharides in ileal effluent, appearing only after bacterial breakdown of the intact fiber polysaccharide once it has already reached the colon.

Combining these considerations, I estimate total free sugars (glucose, maltose, short oligosaccharides, fructose, and trace contributions from other monosaccharides) reaching the large intestine at approximately 1–5 g/day. The quantitative resource-limitation model (Section 8) uses this combined estimate (2.5 g/day, the central value of the 1–5 g/day range) as this resource's input, capturing overall soluble-sugar delivery rather than only the narrower, starch-derived component (Eq. 9, 1.0–1.6 g/day).

**Summary and confidence classification** Overall, the amount of soluble sugars reaching the large intestine can be estimated with low confidence, now understood to span an estimated 1–5 g/day once both the starch-derived (glucose, maltose, short oligosaccharides) and non-starch (chiefly fructose) contributions are considered together. Amounts are well established to be low compared to resistant starch and fiber influx, in line with strong absorption of sugars along the small intestine. It is also well supported that the starch-derived component stems largely from the digestion of complex carbohydrates, while the fructose component instead reflects the saturable capacity of its dedicated transporter relative to typical intake. However, exact influx numbers remain challenging to measure and come with high uncertainty, and the fructose contribution specifically is inferred from absorption-capacity and intake data rather than direct ileal measurement. With the exception of glucose, the types of soluble sugars entering the large intestine, such as the abundance of xylose, arabinose, fructose, or galactose, are also less well characterized by direct measurement. Note that these considerations highlight the need for modern metabolic measurements, systematically quantifying the concentration of different free sugar types in ileal effluent. Modern targeted methods such as high-resolution anion-exchange chromatography and sugar-specific enzymatic assays have already been applied to human ileal and ileostomy samples [16, 23]. Their systematic use in controlled dietary studies now provides a unique opportunity to generate such data with much improved sugar-specific resolution.

#### 2.2 Dietary proteins, peptides, and amino acids

**Protein digestion and absorption in the small intestine** During their passage through the stomach and small intestine, dietary proteins are efficiently hydrolyzed by host-derived proteases into peptides and free amino acids, the majority of which are subsequently absorbed by the small intestine. However, a small but substantial fraction of dietary protein still reaches the large intestine [24].

The amount of total amino acids escaping small-intestinal absorption is comparatively well characterized, reflecting extensive work in human nutrition aimed at quantifying protein digestibility and comparing the nutritional quality of different protein sources. A broad range of literature has quantified amino acid digestibility and host absorption, incorporating human ileostomy studies, sampling through naso-ileal tubing, and isotope labeling [25, 26]. Rigorous measurements in growing pigs, including direct comparisons with humans, have established the pig as a suitable model for human protein digestion in the small intestine. Standardized digestibility data are now available for a broad range of foods and the impact of food preparation has been studied [27, 28]. I here utilize these measurements to systematically evaluate the amount of dietary proteins escaping absorption along the small intestine and reaching the large intestine.

**True ileal digestibility** These studies determine protein digestibility by measuring for different diets and protein types the loss of total amino acids (sum of free amino acids, peptides, and proteins) in ileum fluids. Comparing this loss to the amount of amino acids ingested provides a digestion efficiency. Since the host also secretes proteins into the intestine in form of mucin, digestion enzymes or other amino acid containing substrates, the comparison corrects for the endogenous amino acid secretion by also quantifying the efflux of amino acids in a protein-absent diet. Subtracting this endogenous efflux from that observed when protein-containing diets are consumed provides the basis for estimating *true ileal digestibility*, i.e. the fraction of dietary amino acids absorbed by the host and thus not reaching the large intestine [29, 25]. The true ileal digestibility ranges between 85–98% varying strongly with food type. Systematic datasets now exist for a wide range of individual food ingredients, with animal-derived proteins generally exhibiting higher digestibility than plant-derived proteins [28, 27]. Similar to resistant starch, food processing can further modify these values. For example, for a diet rich in soya bean proteins amino acid and total nitrogen efflux from the ileum were almost twice as high for whole compared to pureed soya beans [30]. However, this preparation dependence is not systematically explored across food types.

**Distinguishing between free amino acids, peptides, and proteins** There is also some information on whether amino acids enter as free amino acids, peptides. Specifically, the same systematic characterization of ileal effluent mentioned earlier [30], also quantified intact proteins, peptides, free amino acids, ammonia, and urea under controlled dietary interventions. Their measurements demonstrate that proteinaceous material accounts for the overwhelming majority of ileal nitrogen, whereas free amino acids contribute only a comparatively small fraction (as do ammonia and urea, see Section 3.4). Independent measurements similarly conclude that approximately 86% of terminal ileal nitrogen is proteinaceous [31, 32].

**Mapping diet to amino acid flux** To estimate the amount of amino acids reaching the large intestine, I implemented two mappings with different levels of detail.

**Simple mapping.** I start with total dietary protein intake $\mathrm{Prot}_i$ (`DR1IPROT`) and apply a single, population-wide true ileal digestibility (TID) value to estimate the fraction escaping small-intestinal absorption:

$$\mathsf{AA}_i^{\mathsf{simple}} = \left(1 - \frac{\mathsf{TID}}{100}\right) \mathrm{Prot}_i. \tag{10}$$

I assume TID $= 91\%$, the average of two literature-derived brackets [28, 27]: TID $= 95\%$, representing an animal-protein-dominated diet (meat, dairy, fish, eggs), and TID $= 87\%$, representing a plant-protein-dominated diet (pulses, grains, wheat, soy). Applying TID $= 91\%$ to the NHANES 2017–2018 adult population gives a dietary amino-acid flux reaching the large intestine of 6.44 g/day (5th–95th percentile 2.37–14.17, $n = 4741$).

**Detailed mapping.** I resolve each person's reported foods into base ingredients as above, assign each ingredient a protein content $\pi_k$ (g protein / 100 g ingredient) from FNDDS, and classify each ingredient into one of the food-source categories $c$ for which Hodgkinson et al. [28] report a directly measured category-median TID, $\mathsf{TID}_c$ (a 97-food human ileostomy/pig-cannulation study). Per-category protein intake is $\mathrm{Prot}_{i,c} = \sum_{k:c(k)=c} g_{i,k}\pi_k/100$, and total amino-acid flux is

$$\mathsf{AA}_i^{\mathsf{detailed}} = \sum_c \mathrm{Prot}_{i,c} \left(1 - \frac{\mathsf{TID}_c}{100}\right). \tag{11}$$

This lets each person's own reported food-source mix determine their effective digestibility, rather than a single population-wide animal/plant bracket. Applied to the same population, this gives a lower flux of 4.24 g/day (5th–95th percentile 1.52–9.32), corresponding to a population-mean effective TID of

$$\overline{\mathsf{TID}} = 100 \left(1 - \frac{\sum_i \mathsf{AA}_i^{\mathsf{detailed}}}{\sum_i \mathrm{Prot}_i}\right) = 93.7\%, \tag{12}$$

higher than the simple mapping's unweighted 91% bracket midpoint – i.e. the average US diet's food-source mix is somewhat more digestible than the naive animal/plant midpoint would suggest.

**Comparison of simple and detailed mapping.** Because the detailed mapping rests on directly measured, food-category-specific ileal digestibility rather than a literature-averaged bracket, I treat it as the better-grounded estimate and adopt it as my primary working range for the remainder of this work. To test whether the two approaches are otherwise structurally consistent, I recomputed the simple mapping using the detailed mapping's own population-mean digestibility, $\overline{\mathsf{TID}} = 93.7\%$, in place of the ad hoc 91% bracket midpoint:

$$\mathsf{AA}_i^{\overline{\mathsf{TID}}} = \left(1 - \frac{93.7}{100}\right) \mathrm{Prot}_i. \tag{13}$$

This gives 4.51 g/day (5th–95th percentile 1.66–9.93), closely matching the detailed mapping's 4.24 g/day

(1.52–9.32) – a $\sim$6% difference at the median, down from the $\sim$50% difference between the original 91%-bracket simple mapping and the detailed mapping (Figure S1). I adopt $J^{\mathrm{LI}}_{\mathrm{AA},i} = \mathrm{AA}^{\mathrm{detailed}}_i$ as the amino-acid influx carried forward into the elemental-conversion step (Section 9), per the better-grounded status established above.

**Summary and confidence classification** Overall, the total amount of amino acids (free amino acids, peptides, and proteins) reaching the large intestine can be estimated with high confidence, thanks to systematic studies of protein digestibility conducted for human nutrition, which can be utilized for this purpose. Differences in digestibility between plant and animal proteins are well established. Only the dependence on food processing has been studied less systematically, and ideally warrants further attention in future studies. Given the strong variation in consumed protein amounts and protein types, the amount of dietary amino acids reaching the large intestine varies widely across the population: using my detailed, food-category-specific digestibility mapping, I estimate a 5th–95th percentile range of 1.5–9.3 g/day, with a median of around 4.2 g/day, for the US population.

#### 2.3 Dietary lipids

**Lipid influx and turnover** Dietary lipids including triglycerides (commonly referred to as 'fat') phospholipids, and sterols constitute the third major macronutrient class, alongside proteins and carbohydrates. While consumed in substantial daily amounts, lipids are generally well digested, and their absorption efficiency in the small intestine is high. Accordingly, only a comparatively small amount of dietary lipids reaches the large intestine. Controlled ileostomy studies across a range of high- and low-fat diets show that the amount of lipids reaching the large intestine scales with dietary fat intake with the amount of lipid leaving the small intestine typically corresponding to 3–5% of dietary fat intake [33, 34]. However, substantial deviations from this range seem possible and 12% has been reported for high-fiber diets [33].

Independent measurements of fat excretion in feces are consistent with these numbers, with about 2–4% of dietary fatty acids recovered in feces [35, 36]. Based on this consistency, fecal fat measurements have even become a standard clinical assay for impaired fat digestion and absorption [37].

**Specific lipids and fatty acids** What is much less well characterized is the molecular composition of the lipids and fatty acids entering the large intestine. I am, for example, not aware of direct measurements of the relative abundance of phospholipids, triglycerides, or other lipid classes in ileal effluent, nor of the concentrations of free long-chain fatty acids and glycerol. In contrast, some of these measurements are available for fecal samples. Fatty acids detected in feces predominantly comprise C12–C18 species, broadly reflecting the fatty acid composition of the diet. A notable exception is oleic acid, which is recovered at lower abundance than saturated long-chain fatty acids. This could reflect preferential microbial utilization during colonic transit [35], although other explanations, including more efficient host absorption, remain possible. More detailed analyses of fecal lipid classes further suggest that triglycerides account for only a minor fraction of fecal lipids in healthy individuals [38], indicating efficient digestion and absorption of dietary triglycerides in the small intestine. However, the relative abundance of phospholipids remains poorly quantified. Modern lipidomic studies identify numerous phospholipid species, together with free fatty acids, sphingolipids, and bile lipids in feces [39], while recent capsule-based sampling studies similarly reveal a diverse lipid composition throughout the intestinal tract [40].

**Limited microbial utilization of fatty acids** The substantial recovery of long-chain fatty acids in feces, broadly matching their influx from the ileum, suggests that gut bacteria do not extensively catabolize these compounds during colonic transit. This is consistent with the requirement for external electron acceptors during fatty-acid oxidation, which are scarce in the largely anoxic gut lumen [41] (see also Section 7). Anabolic incorporation also appears unlikely to account for most of this lipid pool: from simultaneous measurements of fecal fat content and fecal dry mass, I estimate that bacterial fatty acids could account for at most approximately 15% of the fatty acids recovered in feces.

Notably, this limited anabolic and catabolic use of fatty acids does not exclude a potentially substantial lipid digestion by bacteria into fatty acids. Many gut bacteria encode lipases and phospholipases capable of hydrolyzing lipids into fatty acids and glycerol. Further studies are needed to better understand the extent to which bacterial lipid hydrolysis occurs within the human large intestine and how bacterial lipid hydrolysis compares to host-mediated lipid digestion in the small intestine.

**Modeling lipid delivery to the large intestine** Given the evidence above that lipid recovery in the large intestine scales approximately linearly with dietary fat intake, I model lipid delivery for person $i$ as a fixed fraction of consumed dietary fat,

$$L_i = \eta \, T_i, \tag{14}$$

where $T_i$ is NHANES-reported total dietary fat intake (DR1ITFAT) and $\eta = 0.04$ is the recovery fraction, chosen within the 3–5% range established by the controlled ileostomy studies discussed above. I do not distinguish between lipid classes (triglycerides, phospholipids, sterols) or between intact lipid and free fatty acid/glycerol forms reaching the large intestine, given the lack of ileal-effluent-level compositional data noted above. Applied to the NHANES 2017–2018 adult population (dietary fat intake 5th–95th percentile 22.8–176.9 g/day, $n = 4741$), this gives a lipid flux reaching the large intestine of 0.9–7.1 g/day (5th–95th percentile), with a median of 3.1 g/day. I adopt $J^{\mathrm{LI}}_{\mathrm{lipid},i} = L_i$ as the lipid influx carried forward into the elemental-conversion step (Section 9).

**Summary and confidence classification** Overall, I can estimate the total influx of lipids into the large intestine with moderate confidence. The physiology of fat digestion and absorption is well understood. Furthermore, several independent studies report broadly consistent absorption values, with about 4% of consumed lipids reaching the large intestine, corresponding to a 5th–95th percentile range of 0.9–7.1 g lipids entering the large intestine per day for the NHANES cohort (median 3.1 g/day). However, how absorption depends on fat type and food preparation is less well characterized. Furthermore, the molecular composition of lipids entering the large intestine remains comparatively poorly characterized. Modern lipidomic studies identify a wide diversity of lipid species but rarely report absolute abundances, whereas classical physiological studies quantify total lipid fluxes without resolving lipid classes. There is therefore a need for modern lipidomic studies systematically quantifying the molecular composition of cecal and ileal contents.

#### 2.4 Nucleic acids and nucleotides

Dietary nucleic acids constitute an additional source of carbon, nitrogen, and phosphorus that could potentially become available to the large-intestinal microbiota. Nucleic acids are present in essentially all cellular foods, with particularly high concentrations in foods rich in cellular material such as organ meats, seafood, and yeast. Estimates of dietary nucleic-acid intake are generally on the order of approximately 0.1-1 g/day [42], substantially smaller by mass than the dietary protein and carbohydrate fluxes considered above.

**Digestion and absorption** Dietary DNA and RNA are extensively digested in the gastrointestinal tract. Pancreatic nucleases and intestinal phosphatases and nucleotidases convert nucleic acids successively into nucleotides, nucleosides, and nucleobases. Nucleosides represent important products of nucleoprotein digestion and can be absorbed by intestinal epithelial cells through concentrative and equilibrative nucleoside transporters [43]. The quantitative physiology of nucleobase absorption in humans is less well resolved, and important differences exist between humans and commonly used experimental animals [44].

These observations indicate that a substantial fraction of dietary nucleic-acid material is processed and absorbed before reaching the large intestine. However, I am not aware of direct human measurements quantifying total nucleic acids, nucleotides, nucleosides, and nucleobases leaving the terminal ileum, nor of sufficiently well-constrained absorption efficiencies that would allow this flux to be reconstructed reliably from dietary intake.

**Additional endogenous sources** Dietary intake is furthermore not necessarily the only source of nucleic-acid-derived material and purines entering the large intestine. Sloughed epithelial cells, mucus-associated cellular material, and other endogenous secretions contain DNA and RNA and could contribute to the luminal nucleotide pool but the extent to which this material is degraded or absorbed before becoming available to large-intestinal microorganisms is not quantitatively established.

In addition to nucleic acids released with cellular material, soluble purine metabolites can be supplied directly by the host. Uric acid, the major end product of human purine degradation, is excreted not only through the kidney but also into the intestinal tract. Recent work further demonstrated that gut bacteria from several phyla can anaerobically catabolize purines, including uric acid, and that this microbial activity can alter intestinal and systemic purine concentrations [45, 46]. Thus, the resource pool available for microbial purine metabolism cannot be inferred from dietary nucleic-acid intake alone, but reflects dietary, cellular, and host-derived purine fluxes.

**Contribution to overall influx** At a dietary intake on the order of 0.1-1 g nucleic acids/day, nucleic acids could in principle contribute a measurable amount of nitrogen and phosphorus to the intestinal resource landscape. However, because the fraction escaping small-intestinal digestion and absorption is unknown, I do not include dietary nucleic acids as a quantitatively resolved contribution to the elemental growth analysis. This omission is unlikely to dominate the total nitrogen influx given the substantially larger protein-, amino-acid-, ammonium-, and urea-derived nitrogen fluxes, but could be more relevant for phosphorus because of the smaller total phosphorus requirement of microbial biomass.

**Summary and confidence classification** The digestion pathways and intestinal transport systems for nucleic-acid-derived metabolites are reasonably well characterized, and recent work establishes that purines reaching the intestine can constitute physiologically relevant microbial substrates [45]. However, their quantitative influx into the human large intestine remains poorly constrained. In particular, I found no direct human ileal mass-balance measurements that would allow dietary nucleic acids, nucleotides, nucleosides, and nucleobases to be quantified, and endogenous contributions from shed cells and host-derived soluble purines introduce additional unresolved fluxes. I therefore classify the influx of nucleic acids and their breakdown products into the large intestine as **unknown**. This resource is accordingly not currently included in the quantitative $J_{r,i}^{\mathrm{LI}}$ reconstruction (Section 8).

#### 2.5 Cobalamin, other vitamins and trace elements

Beyond carbohydrates, proteins, and lipids, vitamins are another important class of dietary components shaping gut microbiome dynamics.

##### 2.5.1 Cobalamin

Here, I focus on cobalamin (vitamin $B_{12}$) and other corrinoids, which provide a well-established example of micronutrients shaping microbial interactions in the gut. Cobamide-dependent metabolism is widespread, whereas complete de novo biosynthesis is restricted to a subset of bacteria, creating extensive potential for uptake, salvaging, remodeling, and cross-feeding [47, 48, 49, 50]. Experimental studies further show that corrinoid identity and transporter repertoire can affect bacterial fitness and that complex gut communities can sustain $B_{12}$ production and sharing in vitro [48, 51]. In contrast, the physiological influx of cobalamin and other corrinoids into the human large intestine, and how this external supply compares with microbial production and demand, remain poorly quantified. Here I estimate the influx of canonical cobalamin and compare its scale with fecal corrinoid output.

**Terminology** I distinguish in the following between three related classes of compounds. *Corrinoids* denote the complete family of cobalt-containing corrin-ring compounds. *Cobamides* are corrinoids containing the complete nucleotide loop and constitute the biologically active enzyme cofactors used by microorganisms. *Cobalamin* (vitamin $B_{12}$) refers specifically to the dimethylbenzimidazole-containing cobamide required by human metabolism. While cobalamin is the physiologically relevant molecule for the host, the broader corrinoid pool is likely the ecologically relevant resource for gut microbial communities [52].

**Mapping cobalamin influx into the large intestine** Unlike most other nutrients considered in this work, I estimate cobalamin influx into the large intestine not from ileal digestibility or fecal recovery data, but from a simple whole-body mass-balance argument. Cobalamin is efficiently absorbed in the terminal ileum and secreted into the bile, so both dietary intake and biliary secretion feed the same luminal pool that ultimately determines colonic influx [53]. At first sight, this recycling complicates the estimate, since intestinal influx depends on both uptake and recycling in the body. However, the balance simplifies over sufficiently long timescales. Human body stores of cobalamin change slowly, and urinary losses are small relative to gastrointestinal turnover [53]. Under approximate steady state, net gastrointestinal and urinary losses of body cobalamin must be balanced by absorbed dietary cobalamin. Because urinary losses are small, combining this whole-body balance with incomplete absorption of dietary cobalamin leads to the first-order result

$$J_{\text{colon}} \approx J_{\text{diet}}.$$

With only limited $B_{12}$ absorption along the large intestine [54], the average dietary intake alone provides a reasonable first-order estimate of average cobalamin influx into the large intestine available for the microbiome, without needing to specifically track biliary secretion and reabsorption. I therefore model cobalamin influx into the large intestine directly as equal to dietary cobalamin intake, i.e. for each NHANES individual $i$,

$$J^{\text{LI}}_{B12,i} = D_{B12,i}, \qquad D_{B12,i} = \texttt{DR1IVB12}.$$

Notably, this argument concerns long-term average fluxes and does not imply a constant daily influx. Dietary intake and biliary secretion vary over time, so short-term $B_{12}$ delivery to the large intestine is also expected to fluctuate.

**Average dietary influx and variation** NHANES 2017–2018 data give a population mean dietary cobalamin intake of 4.64 $\mu$g/day (median 3.37 $\mu$g/day, 5th–95th percentile 0.51–12.54 $\mu$g/day) among US adults, reflecting substantial inter-individual variation.

Under the steady-state approximation above, this corresponds to a typical cobalamin influx into the large intestine of a few micrograms per day.

**From cobalamin to corrinoids** The previous argument applies specifically to canonical cobalamin. For microbial ecology, however, the broader corrinoid pool is relevant. Non-cobalamin corrinoids have been identified in several foods and can even constitute the predominant corrinoid in particular food sources. However, canonical cobalamin appears to predominate across the major conventional dietary sources of vitamin $B_{12}$, including meat, dairy products, fish, and most shellfish, whereas high abundances of non-cobalamin corrinoids have been reported primarily for particular food sources [55, 56]. I am not aware of systematic measurements quantifying the total dietary or biliary influx of non-cobalamin corrinoids, preventing an analogous estimate of total corrinoid influx into the large intestine. Available food-composition data do not suggest that non-cobalamin corrinoids contribute a dietary influx orders of magnitude larger than canonical cobalamin. I therefore expect total dietary corrinoid influx to be of a similar order of magnitude to the cobalamin influx estimated above, before accounting for corrinoid production within the large intestine itself. This resource is accordingly not currently included in the quantitative $J^{\text{LI}}_{r,i}$ reconstruction (Section 8).

**Comparison with fecal measurements** Direct measurements of human feces provide an important complementary constraint. One study quantified approximately 1.3 $\mu$g corrinoid equivalents $\text{g}^{-1}$ wet feces, corresponding to roughly 100–300 $\mu$g corrinoid equivalents/day depending on fecal output [57]. Remarkably, more than 98% of these compounds consisted of corrinoid analogues rather than canonical cobalamin. Earlier 24-hour stool collections reached similar conclusions, reporting only approximately 5 $\mu$g/day canonical cobalamin but nearly 100 $\mu$g/day apparent corrinoid analogues [58].

These measurements suggest that while canonical cobalamin entering the large intestine is largely constrained by dietary intake, the total corrinoid pool exported from the colon exceeds dietary cobalamin influx by roughly one to two orders of magnitude. This large imbalance points to substantial microbial corrinoid synthesis within the large intestine, accompanied by remodeling and exchange. This interpretation is also consistent with in-vitro fecal-community experiments showing de novo $B_{12}$ accumulation even in the absence of exogenous $B_{12}$ supplementation [51].

**Summary and confidence classification** The average influx of cobalamin into the large intestine can be estimated with moderate confidence: human cobalamin turnover is well understood, and long-term mass conservation allows dietary intake to serve as a reasonable proxy for intestinal influx, giving a working range of 0.5–12.5 $\mu$g/day (5th–95th percentile) with a population mean of 4.6 $\mu$g/day. In contrast, the external influx of non-cobalamin corrinoids, an ecologically important quantity for the microbiota, can currently only be assessed with unknown confidence: no direct influx measurements exist, and fecal corrinoid measurements reflect total extractable corrinoids without distinguishing intracellular bacterial corrinoids from freely dissolved luminal molecules available for uptake. Future studies directly measuring cobalamin and corrinoid fluxes entering the large intestine would be informative, particularly quantitative LC-MS characterization of ileal effluent from terminal ileostomy patients in controlled dietary setups. Combined with dietary intake, biliary physiology, and fecal measurements, such studies could establish corrinoid mass balances much more precisely, and quantify the rates of microbial corrinoid synthesis, remodeling, recycling, and export within the human gut ecosystem.

##### 2.5.2 Other vitamins and trace elements

Beyond cobalamin and corrinoids, other vitamins and cofactors can be highly relevant for microbial growth in the gut, including thiamine (B1), riboflavin (B2), niacin (B3), pantothenate (B5), pyridoxine (B6), biotin (B7), folate (B9), vitamin K, and heme. For example, a systematic genome-based assessment of eight B-vitamin biosynthesis pathways across 256 common human gut bacteria found that biosynthetic capacity varies substantially by vitamin and by species, with biotin, cobalamin, and folate biosynthesis genes notably absent or incomplete in many taxa, while pathways for niacin, pantothenate, pyridoxine, riboflavin, and thiamine are more broadly conserved [59, 60]. Such findings are in line with the supply of vitamins required in minimal media to enable support of many bacteria.

As such, the influx of vitamins other than cobalamin would be important to estimate as well.

Although dietary intake and intestinal absorption mechanisms are well characterized for many vitamins, I am not aware of systematic quantitative reconstructions of their net influx into the human large intestine across diets. Since I focus on corrinoids in the main text, I do not attempt to provide quantitative influx estimates for other vitamins here and instead flag this as a valuable direction for future work, including the use of modern metabolomics approaches to systematically quantify vitamin concentrations and fluxes in ileal effluent. This resource is accordingly not currently included in the quantitative $J_{r,i}^{\mathrm{LI}}$ reconstruction (Section 8).

**Summary and confidence classification** I classify the influx of vitamins other than cobalamin into the large intestine as **unknown**: no quantitative reconstruction is attempted here, pending future systematic measurements of vitamin concentrations and fluxes in human ileal effluent.

#### 2.6 Polyphenols

Beyond the macronutrients discussed above, dietary polyphenols constitute a further, quantitatively substantial class of plant-derived compounds reaching the large intestine, with increasingly well-characterized relevance to gut microbial ecology.

**Definition and dietary sources** Polyphenols comprise a chemically diverse class of plant secondary metabolites, broadly classified into flavonoids (flavonols, flavones, flavanones, isoflavones, anthocyanins, flavan-3-ols/catechins) and non-flavonoid compounds (phenolic acids, tannins, stilbenes, lignans). Dietary sources are predominantly plant-based (fruits, vegetables, whole grains, and legumes) with coffee and tea contributing a large share of total intake in Western populations.[JC: CITE]

**Total dietary intake** NHANES 2013–2016 data give a mean total polyphenol intake of 1657 mg/day among US adults, dominated by phenolic acids (1006 mg/day, with coffee contributing $\sim$40% of total polyphenol intake) and flavonoids (379 mg/day) [61]. This is broadly consistent with independent estimates across European populations using different dietary assessment methods, which span approximately 600–1800 mg/day [62]. Thus, despite substantial variation among populations and diets, total dietary polyphenol intake is consistently on the order of 1 g/day.

**Fraction reaching the large intestine** The fraction of consumed polyphenols escaping small-intestinal absorption varies substantially among compounds and food sources, as demonstrated directly in controlled ileostomy studies. Following apple juice consumption, approximately 90% of ingested procyanidins were recovered in ileal effluent, indicating very limited absorption of these oligomeric compounds in the small intestine [63]. In contrast, an ileostomy study of olive-oil phenols estimated that at least 55–66% of the administered phenol dose was absorbed in the small intestine, although recovery and apparent absorption differed substantially among individual compounds [64]. Following consumption of Concord grape juice containing a mixture of phenolic and polyphenolic compounds, approximately 40% of the ingested dose was recovered intact in ileal effluent [65]. Together, these measurements demonstrate substantial compound- and food-dependent variation and preclude assigning a well-supported universal escape fraction to dietary polyphenols.

For an approximate estimate of the magnitude of polyphenol influx, I use 50% as an illustrative escape fraction. This value is not intended as a population-weighted estimate of polyphenol absorption, but lies within the broad range observed in the direct ileostomy measurements discussed above. For individual $i$, this gives

$$J^{\mathrm{LI}}_{\mathrm{poly},i} = f_{\mathrm{escape}}\, D_{\mathrm{poly},i}, \qquad f_{\mathrm{escape}} \approx 0.5,$$

where $D_{\mathrm{poly},i}$ is dietary polyphenol intake. Note that, unlike every other NHANES-derived resource in this work, $D_{\mathrm{poly},i}$ is not available for the 2017–2018 cycle used elsewhere; the population-level intake figures below instead come from the NHANES 2013–2016 cycle [61], so this resource is not currently integrated into the same per-individual reconstruction as the other dietary resources.

Applying this illustrative fraction to the mean NHANES intake of 1.66 g/day gives approximately 0.8 g/day entering the large intestine. For comparison, applying the same assumption to the approximately 0.6–1.8 g/day dietary intake observed across European populations would correspond to approximately 0.3–0.9 g/day. These calculations suggest that dietary polyphenol delivery to the large intestine is plausibly on the order of several hundred milligrams per day, while the available data do not support a more precise estimate of the total influx.

Importantly, this influx should not be interpreted as equivalent to an equal mass of fermentable carbon. The extent to which individual polyphenols can serve as carbon sources, energy substrates, electron acceptors, or otherwise influence microbial physiology varies strongly among compounds and microbial taxa. Thus, while polyphenols constitute a quantitatively substantial chemical influx into the large intestine, the contribution of this influx to microbial carbon and energy supply cannot be inferred directly from its mass.

**Bacterial catabolism** Gut bacteria transform polyphenols through a diverse set of reactions, including ring cleavage, dehydroxylation, reduction, and decarboxylation, generating smaller phenolic metabolites that can subsequently undergo further microbial metabolism or host absorption. Particularly notable is catechol dehydroxylation by members of the *Coriobacteriia*, including *Eggerthella lenta*. Work on the molybdenum-dependent catechol dehydroxylase system demonstrated that catechol compounds can serve as alternative electron acceptors and that this metabolism can support bacterial growth [66]. This demonstrates that at least some dietary phenolic compounds can function as microbial electron acceptors, providing a conceptual connection to the electron-acceptor influxes discussed in Section 7, although the energetic consequences are likely substrate- and organism-dependent. Other transformations, including decarboxylation of hydroxycinnamic acids, can additionally contribute to resistance against the antimicrobial effects of phenolic compounds. Thus, microbial transformation of polyphenols can reflect both resource utilization and detoxification, depending on the compound and organism.

**Host uptake** Microbial degradation of polyphenols generates numerous smaller phenolic metabolites, many of which can subsequently be absorbed across the colonic epithelium and enter host circulation [65].

Thus, as for other resources considered here, the amount reaching the large intestine represents an influx into a dynamic microbial–host transformation system rather than the amount ultimately excreted in feces. Quantitatively resolving these downstream transformation and absorption fluxes remains substantially more difficult than estimating dietary intake.

**Summary and confidence classification** The compound classes, dietary sources, and population-level intake of polyphenols are relatively well characterized, with total dietary intake consistently on the order of 1 g/day. Direct human ileostomy studies further establish that substantial fractions of at least some dietary polyphenol classes reach the terminal ileum. Thus, the available evidence supports polyphenols as a quantitatively non-negligible chemical influx into the large intestine, plausibly on the order of hundreds of milligrams per day.

However, the fraction escaping small-intestinal absorption varies strongly among compounds and food sources, and no population-representative analysis currently combines compound-resolved dietary intake with corresponding ileal escape fractions. I therefore classify the *total influx* of polyphenols into the large intestine as **low confidence**. For illustration, assuming that 50% of the mean US dietary intake of 1.66 g/day reaches the large intestine gives an influx of approximately 0.8 g/day, but this value should be interpreted as an order-of-magnitude estimate rather than a physiological population mean. A food-composition-database approach linking compound-class-resolved escape fractions to population dietary intake data, analogous to the approach used for resistant starch and protein digestibility elsewhere in this work (Sections 2.1.1, 2.2), would substantially improve this estimate.

### 3 Host-derived resources entering from the ileum

As second major category of resources I consider resources supplied by the host. In contrast to the dietary resources discussed above, they originate from release by the body itself rather than directly from food. In the following I distinguish two subcategories: (i) endogenous resources that enter similarly to dietary resources, at the ileum, and (ii) endogenous resources entering through the gut epithelium along the large intestine. Unlike the very large number of distinct dietary compounds discussed above, the major endogenous compounds released into the ileum are comparatively few and well characterized, comprising primarily mucin, endogenous protein and amino acids, bile acids, and urea and ammonium. I consider each of these in turn below, before turning to the second subcategory of resources entering directly across the epithelium of the large intestine.

#### 3.1 Mucus components entering from the ileum

Mucin is an important host-derived carbon and nitrogen source entering the large intestine. Mucins are continuously secreted by the host as the principal structural component of the intestinal mucus layer, which gets renewed. Mucins therefore provide a persistent resource that is less dependent on dietary intake.

**Measuring mucin-related carbohydrates entering the large intestine** The amount of mucin-derived material entering the large intestine has been quantified through the quantification of mucin-specific components in ileal effluent, specifically different carbohydrates [67].

One of the first studies estimated from the measured loss of hexosamine (containing the amino sugars glucosamine and galactosamine, common in mucus) in the gut that this loss corresponds to about 3–4 g mucin per day under fasting conditions [13]). A different study estimated about 1.5 g mucin carbohydrates per day for ileostomists consuming a fiber-free diet [68]. Similar numbers were confirmed by another study estimating, through GalNAc and GlcNAc measurements, the mucus loss in ileal effluent, reporting an average of 3 g mucin-related carbohydrates per day, corresponding to less than 4 g mucin per day [69]. Notably, a study of ulcerative colitis ileostomy patients, based on the measurement of *N*-acetylgalactosamine content recovered in ileal effluent across twelve ileostomy patients, suggests an average amount of mucin carbohydrates of about 15 g per day leaving the small intestine [70].

**Mucin proteins** Typical mucin contains 20% protein and 80% carbohydrate by weight. Thus, from the roughly 4 g of mucin entering the large intestine each day, less than 1 g is protein.

**Bacterial mucus utilization in the large intestine** Of the mucus-related carbohydrates, only a tiny fraction is detected in feces, suggesting that most mucin entering the colon, together with mucus secreted by the colon itself, is degraded by the resident microbiota before excretion [70].

**Mapping mucus supply** Here I assume that, independent of diet, 4 g of mucus reaches the large intestine each day. I adopt 4 g/day as representative of the three more consistent estimates, specifically noting that this is only a proxy. Without more systematic investigations of mucus fluxes in controlled dietary studies, I do not explicitly model the potentially strong dependence on diet. Formally,

$$J^{\mathrm{LI}}_{\mathrm{mucin},i} = c_{\mathrm{mucin}} = 4 \text{ g/day},$$

a population-wide constant independent of $D_i$.

**Summary and confidence classification** Overall, mucin influx from the small intestine can be estimated with low confidence. Several independent measurements support an influx on the order of a few grams per day, motivating my working value of approximately 4 g/day, but the estimates vary substantially among studies and analytical approaches, with at least one considerably higher reported value. Furthermore, systematic measurements of how ileal mucin flux varies with diet are lacking. Future measurements of human mucus turnover and ileal mucin delivery using modern molecular and isotope-based approaches would substantially improve this estimate.

#### 3.2 Endogenous amino acids entering from the ileum

Besides mucus proteins, digestive enzymes and other proteins can also be part of the endogenous amino acid pool that enters the large intestine.

The total amount of endogenous amino acids has been measured well in fasting conditions or when consuming low-nitrogen diets, also due to its importance in estimating true ileal digestibility of consumed dietary amino acids [71, 32]. For example, Chacko and Cummings measured approximately 0.91 g nitrogen/day reaching the terminal ileum on a low-nitrogen diet, including approximately 2.7 g/day protein and a smaller contribution from free amino acids [30]. Rowan et al. reported a similar endogenous amino-acid loss, corresponding to approximately 1.4 g total amino acids per major meal [29].

Most of these amino acids likely arrive in the form of intact proteins and peptides, not free amino acids, as most amino acids exit in the form of proteins and peptides (see SI Section 2.2).

**Endogenous amino acids in non-fasting conditions** Notably, while most endogenous protein loss has been measured under fasting or low-nitrogen conditions, a few studies have suggested that endogenous protein secretion (or, more generally, endogenous amino acid secretion) might vary substantially with diet, for example because of variation in the amount of digestive enzymes secreted.

**Mapping endogenous amino acid amounts reaching the large intestine** To account for endogenous amino acid flux in my quantitative resource mapping, I assume a fixed influx equivalent to approximately 3 g/day of endogenous protein into the large intestine, of which approximately 1 g/day is assigned to mucosal origin. Formally,

$$J^{\mathrm{LI}}_{\text{endogAA},i} = c_{\text{endogAA}} = 3 \text{ g/day},$$

a population-wide constant independent of $D_i$. This endogenous contribution is kept separate from the dietary protein influx estimated above using true ileal digestibility. By definition, true ileal digestibility corrects measured ileal amino acid losses for basal endogenous losses and therefore estimates the dietary contribution independently of this endogenous pool. The principal remaining uncertainty is that endogenous secretion itself can depent on diet and food composition, whereas in the simplest model I represent it here as a fixed flux. Consequently, my separation of dietary and endogenous protein should be regarded as approximate, even though their combined order of magnitude is comparatively well constrained by direct ileal measurements.

**Summary and confidence classification** Overall, the total endogenous amount can be estimated with moderate confidence, higher than for mucin, since it rests on a substantially larger body of true ileal amino acid digestibility measurements. Together with dietary protein, the total protein influx can be well estimated. What is more difficult to measure, and currently impossible to model explicitly with the available data, is the shift in endogenous protein secretion with diet, which may partly replace the influx of dietary protein; as noted above, however, this uncertainty affects the split between dietary and endogenous origin rather than the total combined flux, which remains comparatively robust.

Nevertheless, important opportunities remain. Existing digestibility studies have primarily focused on protein quality for the host rather than microbial resource supply, and the dependence of protein delivery on overall diet composition, food processing, and eating habits has not yet been systematically analyzed from a microbiome perspective.

#### 3.3 Bile acids entering from the ileum

Another important class of endogenous resources is bile acids, host-derived molecules that are secreted into the small intestine and extensively recycled through the enterohepatic circulation. Bile acids facilitate lipid digestion and absorption, contribute to cholesterol homeostasis, and provide an important route for the excretion of cholesterol, bilirubin, drugs, and other metabolites. At the same time, bile acids constitute an important ecological factor shaping the gut microbiome through their antimicrobial activity [72], while gut bacteria in turn extensively modify bile acid chemistry, thereby influencing host physiology and the composition of bile acids entering and passing through the large intestine. A broad literature on bile acid physiology allows their turnover along the intestinal tract to be reconstructed quantitatively, including delivery from the ileum to the large intestine and eventual loss through feces.

**Bile acid turnover and intestinal flux** Amounts of bile acids secreted into the small intestine are typically of the order of 10–35 g/day, corresponding to a bile acid pool of roughly 2–4 g that recycles multiple times each day through the enterohepatic circulation [73, 74]. The great majority of secreted bile acids are reabsorbed by the host, primarily in the terminal ileum [75]. This is supported by studies measuring bile acid loss in ileal effluent from ileostomy patients. Specifically, Stiehl et al. report a mean daily loss of approximately 1700 $\mu$mol/day ($\approx 0.8$ g/day for a representative bile acid molecular weight of 475 g/mol) [76]. Other ileostomy studies report values between roughly 750 and 1800 $\mu$mol/day ($\approx 340$–900 mg/day), confirming the order of magnitude [77, 78].

Measurements in feces provide a consistent picture, although daily bile acid losses are lower than ileal delivery, consistent with substantial continued absorption of bile acids in the large intestine. Specifically, Cummings et al. report values ranging from approximately 90 to 430 mg/day ($\approx 180$–960 $\mu$mol/day) under different dietary conditions [35], while Natori et al. report $235 \pm 61$ mg/day in healthy controls [79]. Setchell et al. further demonstrated substantial day-to-day variation in fecal bile acid excretion within individuals, with fluctuations on the order of 50–100 mg/day ($\approx 100$–220 $\mu$mol/day) [80].

Cummings et al. further investigated the dependence of fecal bile acid loss on dietary fat using controlled dietary studies, reporting approximately $140 \pm 63$ mg/day for a low-fat diet (62 g fat/day) and $320 \pm 120$ mg/day for a high-fat diet (152 g fat/day) [35]. These observations are consistent with the physiological role of bile acids in facilitating fat digestion and suggest a higher bile acid turnover when more fat is consumed.

**Bile acid modifications by bacteria** Importantly, while I focus in this article on the total flux of bile acids entering the large intestine, microbial activity substantially modifies bile acids, giving rise to a remarkable diversity of bile acid-derived metabolites. Some of these modifications already begin in the small intestine. I therefore briefly summarize these processes here.

Analyses of localized samples from different parts of the intestine have shown that bile acids remain predominantly glycine- or taurine-conjugated throughout the duodenum and jejunum, whereas bacterial deconjugation becomes substantial toward the distal small intestine, with unconjugated bile acids accounting for up to approximately half of the bile acid pool in the lower ileum [81, 77]. Recent spatial metabolomic analyses of the human intestinal tract using ingestible sampling capsules further demonstrate pronounced spatial gradients in bile acid composition, providing independent support that microbial bile acid modification begins before the large intestine [82]. By the time bile acids reach the human cecum, approximately 90% are unconjugated [83]. This deconjugation is mediated by bacterial bile salt hydrolases [84, 85].

More recently, metabolomic analyses have transformed my view of bile acids from a relatively small family of host-derived molecules involved primarily in lipid digestion into a highly diverse chemical space jointly shaped by host and microbial metabolism. Besides deconjugation, classical microbial transformations include $7\alpha$-dehydroxylation, oxidation, and epimerization [86, 85]. Recent metabolomic studies have revealed an even broader diversity of microbially modified bile acids, including conjugation with a variety of amino acids and numerous additional bile acid derivatives [87, 88, 85]. Although some microbial transformations already occur in the distal small intestine, extensive diversification of the bile acid pool continues during passage through the large intestine. Together, these transformations constitute a remarkable example of host–microbiome co-metabolism.

**Modeling bile acid flow** Because direct measurements of how ileal bile acid delivery varies with dietary fat intake are not available, I use the observed dependence of fecal bile acid excretion on dietary fat as a proxy for its relative diet dependence, while independently anchoring the absolute magnitude of the influx to direct ileostomy measurements [35, 76, 77].

The two dietary conditions reported by Cummings et al. give an approximately linear relationship between dietary fat intake $T$ and fecal bile acid loss,

$$J_{\text{BA,fecal}}(T) \approx 15.5 + 2.00T \quad \text{mg/day},$$

where $T$ is dietary fat intake in g/day, as reported for the two conditions of that study. To estimate ileal delivery from this relative diet dependence, I rescale this relationship to the absolute magnitude indicated by the direct ileostomy measurements. Taking approximately 1.5 mmol/day, or 0.7 g/day for a representative bile acid molecular mass of 475 g/mol, as a representative ileal influx, and approximately 0.23 g/day as a representative fecal loss based on the measurements above, gives a scaling factor of approximately

$$\alpha \approx \frac{0.7}{0.23} \approx 3.$$

I therefore approximate diet-dependent bile acid influx into the large intestine, for each NHANES individual $i$ with dietary fat intake $T_i$ (Section 2.3, `DR1ITFAT`), as

$$J^{\text{LI}}_{\text{BA},i} \approx 3\,(15.5 + 2.00\,T_i) \quad \text{mg/day}.$$

Across the two dietary fat conditions measured by Cummings et al. ($T = 62$ and 152 g/day), this relationship predicts bile acid influxes of approximately 420 and 960 mg/day, respectively, consistent with the range observed in direct ileostomy measurements.

This scaling assumes that the ratio between ileal bile acid delivery and fecal bile acid loss remains approximately constant across dietary fat intakes. Because fecal bile acid output additionally depends on colonic reabsorption, transit, and microbial transformation, this relationship should be interpreted as a phenomenological first-order approximation rather than a direct measurement of diet-dependent ileal influx. In the absence of more systematic data, I do not model additional dependencies on other dietary components.

**Summary and confidence classification** There is a strong mechanistic understanding of bile acid synthesis in the liver, their efficient reabsorption in the terminal ileum, and their extensive modification by the gut microbiota, though only a handful of studies have directly quantified the flux of bile acids entering the large intestine. Relatively little is known about how this influx varies with diet, as dietary effects are inferred primarily from fecal measurements rather than direct measurements of ileal efflux. I therefore assess the influx of bile acids into the large intestine, and particularly its dependence on diet, with moderate confidence. Direct measurements support a working range of roughly 1–2 mmol/day ($\approx 450$–950 mg/day), while my phenomenological fat-dependent model gives a similar range of approximately 0.4–1.0 g/day across the dietary fat intakes examined above. Future studies extending direct measurements of ileal bile acid delivery and coupling them with recent advances in understanding microbial bile acid metabolism could quantify the rates of individual bile acid transformations and systematically relate bile acid metabolism to diet and microbiome function.

##### 3.4 Urea and ammonium entering from the ileum

The outflux of urea and ammonium from the ileum has also been measured. Across different diets, studies with ileostomists reported an efflux of 0.6–1.2 mmol/day for ammonium and 0.5–1.9 mmol/day for urea [30]. Together, urea and ammonium account for only 3–5% of the total ileal nitrogen efflux measured. Miner-Williams et al. reported a similar range, with ammonium and urea together accounting for up to 9% of all observed nitrogen [71].

**Mapping urea and ammonium into the large intestine** Given the small magnitude of this flux relative to dietary and endogenous amino acid influx, and the absence of systematic diet-dependence data, I model urea and ammonium entering via the ileum as a fixed influx, using the values reported above. Formally,

$$J^{\mathrm{LI}}_{\text{urea-ileal},i} = c_{\text{urea-ileal}} = 1.2 \text{ mmol/day}, \qquad J^{\mathrm{LI}}_{\mathrm{NH_4}\text{-ileal},i} = c_{\mathrm{NH_4}\text{-ileal}} = 0.9 \text{ mmol/day},$$

both population-wide constants independent of $D_i$.

**Summary and confidence classification** Overall, the low influxes of ammonium and urea entering via the ileum are quantified with moderate confidence, with a consistent range of approximately 0.5–2 mmol/day for both urea and ammonium. A possible variation with diet is not well established, but these influxes are low compared with the overall turnover of urea and ammonium in the large intestine (see Section 6.2).

#### 4 Trace metals and phosphorus

Beyond the major inorganic ions considered above, the microbiota requires trace metals and phosphorus for growth and metabolism. Iron and zinc are required in much smaller amounts but serve as essential cofactors for numerous cellular processes, whereas phosphorus is a major biomass-forming element with substantially larger anabolic demand.

##### 4.1 Iron

Beyond the major inorganic ions discussed above, iron is a further micronutrient of particular relevance to gut microbial ecology, given its essential role in bacterial redox enzymes and its increasingly well-documented influence on microbiome composition.

**Dietary intake and absorption** Typical dietary iron intake is on the order of 10–18 mg/day, predominantly as non-heme iron ($\sim$85–90% of total dietary iron in an omnivorous diet), with heme iron contributing the remainder. Small-intestinal absorption of non-heme iron is generally incomplete and varies strongly with host iron status and dietary composition. A systematic analysis of whole-diet studies reported mean non-heme iron absorption ranging from 0.7–22.9%, with values typically around 5–8% [89]. Thus, under typical dietary conditions, a substantial fraction of ingested iron escapes systemic absorption in the small intestine and can reach the distal gut.

**Ileal and fecal measurements** Direct measurements of iron reaching the terminal ileum are considerably more limited. In a metabolic-balance study of eight ileostomy patients, approximately 8 mg/day of iron was recovered in ileostomy effluent [90]. This magnitude is broadly consistent with the incomplete small-intestinal absorption expected from dietary iron intake and absorption measurements. Reported fecal iron measurements vary considerably among studies [91, 92]; however, the quantitative values reported in these studies require careful consideration of dietary iron intake, supplementation, analytical methodology, and units before they can be directly compared with ileal flux measurements.

**Colonic absorption capacity** Although the duodenum and proximal small intestine are the dominant sites of dietary iron absorption, the human colon retains a secondary capacity for iron absorption. Direct administration studies demonstrated that iron can be absorbed from the human large intestine [93]. More recent analyses of human intestinal biopsies identified the principal non-heme iron transporters DMT1 and ferroportin (FPN1) throughout the colon: DMT1 abundance was comparable to that in the duodenum, whereas FPN1 abundance was approximately one-quarter of duodenal levels [94]. In patients with active ulcerative colitis and iron deficiency, expression of both transporters was increased, and enhanced DMT1-dependent $^{59}Fe$ uptake provided direct functional evidence that colonic iron absorption can increase under these conditions [94]. Thus, although colonic iron absorption appears to make a comparatively minor contribution under normal physiological conditions, the large intestine possesses the transport machinery required for iron uptake and its absorptive capacity can respond to host iron status.

**Microbial competition and host regulation** Iron availability in the intestinal lumen is shaped not only by total iron influx but also by host sequestration and microbial competition for the available iron pool. During infection and inflammation, the host restricts microbial access to iron through nutritional-immunity mechanisms involving iron-binding and siderophore-binding proteins such as lactoferrin and lipocalin-2, while gut bacteria deploy diverse iron-acquisition systems, including siderophores, xenosiderophore uptake, and ferrous-iron transport [95, 96]. Competition for iron can therefore influence bacterial fitness and colonization in the gut. However, these mechanisms are best established in the context of pathogens and intestinal inflammation; the extent to which host-mediated iron restriction constitutes a bulk growth limitation for the resident microbiota in the healthy colonic lumen remains less well resolved.

**Summary and confidence classification** Overall, total iron influx into the large intestine can be estimated with **low confidence**. Dietary intake and small-intestinal absorption are reasonably well characterized, and the available ileostomy measurements place the influx on the order of several milligrams per day. However, direct human ileal measurements remain sparse, and iron availability is strongly affected by chemical speciation, host binding proteins, microbial uptake, and host iron status. Thus, the uncertainty in microbially available iron is substantially greater than the uncertainty in total iron mass entering the large intestine. Formally, $J^{\mathrm{LI}}_{Fe,i} = c_{Fe} = 8$ mg/day, a population-wide constant independent of $D_i$. Additional controlled ileostomy measurements across dietary iron intakes and host iron states would substantially improve this estimate.

#### 4.2 Zinc

Zinc is an essential trace element required by both the host and the gut microbiota, where it serves as a cofactor for numerous enzymes and regulatory proteins. Dietary zinc intake in adults is typically on the order of 10 mg/day, but intestinal absorption varies substantially with diet composition and host zinc status. In particular, phytate strongly reduces zinc bioavailability, making the chemical composition of the diet an important determinant of the amount of zinc escaping absorption in the small intestine.

**Ileal influx** Direct measurements in ileostomy patients provide a direct estimate of zinc delivery to the large intestine. Sandberg et al. measured dietary intake and ileostomy output of several minerals, including zinc, under controlled dietary conditions [90]. Zinc delivery to the terminal ileum was on the order of several to approximately 10 mg/day. Addition of 16 g/day wheat bran increased zinc delivery and significantly decreased apparent zinc absorption, consistent with the known inhibitory effect of phytate on zinc absorption. These measurements therefore establish both the approximate magnitude of zinc influx into the large intestine and its potential dependence on dietary composition.

**Fecal measurements** Independent measurements of fecal zinc excretion agree remarkably well with this order of magnitude. The compilation by [97] reports values of approximately 5–13 mg Zn/day across several studies, with individual estimates of 7.85, 5–10, 10.68, and 13.31 mg/day. Thus, both direct ileal measurements and fecal measurements consistently place zinc flux through the large intestine on the order of 10 mg/day.

The comparison between ileal and fecal measurements should nevertheless not be interpreted as a simple balance of unabsorbed dietary zinc. Gastrointestinal secretion and reabsorption of endogenous zinc contribute to whole-body zinc homeostasis, and fecal zinc therefore contains both unabsorbed dietary zinc and endogenous zinc. Moreover, the total zinc flux quantified here does not necessarily correspond to the zinc available to microorganisms, because zinc speciation and binding to dietary and host-derived compounds can strongly affect its bioavailability.

**Summary and confidence classification** Overall, direct ileostomy measurements place total zinc influx into the large intestine on the order of several to approximately 10 mg/day, while fecal measurements of approximately 5–13 mg/day provide an independent order-of-magnitude consistency check. Because fecal zinc additionally includes endogenous secretion and reflects subsequent colonic handling, it should not be interpreted as a direct measure of ileal influx. The principal remaining uncertainty concerns diet-dependent variation, particularly the strong effects of phytate and other dietary components on zinc absorption, and the distinction between total and microbially available zinc. I therefore classify total zinc influx into the large intestine as moderate confidence, while the concentration or flux of bioavailable zinc available for microbial growth is considerably less well constrained. Formally, $J^{\mathrm{LI}}_{Zn,i} = c_{Zn} = 10$ mg/day (range 5–13 mg/day), a population-wide constant independent of $D_i$.

##### 4.3 Phosphorus

Phosphorus is another element required in large quantities to support microbial growth, serving as a major component of nucleic acids, phospholipids, ATP, and many phosphorylated metabolites.

**Dietary sources** Unlike several other resources discussed in this work, dietary phosphorus does not reach the colon primarily because a single dominant chemical form escapes digestion uniformly – rather, its fate depends strongly on which chemical form it is consumed in. Phytate (myo-inositol hexakisphosphate), the principal storage form of phosphorus in plant seeds, grains, legumes, and nuts (representing 60–90% of total phosphorus in some seeds), is physiologically analogous in this respect to resistant starch (Section 2.1.1): humans lack sufficient endogenous phytase to liberate its bound phosphate, so phytate-phosphorus is substantially less bioavailable than many other dietary phosphorus forms and a substantial fraction can reach the distal gut without absorption [98]. By contrast, phosphorus from animal protein sources (meat, dairy, eggs) is substantially more bioavailable ($\sim$40–60% absorbed).

Inorganic phosphate food additives were long assumed to be even more bioavailable still ($\sim$90%+), based on in vitro digestibility assays; however, controlled-feeding studies comparing dietary phosphorus source with 24-hour urinary phosphorus recovery, which under controlled steady-state conditions provides a physiological proxy for absorbed dietary phosphorus [99], have found that food-additive phosphorus is, contrary to this expectation, incompletely absorbed [100]. This discrepancy between in vitro digestibility and urinary-recovery measurements is itself informative: it suggests that digestibility-based bioavailability estimates for inorganic additive phosphate may not translate directly to the fraction actually reaching systemic circulation. As such, it also leaves open the possibility that a non-trivial fraction of additive phosphorus escapes small-intestinal absorption despite its high aqueous solubility.

More broadly, across the typical adult intake range, net phosphorus absorption efficiency is close to constant, with reported values of 55–70% (most commonly cited as 60–65%) showing no significant dependence on intake level; in the primary balance studies underlying this estimate, the intercept of the regression of fecal phosphorus on dietary phosphorus intake is not significantly different from zero, and the relationship remains linear up to at least 3.1 g/day intake [99]. This is a notable contrast with calcium, for which absorption efficiency increases adaptively as intake falls; no comparable adaptive mechanism has been identified for phosphorus. At the level of a typical mixed diet, these measurements imply that approximately 30–45% of dietary phosphorus escapes net absorption, with a typical value around 35–40%. As such, phosphorus delivery can be reasonably approximated as a near-constant fraction ($\sim$35–40%) of dietary intake, without requiring a diet-level correction for absorption efficiency itself, though this uniform-fraction relationship still leaves open substantial food-composition-driven variation in *which fraction of intake* is bioavailable in the first place (the plant/animal/additive distinction above).

This suggests that diet composition – specifically the balance of plant, animal, and processed-food phosphorus – may be at least as important a determinant of colonic phosphorus delivery as total phosphorus intake, though I am not aware of a study directly quantifying this compositional effect on ileal or colonic phosphorus flux at the population level. It is also worth noting that, at the population level, variation in total phosphorus intake (mg/day) appears to be driven substantially by differences in total food intake rather than by differences in food composition per se, since phosphorus content scales fairly consistently with energy content across a mixed diet ($\sim$62 mg P per 100 kcal on average) [99]. Composition-driven variation in *bioavailability* – and hence in the fraction of ingested phosphorus reaching the colon – is therefore likely to be a distinct and largely independent source of variation from composition-driven variation in raw intake.

Consistent with active microbial participation in phytate degradation, experiments in germ-free and conventionally colonized animals have demonstrated that intestinal microorganisms can make a major contribution to phytate hydrolysis. In one classical study, phytate hydrolysis was negligible in germ-free rats but reached 22–56% in conventionally colonized animals, depending on dietary conditions [101]. Recent mechanistic work further demonstrated microbial degradation of phytate and cross-feeding of phytate-derived metabolites between gut bacterial species, ultimately supporting production of propionate [102].

**Ileal and fecal measurements** Two ileostomy studies report phosphorus fluxes of approximately 0.25–0.5 g P/day leaving the ileum [103, 90]. Both quantified phosphorus by wet-ashing ($H_2SO_4$/$H_2O_2$) followed by the colorimetric method of Fiske & Subbarow (1925); this digestion step converts organic and inorganic phosphorus alike to orthophosphate prior to colorimetric determination, so the reported values should be read as genuine total phosphorus regardless of chemical form or whether the phosphorus was free in solution or incorporated into bacterial biomass at the time of collection.

Independently, a balance-study-based estimate of fecal phosphorus can be derived from the intake-dependent absorption relationship described above: applying a $\sim$60–65% absorption fraction to typical adult intake levels (1.0–1.4 g/day) predicts fecal phosphorus output of approximately **0.35–0.56 g/day**. Several studies have also directly measured total phosphorus in human feces across more heterogeneous populations and methods, reporting a considerably higher pooled median of $\sim$0.9 g/day, with substantial variation ranging from 0.25 to 2.5 g/day across individuals, diets, and studies [97]. The fecal values compiled in [97] are drawn from a heterogeneous set of source studies without a uniform reported method or matched dietary intake, so some of this discrepancy with the balance-study-based estimate may reflect differences in study population, diet, or measurement approach rather than a genuine difference in typical fecal phosphorus output; I discuss this further below.

Only [90] further partitioned ileal-effluent phosphorus into phytate-P and non-phytate-P, using Fe-precipitation of intact phytate followed by P determination on the precipitate, with non-phytate-P taken as the difference from total P. On the low-fibre control diet, phytate-P made up only a small fraction of total ileal-effluent P: approximately 2% (period 1: 0.009 of 0.41 g P/day; period 3, three additional patients: 0.009 of 0.48 g P/day). Adding 16 g wheat bran/day raised this to roughly 11% (period 2: 0.059 of 0.54 g P/day), consistent with bran phytate only being partially digested before the ileum. Non-phytate-P – comprising inorganic phosphate plus organically bound but non-phytate P (protein-, nucleotide-, and phospholipid-derived) – therefore dominates ileal effluent P even under bran supplementation. These figures should be treated as rough and study-specific (n = 8 ileostomy patients, one analytical method, one dietary pattern): the Fe-precipitation assay targets intact phytate (IP6) and may undercount partially dephosphorylated phytate breakdown products (IP3–IP5), so the true phytate-derived fraction of ileal P could be somewhat higher than these figures suggest, with the excess misclassified as non-phytate-P.

**Limited evidence for epithelial phosphate secretion** Unlike urea, bicarbonate, potassium, and sulfate discussed elsewhere in this work, there is little evidence for quantitatively important active phosphate secretion by the colonic epithelium into the lumen. Intestinal phosphate transport – predominantly paracellular, with a minor and still-debated active (NaPi-mediated) component – is primarily absorptive throughout the intestine, including the colon, though colonic absorptive capacity is substantially smaller than that of the small intestine and is generally considered physiologically minor except under high luminal phosphate concentrations [104, 105, 106]. I therefore find no evidence for a quantitatively important host-driven phosphate secretion into the large-intestinal lumen analogous to the secretory fluxes of urea, bicarbonate, or potassium discussed elsewhere in this work. Colonic phosphate absorption, conversely, provides a potential route for a modest decrease in luminal phosphorus flux.

Given this, fecal total phosphorus output should, to first order, approximate net phosphorus delivery to the colon via the ileum, whether that phosphorus passes through unchanged or is transiently incorporated into and released from bacterial biomass along the way – since total-phosphorus assays (Fiske-Subbarow or equivalent) do not distinguish these fates. This makes fecal phosphorus a reasonable proxy for colonic phosphorus influx, with the caveat that it reflects net rather than gross flux if any colonic host absorption occurs.

Comparison of the direct ileal measurements ($\sim$0.25–0.5 g/day) with the pooled fecal value ($\sim$0.9 g/day, [97]) appears to suggest an increase in total phosphorus during colonic transit. However, microbial incorporation cannot itself explain such an increase in total phosphorus flux, because incorporation into and release from bacterial biomass changes the chemical form of phosphorus but not its total mass. With the addition of the balance-study-based fecal estimate above ($\sim$0.35–0.56 g/day at typical intake), the apparent discrepancy is substantially reduced: the ileal measurements from [90] (0.41–0.54 g/day) fall within, or only modestly below, the independently predicted fecal range. I therefore consider differences in dietary intake, study population, and analytical methodology between the small controlled ileostomy studies and the heterogeneous fecal literature to be more plausible explanations for the higher pooled fecal values than a genuine increase in phosphorus flux within the colon. Direct matched measurements of ileal and fecal phosphorus within the same individuals would be required to resolve any remaining discrepancy.

**Relative contribution of different phosphorus sources to ileal delivery** Combining the bioavailability figures above with typical dietary composition suggests that dietary non-phytate organic phosphorus – particularly protein-bound phosphorus from animal sources, given its intermediate bioavailability ($\sim$40–60% absorbed) applied to what is often the largest single category of dietary phosphorus intake – may plausibly constitute the largest single dietary contributor to ileal phosphorus, rather than phytate-phosphorus as might be assumed from the fiber and mineral-binding literature. Phytate-phosphorus, despite being very poorly absorbed, occupies a comparatively small share of total phosphorus intake unless the diet is dominated by whole grains, legumes, nuts, or seeds; consistent with this, phytate-P made up only $\sim$2–11% of ileal-effluent P in [90] (see above), though that study's low-fibre background diet likely understates phytate's contribution relative to higher-fiber diets. Inorganic additive phosphate, despite its high aqueous solubility, may contribute more to ileal delivery than previously assumed given the urinary-recovery evidence of incomplete absorption discussed above; I treat its net contribution as uncertain rather than negligible, in contrast to earlier assessments based on digestibility data alone. The endogenous contribution (pancreatic and biliary secretions, sloughed enterocytes) is essentially unconstrained by direct measurement, but – unlike the dietary fractions above – is unlikely to scale strongly with intake, so its proportional importance may be larger in low-phosphorus-intake or low-fiber experimental diets, such as the controlled diets used in the available ileostomy studies, than in a typical higher-intake Western diet. I emphasize that this ordering is inferred from bioavailability estimates and coarse dietary composition rather than measured directly, and a study resolving ileal phosphorus by chemical source (for example via isotopic labeling of specific dietary phosphorus pools) would be needed to test it.

**Food sources of dietary phosphorus** Dairy products carry a disproportionately high phosphorus density relative to most other food groups, and colas and other soft drinks using phosphoric acid as an acidulant can contribute substantially to total intake when consumed in quantity (five or more servings/day), despite each serving containing only a small absolute amount (~50 mg per 12 oz serving, ~5% of typical adult female intake) [99]. More broadly, phosphorus content scales closely with energy content across a typical mixed diet, meaning that much of the population-level variation in absolute phosphorus intake reflects variation in total food intake rather than in food type per se [99]; this is distinct from, and should not be conflated with, food-composition-driven variation in phosphorus *bioavailability* discussed above.

**Modeling phosphorus reaching the large intestine** Building on the bioavailability considerations above, I implement three complementary mappings with different physiological assumptions to estimate phosphorus delivery to the large intestine across the NHANES 2017–2018 adult population.

**Simple mapping.** I first apply a single, population-wide fraction $f$ of dietary phosphorus intake $I_i$ (`DR1IPHOS`) directly to each person,

$$P_i^{\text{simple}} = f\, I_i, \tag{15}$$

with $f = 0.425$ (bracketed $0.30$–$0.55$), corresponding to the approximately 55–70% net absorption range reported for mixed diets.

**Plant/non-plant tiered mapping.** To capture the source-dependent bioavailability discussed above, I resolve each NHANES food code's phosphorus into a plant-derived fraction $\alpha_i \in [0, 1]$ and a non-plant (animal + additive) fraction $1 - \alpha_i$, using ingredient-level phosphorus data from FNDDS (falling back to WWEIA food-category classification for the 409 of 7083 food codes lacking ingredient-level phosphorus data).

This plant/non-plant split is necessarily coarse: plant-derived phosphorus includes both phytate and non-phytate forms, while the non-plant category combines animal-derived organic phosphorus with inorganic additives. I therefore use this mapping primarily as a sensitivity analysis rather than as a chemically resolved model of phosphorus absorption.

Plant and non-plant phosphorus are then each multiplied by their own source-specific escape fraction,

$$P_i^{\text{tiered}} = \alpha_i I_i\, f_{\text{plant}} \;+\; (1 - \alpha_i) I_i\, f_{\text{non-plant}}, \tag{16}$$

with $f_{\text{plant}} = 0.53$ (bracketed $0.48$–$0.59$) and $f_{\text{non-plant}} = 0.51$ (bracketed $0.28$–$0.64$, reflecting the additive-phosphorus absorption uncertainty discussed above). Averaged across the population, $39.6\%$ of dietary phosphorus is plant-derived.

**Calcium-dependent mapping.**

Following the balance-study regression of Heaney & Nordin [107], I instead predict fecal phosphorus and use it as a proxy for phosphorus delivery to the large intestine, based on the argument above that quantitatively important colonic phosphorus secretion is not known and colonic absorption appears limited.

$$P_i^{\text{Ca-dep}} = \text{clip}\Big(c_0 + c_f\, F_{\text{Ca},i} + c_p\, I_i,\ 0,\ I_i\Big), \tag{17}$$

with fecal calcium estimated from dietary calcium intake $I_{\text{Ca},i}$ (DR1ICALC) using the relationship reported by Heaney & Nordin between fecal calcium and calcium intake,

$$F_{\text{Ca},i} = 0.89\, I_{\text{Ca},i}.$$

The coefficient 0.89 is the reported slope of fecal calcium versus calcium intake, rather than a calcium-absorption fraction. This clipping is a pragmatic constraint imposed for the present influx mapping; the original regression itself can predict fecal phosphorus greater than dietary phosphorus under high Ca:P conditions, which the authors interpret in terms of the model's phosphorus-binding capacity rather than as a literal dietary mass-balance prediction.

**Comparison of the three mappings.** All three mappings give broadly comparable, and in two cases closely matching, population medians: $522$ mg/day (simple), $636$ mg/day (tiered), and $509$ mg/day (calcium-dependent). The simple and calcium-dependent mappings agree particularly closely (within $2.5\%$ at the median), despite the calcium-dependent mapping using no fitted population-wide escape fraction at all. The tiered mapping runs $\sim$25% higher, because its source-specific escape fractions ($0.53$ plant, $0.51$ non-plant), weighted by the population's actual $39.6\%$ plant share, imply a higher average escape fraction ($\sim$52%) than the simple mapping's flat $42.5\%$ – i.e. the discrepancy traces to the tiered mapping's absorption-fraction assumptions themselves, not to a difference in modeling structure. The calcium-dependent mapping's two independent datasets also agree closely with each other (dataset 1 median $509$ mg/day, 5th–95th percentile $203$–$1082$; dataset 2 median $524$ mg/day, $166$–$1166$), and its population median is consistent with the direct ileal and balance-study literature estimates discussed above ($\sim$350–560 mg/day). I adopt the calcium-dependent mapping as my primary working estimate because it is grounded in a directly fitted human balance-study regression and agrees closely with the available direct ileal measurements. I emphasize, however, that this regression predicts fecal phosphorus rather than ileal delivery itself; its use here as an influx proxy rests on the limited evidence for quantitatively important phosphate exchange across the colon discussed above. I adopt $J_{P,i}^{\text{LI}} = P_i^{\text{Ca-dep}}$ as the phosphorus influx carried forward into the elemental-conversion step (Section 9).

**Summary and confidence classification** Together, these measurements allow total phosphorus influx into the large intestine to be estimated with **moderate confidence**. Direct ileal measurements and human balance studies consistently place typical phosphorus delivery on the order of several hundred milligrams per day. Using my adopted calcium-dependent NHANES mapping, I estimate a 5th–95th percentile range of 0.20–1.08 g P/day, with a median of 0.51 g/day, across the US adult population. Considerably greater uncertainty remains regarding the chemical forms and dietary origins of this phosphorus influx and the fraction immediately accessible to microbial growth.

### 5 Abundance and exchange of major ions

Major inorganic ions constitute a distinct class of chemical inputs encountered by the gut microbiota. Unlike carbon, nitrogen, phosphorus, and sulfur, their cellular requirements are generally more flexible and they should therefore not be interpreted as stoichiometrically fixed biomass-building resources. Nevertheless, $Na^+$, $K^+$, $Ca^{2+}$, $Mg^{2+}$ and $Cl^-$ are present at substantial fluxes, contribute to the physicochemical environment of the gut, and perform essential cellular functions ranging from osmotic and electrochemical homeostasis to enzyme activity and macromolecular stabilization. I therefore quantify their influxes separately from the major biomass-forming elements.

#### 5.1 Potassium

Potassium influx into the large intestine has been well characterized. Across ileostomy studies, approximately 0.2–0.9 g/day (6–22 mmol/day) enters the large intestine [108]. Fecal potassium excretion is reported to range between approximately 0.2 and 1 g/day, with somewhat higher values occasionally observed [97]. As such, fecal potassium excretion is of a similar magnitude to the amount entering from the ileum, indicating that potassium is not subject to the strong net colonic absorption observed for sodium. This is consistent with the known combination of absorptive and secretory potassium transport pathways in the colon. Formally, $J^{\mathrm{LI}}_{K,i} = c_K = 0.55$ g/day, a population-wide constant independent of $D_i$.

#### 5.2 Sodium

Sodium enters the large intestine in substantially larger quantities than potassium. Approximately 2.8–3.5 g/day (120–150 mmol/day) has been reported to leave the ileum [108]. In contrast, only about 0.1 g/day is typically excreted in feces [97], reflecting the well-established active absorption of sodium throughout the large intestine.

Together with chloride and bicarbonate, sodium constitutes the dominant contributor to the osmolarity of intestinal contents, as in other extracellular fluids [97]. Formally, $J^{\mathrm{LI}}_{Na,i} = c_{Na} = 3.15$ g/day, a population-wide constant independent of $D_i$.

#### 5.3 Calcium

Sandberg et al. reported approximately 0.8 g/day calcium (20 mmol/day) entering the large intestine in ileostomy subjects [90].

Reported fecal calcium excretion is somewhat lower, approximately 0.5 g/day [97], consistent with net calcium absorption during colonic transit, although these measurements were not obtained in matched individuals and therefore do not permit a precise colonic mass balance. Dietary fiber composition may also modulate colonic calcium handling [109]. Formally, $J^{\mathrm{LI}}_{Ca,i} = c_{Ca} = 0.8$ g/day, a population-wide constant independent of $D_i$.

#### 5.4 Magnesium

Sandberg et al. reported approximately 0.12 g/day magnesium (5 mmol/day) entering the large intestine in ileostomy subjects [90]. Notably, this is lower than reported fecal magnesium excretion of approximately 0.2 g/day [97], the opposite pattern from potassium, sodium, and calcium above, each of which show fecal output at or below ileal influx.

Taken at face value, this comparison would imply net magnesium secretion into the colon. However, the ileal and fecal values derive from different study populations and methodologies, and I therefore do not interpret their difference as evidence for a quantitatively established secretory flux. Instead, I regard the direction of net colonic magnesium transport as poorly constrained by the available mass-balance data. Formally, $J^{\mathrm{LI}}_{Mg,i} = c_{Mg} = 0.12$ g/day, a population-wide constant independent of $D_i$.

#### 5.5 Summary and confidence classification for major inorganic ions

Overall, the influxes of potassium, sodium, calcium, and magnesium into the large intestine can be estimated with **moderate confidence**. Direct ileostomy measurements establish their approximate magnitudes, and independent fecal measurements provide useful constraints on subsequent colonic handling. Sodium shows clear and extensive net colonic absorption, and calcium appears to be absorbed to a lesser extent. For potassium, ileal and fecal ranges substantially overlap, while fecal magnesium output can exceed the available ileal influx estimates, making the net colonic handling of these two ions less certain.

The principal remaining uncertainty is therefore not their overall order of magnitude, but variation across individuals and diets and, particularly for potassium and magnesium, the magnitude and direction of net colonic transport. Matched measurements of ileal influx and fecal output within the same individuals under controlled dietary conditions would substantially improve these estimates.

### 6 Host-derived resources entering across the large-intestinal epithelium

#### 6.1 Transport processes through the epithelium of the large intestine

As a second major class of endogenously supplied resources, I consider substrates released into the large intestine across the epithelium. To approach this class of resources more systematically, I first consider the transport capabilities of the large-intestinal epithelium. The major transport processes across the colonic epithelium are well described. Major active and carrier-mediated transport processes are listed in SI Text Table 1 and include the exchange of ions (sodium, potassium, chloride, and bicarbonate) important in electrolyte homeostasis and pH control. Other processes are short-chain fatty acid uptake, urea export, and sulfate exchange. A smaller group of micronutrient- and peptide-specific transporters (biotin/pantothenate, folate, di-/tripeptides) have also been reported.

Besides these active processes, passive transport also contributes to epithelial exchange of small, membrane-permeable molecules. This includes ammonia, dissolved gases (oxygen, carbon dioxide, hydrogen sulfide), and the protonated fraction of short-chain fatty acids (SI Text Table 2). In the following, I focus on epithelial processes that provide quantitatively substantial or metabolically important inputs into the large intestine. I also consider the passive transport of oxygen in Section 7.1 because of its importance in shaping microbial metabolic strategies.

| Transporter | Gene | Substrate | Direction | Reference |
|---|---|---|---|---|
| ENaC | SCNN1A/B/G | $Na^+$ | Apical absorption | [110] |
| NHE2, NHE3 | SLC9A2/3 | $Na^+/H^+$ | Apical | [110] |
| NKCC1 | SLC12A2 | $Na^+/K^+/2Cl^-$ | Basolateral | [110] |
| CFTR | CFTR | $Cl^-$ ($HCO_3^-$) | Apical secretion | [110] |
| DRA | SLC26A3 | $Cl^-/HCO_3^-$; sulfate/oxalate | Apical; sulfate secretion demonstrated in mouse cecum | [111] |
| Colonic H,K-ATPase | ATP12A | $K^+/H^+$ | Apical | [112] |
| BK channel | KCNMA1 | $K^+$ | Apical secretion | [112] |
| MCT1, MCT4 | SLC16A1/3 | SCFA, lactate | $H^+$-coupled | [113] |
| SMCT1, SMCT2 | SLC5A8/12 | SCFA | $Na^+$-coupled, apical | [113] |
| UT-B (±UT-A) | SLC14A1 | Urea | Apical; net host→lumen | [114] |
| SMVT | SLC5A6 | Biotin, pantothenate | $Na^+$-coupled, apical | [115] |
| PCFT / RFC1 | SLC46A1/SLC19A1 | Folate | Apical | [116] |
| PEPT1 | SLC15A1 | Di/tripeptides | Inducible (inflammation, butyrate) | [117] |

**SI Text Table 1.** Major active transport processes across the colonic epithelium.

#### 6.2 Urea secretion and hydrolysis along the large intestine

**Strong influx of urea into the large intestine** A substantial fraction of the urea produced by the liver is not released into urine but disposed of into the gut. This notion goes back to early isotope tracer studies establishing the use of isotope dilution kinetics to show that endogenous urea production consistently exceeded urinary urea excretion, implying a substantial non-renal disposal route [118]. Subsequent work showed that antibiotic treatment reduces the discrepancy between production and excretion via urine, suggesting that the gap in urea disposal is mediated by the bacteria in the gut. The fraction of urea production hydrolyzed by gut bacteria varies considerably with diet: across the wider isotope-kinetic literature, reported hydrolyzed fractions span 20–70%, with compartmental model predictions falling within a narrower 25–60% range [119, Table 5]. While these studies cannot localize urea hydrolysis along the gut more directly, hydrolysis is thought to occur mostly within the large intestine given that most bacteria reside there and that urea and ammonium fluxes from the small to the large intestine are low in comparison (Section 3.4).

**Urea hydrolysis and nitrogen recycling back to the host** Studies with doubly-labeled ([$^{15}N^{15}N$]) urea tracers have further estimated the recycling of released urea by tracking if unhydrolyzed urea is reabsorbed into the body (retains both $^{15}N$ labels) or hydrolyzed to ammonia in the gut, reabsorbed, and resynthesized by the liver (singly labeled urea [$^{14}N^{15}N$] once mixed with the body's much larger pool of ordinary

| Substance | Mechanism | Notes |
|---|---|---|
| $NH_3$ | Passive, pH-dependent diffusion | Net lumen→blood; bidirectional |
| $O_2$ | Passive radial diffusion | Net flux minimized by colonocyte consumption |
| $H_2S$ | Passive diffusion into colonocyte | Followed by mitochondrial sulfide-oxidizing unit |
| $CO_2$ | Passive, bidirectional | Tied to bicarbonate/pH buffering |
| SCFA (protonated) | Passive diffusion | Supplements MCT/SMCT carrier route |

**SI Text Table 2.** Passive/diffusive exchange processes across the colonic epithelium.

$^{14}$N-ammonia). [120] directly measured urea production and hydrolysis at two protein intakes in the same subjects: 118 mg N/kg/day production, of which 35 mg N/kg/day (29.7%) was hydrolyzed, on a low-protein diet (30 g protein/day); and 209 mg N/kg/day production, of which 83 mg N/kg/day (39.7%) was hydrolyzed, on a reference diet (70 g protein/day) – more than twice the absolute hydrolyzed flux on the higher-protein diet. Using a body weight of 75 kg, these correspond to roughly 5.4 g urea/day hydrolyzed on the low-protein diet and 12.9 g urea/day on the reference diet. I note that production itself substantially exceeded dietary protein-nitrogen intake at both diets (121% of intake at a comparable 74 g protein/day in an independent study by the same group [121]), consistent with urea production drawing on both dietary nitrogen and continuous endogenous protein turnover, and, in part, on urea nitrogen previously hydrolyzed, reabsorbed, and resynthesized. Furthermore, only about 15% of the hydrolyzed urea nitrogen was recovered through this recycling route, indicating that hydrolysis does not simply result in quantitative return of the released nitrogen to the hepatic urea pool. However, this number reflects whole-body nitrogen kinetics measured via blood and urine sampling and therefore cannot on its own distinguish bacterial uptake from other fates of the hydrolyzed nitrogen.

[119] extended these findings using a postprandial, non-steady-state compartmental model combined with different intrinsically $^{15}$N-labeled proteins in a meal to address how protein source shapes urea kinetics after a meal. The following numbers were obtained from an 8-hour postprandial window rather than for a full daily rate: total urea nitrogen hydrolysis relative to production ranged from about 24% (milk protein) to 52% (soy) to 58% (wheat) over the 8-hour postprandial window, demonstrating that plant protein sources drive substantially higher hydrolysis efficiency than animal protein. Milk protein's 24% falls close to the low-protein-diet fraction measured directly by Meakins & Jackson (29.7%), while soy and wheat both exceed the reference-diet fraction (39.7%) measured in that same study, with wheat in particular substantially above the range directly measured by either study. In absolute terms, total postprandial urea hydrolysis averaged 5.8 g urea over the 8-hour period across all protein-source groups, with an estimated urea salvage of at least 4.7 g urea over the same window, roughly half of the initial 4.3 g dietary nitrogen load provided in the test meal. Protein intake also modulated hydrolysis efficiency independently of protein source, with a higher protein diet reducing the hydrolysis-to-production ratio from 52% to 35%.

**High ammonium in feces** Directly measured ammonia concentrations in feces further confirm a substantial ammonium loss with a strong dependence on diet. Specifically, fecal ammonium roughly doubled, from 12 to 24 mmol/kg, when dietary protein intake was tripled from 68 to 192 g/day [122]. Notably, fecal ammonia concentrations also fell, from 397 to 278 $\mu$g/g wet weight (approximately 23 to 16 mmol/kg wet weight), when resistant starch intake was increased from 5 to 39 g/day [123] – though whether this reflects reduced net ammonium production, increased bacterial uptake, or simple dilution by the increased fecal mass that typically accompanies higher resistant starch intake cannot be distinguished from concentration data alone. Together with the isotope-labeling-based measurements above, these observations confirm that a substantial pool of ammonium is present in the gut lumen under normal dietary conditions, with its magnitude moving in the expected direction with dietary protein intake, and a meaningful fraction of it escaping host and bacterial capture to be measurable in feces.

**Bacterial utilization of urea** The human studies cited above measure whole-body urea kinetics via blood and urine sampling, and can therefore show that hydrolyzed urea-nitrogen leaves the urea pool, but not whether it is subsequently taken up by the host, by gut bacteria, or both. In vivo isotope infusion in mice closes this gap more directly: circulating urea and ammonia, but not circulating amino acids, were shown to penetrate into feces and become incorporated into genus-resolved bacterial protein, demonstrating genuine uptake into bacterial biomass rather than mere presence in the fecal matrix [124]. Critically, this incorporation was abolished in gnotobiotic mice colonized only with urease-negative bacteria, causally demonstrating that the pathway requires bacterial urease specifically – a level of mechanistic proof not achievable with human fecal-labeling data alone. While this result is in mice and cannot be directly quantitatively transposed to the human g urea/day figures above, it provides the clearest available evidence that hydrolyzed, host-derived urea nitrogen becomes genuine bacterial growth substrate, not merely reabsorbed back into host metabolism.

**Modeling urea influx into the large intestine** The main growth-limitation analysis (Figures 3–5) treats epithelial urea-nitrogen secretion as a population-wide constant, using the central value in Table S1 (the midpoint of the two directly measured conditions described below) independent of an individual's own diet. Here I additionally develop a diet-dependent alternative, in which each NHANES individual's urea contribution instead scales with their own reported dietary protein intake; I use this diet-dependent model exclusively as a robustness check (Figure S4A–D), to confirm that the simpler population-wide constant used in the main analysis does not change which resource is identified as limiting. I model urea production as a function of dietary protein intake, following the well-established relationship that urea is the principal disposal route for surplus dietary amino acid nitrogen. I restrict this model to the range directly supported by isotope-kinetic measurement, rather than extrapolating to zero protein intake. Direct isotope-kinetic measurements using prime/intermittent [$^{15}$N$^{15}$N]urea dilution kinetics – the same isotope method underlying the diet-dependent hydrolysis estimates above – report urea production at both a low-protein (30 g protein/day) and a higher-protein (70 g protein/day) intake in healthy adults [120]. These measurements provide two genuine experimental anchor points rather than a single point requiring extrapolation.

Using the Meakins & Jackson data (30 g protein/day: 118 mg N/kg/day; 70 g protein/day: 209 mg N/kg/day), converted to urea for a 75 kg adult, I fit a linear interpolation between these two measured points. For each NHANES individual $i$, reusing dietary protein intake $\mathrm{Prot}_i$ already defined in Section 2.2,

$$U_{\mathrm{prod},i}(\mathrm{Prot}_i) \approx 8.1 + 0.365\,\mathrm{Prot}_i \text{ g urea/day}, \qquad 30 \leq \mathrm{Prot}_i \leq 70 \text{ g protein/day}.$$

I emphasize that this relationship is an interpolation only, valid strictly within the measured range. I do not extrapolate it to protein intakes below 30 g/day, including to zero (protein-free) intake, for two reasons. First, no isotope-kinetic urea production measurements exist at intakes this low; the lowest directly measured point available to me is 30 g protein/day. Second, direct evidence suggests the production-intake relationship may not remain linear as intake is further reduced: Danielsen & Jackson's own comparison of their 30 g/day result against an earlier study at 35 g/day found a further, disproportionate decline in both production and hydrolyzed fraction between these two nearby low intakes, suggesting the relationship steepens or otherwise changes character near the limit of metabolic adaptation to low protein intake, rather than continuing smoothly to a stable floor.

To estimate the urea flux into the large intestine, I apply a constant hydrolyzed fraction of 30% to this predicted production, giving the diet-dependent mapping

$$J^{\mathrm{LI}}_{\text{urea-secreted},i} = 0.30 \times U_{\mathrm{prod},i}(\mathrm{Prot}_i) \ \text{ g urea/day},$$

converted to nitrogen via the 46.7% factor in Section 9.2. This value is close to the directly measured hydrolyzed fraction at the low-protein diet (29.7%, see above), and I therefore expect the model to perform best near the lower end of the measured intake range. However, the directly measured fraction at the reference diet was 39.7%, meaningfully above the assumed constant value; a fixed 30% fraction therefore likely underestimates hydrolyzed urea flux, and hence the ammonium available to bacteria, at higher protein intakes. I explicitly do not attempt to model a diet-dependence of the hydrolyzed fraction itself: the two directly measured data points above indicate the fraction does increase with protein intake within this study, but the Fouillet et al. postprandial data, when compared across protein sources rather than protein levels, do not show a consistent monotonic pattern, and I consider the available data too limited to support a general, quantitative diet-dependent function for this fraction.

Finally, to probe a potential regime of nitrogen-limited bacterial growth in the gut, I also model a constant lower bound of urea influx, anchored to the lowest protein intake for which direct isotope-kinetic measurement exists (30 g protein/day) rather than to an unmeasured, extrapolated zero-intake floor:

$$J^{\min}_{\text{urea,LI}} = 0.30 \times U_{\mathrm{prod},i}(30) \approx 5.7 \ \text{ g urea/day}.$$

This value is close to, and consistent with, the directly measured hydrolyzed flux at the same diet (5.4 g urea/day, see above). I note this should itself be regarded as a conservative rather than absolute lower bound: protein intakes below 30 g/day certainly occur in practice, but the available data cannot currently constrain urea production reliably in that range.

**Summary and confidence classification** Overall, the urea flux into the large intestine can be estimated with good confidence. Many different systematic and independent isotope labeling studies using several distinct isotope-tracer methodologies confirm that the influx is strong. Directly measured hydrolyzed fractions range from 24% to 58% across the studies discussed above [120, 119], consistent with the wider literature range of 20–70% [119, Table 5]. In absolute terms, directly measured hydrolyzed urea flux ranges from 5.4 g/day (low-protein diet) to 12.9 g/day (reference diet) [120]. Studies further consistently show limited recycling and high ammonium in feces, supporting strong urea turnover in the gut and suggesting that ammonium is amply available for bacterial growth. The diet-dependence of urea influx is qualitatively well established and, within the 30–70 g protein/day range, supported by direct interpolation between measured points; it is not well constrained outside this range, and the dependence of the hydrolyzed fraction itself on diet is not systematically quantified. Because $J^{\mathrm{LI}}_{\text{urea-secreted},i}$ already represents hydrolyzed, ammonia-form nitrogen (Section 6.2), I treat it as essentially fully microbially accessible (utilizable fraction 100%, Table S3) rather than applying a further discount for host reabsorption: whole-body recycling of the hydrolyzed nitrogen back to the host is modest (only about 15%, see above), and this residual recycling competes with, rather than precedes, bacterial uptake – the mouse isotope-infusion evidence above demonstrates genuine incorporation of hydrolyzed urea nitrogen into bacterial biomass, so I do not expect host reabsorption to substantially deplete the pool actually available to gut bacteria.

#### 6.3 Bicarbonate and carbon dioxide turnover along the large intestine

Bicarbonate represents a somewhat unusual resource in the large intestine. Unlike fermentation products such as acetate or gases such as $H_2$, which are predominantly generated by microbial metabolism within the lumen, bicarbonate is actively supplied by the host. Colonic epithelial cells secrete $HCO_3^-$ into the lumen, where it contributes importantly to neutralizing the acids generated during microbial fermentation and thereby to maintaining the luminal pH within a range permissive for microbial growth [125]. Thus, although bicarbonate does not primarily serve as a biomass-building nutrient or energy substrate, its host-derived influx constitutes an important environmental resource for the microbiota by providing buffering capacity.

**Active bicarbonate secretion and coupling to SCFA uptake** Bicarbonate secretion by the colonic epithelium is closely coupled to the absorption of short-chain fatty acids (SCFAs), although the underlying transport physiology involves several pathways. Classical human rectal measurements demonstrated substantial bicarbonate appearance in the lumen during SCFA absorption. McNeil and colleagues found that the excess of SCFA absorption over sodium absorption was approximately matched by bicarbonate appearance [126]. In subsequent experiments, bicarbonate concentrations reached approximately 35–37 mM in rectal dialysis fluid during SCFA absorption, with similar bicarbonate appearance when chloride was replaced by sulfate, supporting secretion of bicarbonate in association with absorbed SCFAs [127].

Mechanistically, an apical $\mathrm{SCFA}^-/\mathrm{HCO}_3^-$ exchange process has been demonstrated in human colonic membrane vesicles. Harig et al. found bicarbonate-stimulated butyrate uptake with an apparent $K_m \approx 1.5$ mM and concluded that $\mathrm{SCFA}/\mathrm{HCO}_3^-$ exchange can mediate butyrate uptake across the apical membrane [128]. However, the relationship between SCFA absorption and bicarbonate secretion in vivo is not described by a single fixed stoichiometry. SCFAs can be transported through multiple pathways, and SCFA uptake can additionally stimulate bicarbonate secretion indirectly through epithelial acid–base regulation. Consequently, although my previous modeling work approximated bicarbonate release as a fixed fraction of SCFA uptake [125], the available experimental data do not justify assigning a universal coupling coefficient between SCFA absorption and bicarbonate secretion.

**Interconversion of bicarbonate and carbon dioxide** Quantifying bicarbonate influx is further complicated by the rapid interconversion of bicarbonate, carbonic acid, and dissolved $CO_2$,

$$CO_2 + H_2O \rightleftharpoons H_2CO_3 \rightleftharpoons H^+ + HCO_3^-.$$

Thus, bicarbonate secreted by the host does not remain as a chemically distinct pool. Its abundance depends on luminal pH, while $CO_2$ generated within the lumen contributes to the same carbonate pool. This makes concentration measurements alone insufficient to reconstruct the gross host-derived bicarbonate flux.

The situation is particularly complicated because $CO_2$ has both endogenous and microbial sources and can exchange across the epithelium. Bacterial fermentation generates substantial $CO_2$ within the large intestine, so in contrast to bicarbonate secretion, much of the luminal $CO_2$ pool is produced locally rather than supplied as an external resource. At the same time, dissolved $CO_2$ can diffuse across the colonic epithelium in either direction. The direction and magnitude of net $CO_2$ exchange therefore depend on the local difference between luminal and tissue $CO_2$ concentrations and can change with microbial fermentation activity [125]. For this reason, a net "carbonate influx" cannot simply be inferred from bicarbonate and $CO_2$ concentrations: host bicarbonate secretion, microbial $CO_2$ production, acid–base interconversion, epithelial $CO_2$ exchange, and luminal gas loss all contribute simultaneously to the measured carbonate pool.

**Role in luminal buffering** Despite the difficulty of assigning a daily flux, the physiological role of host-supplied bicarbonate is well established. Bicarbonate secretion increases the capacity of the lumen to buffer fermentation-derived acids and thereby contributes to regulation of luminal pH. Importantly, bicarbonate supply is physiologically coupled to the metabolic activity of the microbiota: increased fermentation generates more SCFAs, while SCFA absorption is itself associated with bicarbonate secretion. In my previous quantitative model of colonic pH dynamics, this coupling provided a negative feedback in which increased fermentation and SCFA production promoted bicarbonate release, which in turn buffered the associated acidification [125]. Thus, bicarbonate supply is not simply a fixed background influx but can respond to microbial fermentation activity.

**Order-of-magnitude estimate** While the available data do not allow a well-constrained quantitative reconstruction of bicarbonate secretion, they provide information about its likely scale. Human rectal experiments directly demonstrate substantial bicarbonate appearance during SCFA absorption, with bicarbonate appearance being of similar molar magnitude to the imbalance between SCFA and sodium absorption under the tested conditions [126, 127]. Independent estimates of total microbial fermentation-product production further indicate that the underlying metabolic flux is very large. Arnoldini et al. estimate a production of approximately 470 mmol/day of fermentation products for a reference Western diet and show that most of the carbon contained in these products is ultimately absorbed by the host [5].

These observations do not establish a fixed stoichiometric relationship between total SCFA absorption and bicarbonate secretion. They do, however, indicate that bicarbonate secretion associated directly or indirectly with SCFA absorption could readily constitute a flux of tens to hundreds of millimoles per day. As an intentionally broad order-of-magnitude range, I therefore consider approximately 10–300 mmol/day of host bicarbonate secretion plausible. This range should not be interpreted as a measured physiological interval: its purpose is to indicate the likely magnitude of the flux while explicitly retaining the substantial quantitative uncertainty. Formally, $J^{\mathrm{LI}}_{\text{bicarbonate},i} = c_{\text{bicarbonate}} \approx 10\text{–}300$ mmol/day, a population-wide constant not currently mapped to individual-level dietary variation.

**Summary and confidence classification** Bicarbonate is actively supplied to the large intestine by the host and constitutes an important environmental resource through its role in buffering microbial fermentation. Human physiological measurements directly demonstrate SCFA-associated bicarbonate secretion, while the large magnitude of colonic fermentation-product production and absorption further suggests that the associated bicarbonate flux is unlikely to be small. However, the gross daily bicarbonate influx cannot currently be reconstructed reliably because the coupling between SCFA absorption and bicarbonate secretion is not known at the whole-colon level and because secreted bicarbonate participates in a dynamic carbonate pool involving microbial $CO_2$ production, acid–base interconversion, epithelial $CO_2$ exchange, and gas loss. I therefore classify the quantitative bicarbonate influx as **unknown** confidence. For comparison with other resource influxes, I use a broad working range of 10–300 mmol/day, intended only to capture the plausible order of magnitude rather than a quantitatively established physiological range.

#### 6.4 Mucus release along the large intestine

Beyond mucin reaching the cecum via the ileum (Section 3), mucus is also continuously turned over along the length of the large intestine itself, as the epithelium replaces mucin lost to bacterial degradation and luminal shedding. No direct measurement of this local release rate is available. I therefore construct a rough estimate from three more basic, independently measured quantities: the thickness of the colonic mucus layer, its mucin content, and its turnover rate.

**Mucus layer geometry** The mucus coating the colonic epithelium consists of two structurally distinct layers: a dense, largely bacteria-impenetrable inner layer directly apposed to the epithelium, and a looser, more penetrable outer layer facing the lumen. In mice and rats, the dense layer has been reported at $50\text{–}100\ \mu$m thickness and the loose layer at $100\text{–}700\ \mu$m [129]. I take the upper end of each range, $w^{\text{dense}}_{\text{muc,LI}} = 100\ \mu$m and $w^{\text{loose}}_{\text{muc,LI}} = 700\ \mu$m, as representative values. Approximating the large intestine as a cylindrical tube of length $l_{\text{LI}} = 150$ cm and diameter $d_{\text{LI}} = 4$ cm, and each mucus layer as a thin shell coating its inner surface, gives layer volumes

$$V^{\text{dense}}_{\text{muc,LI}} = \pi\, d_{\text{LI}}\, l_{\text{LI}}\, w^{\text{dense}}_{\text{muc,LI}} \approx 19\ \text{mL}, \qquad V^{\text{loose}}_{\text{muc,LI}} = \pi\, d_{\text{LI}}\, l_{\text{LI}}\, w^{\text{loose}}_{\text{muc,LI}} \approx 132\ \text{mL}.$$

**Mucin content** Mucin constitutes only a small fraction of total mucus layer mass, reported to range between $\alpha_{\text{mucin}} = 0.2\text{–}5\%$ [130]. I take the upper end, $\alpha^{\text{dense}}_{\text{mucin}} = 5\%$, as representative of the dense layer. The loose layer is understood to be substantially diluted relative to the dense layer, with total gastrointestinal mucus volume expanding by a factor of 3–4 across this transition [131]. I therefore take $\alpha^{\text{loose}}_{\text{mucin}} = 5/3\,\% \approx 1.7\%$. Approximating mucus density as that of water ($\rho_{\text{water}} \approx 1$ g/mL), the total mucin mass present in the large intestine's mucus layers at any given time is

$$M_{\text{mucin,LI}} \approx V^{\text{dense}}_{\text{muc,LI}}\, \rho_{\text{water}}\, \alpha^{\text{dense}}_{\text{mucin}} + V^{\text{loose}}_{\text{muc,LI}}\, \rho_{\text{water}}\, \alpha^{\text{loose}}_{\text{mucin}} \approx 3.1\ \text{g}.$$

**Turnover and daily release** The standing mucin mass above is continuously replaced as mucus is shed and degraded, primarily by resident bacteria, and resynthesized by goblet cells. Assuming exponential turnover, the daily mucin replacement flux follows from the standing mucin mass and turnover half-life $t_{\text{half}}$ as

$$J_{\text{mucin,LI}} = \frac{\ln 2}{t_{\text{half}}} M_{\text{mucin,LI}}.$$

Colonic mucus protein turnover has been directly quantified using isotope labeling in conventional mice, with median protein half-lives of approximately 4.5–5.5 days [132]. Taking $t_{\text{half}} \approx 5$ days as a representative value gives an estimated daily mucin release along the large intestine of

$$J_{\text{mucin,LI}} \approx 0.4 \text{ g/day}.$$

Formally, $J^{\text{LI}}_{\text{mucus-local},i} = c_{\text{mucus-local}} = 0.4$ g/day, a population-wide constant independent of $D_i$, not currently modeled as diet-dependent (see below).

**Comparison with small-intestinal release** An independent, upper-bound estimate of mucin released from the small intestine, and entering the large intestine via the ileum, has separately been derived from tubing-assisted characterization of ileal lumen content, assuming all measured hexosamines derive from mucin: $\frac{d}{dt} M^{\text{overest}}_{\text{mucin,SI}} \approx 3$–$4$ g/day [13]. Combined with the local large-intestinal release estimated above, this gives a total mucin release of

$$\frac{d}{dt} M_{\text{mucin}} = \frac{d}{dt} M_{\text{mucin,LI}} + \frac{d}{dt} M^{\text{overest}}_{\text{mucin,SI}} \approx 4.4 \text{ g/day}.$$

I note this combined figure and the local large-intestinal component discussed above are treated separately elsewhere in this work (Section 3), and refer there for further discussion of the small-intestinal contribution specifically.

**Limitations** This estimate should be treated as a rough, first-order proxy rather than a precise flux measurement, for several compounding reasons. First, all three inputs – mucus layer thickness, mucin content, and turnover half-life – are themselves uncertain, each spanning several-fold ranges in the literature; I have consistently taken values toward the upper end of reported ranges at each step, so the compounded result should be read as biased toward the higher end of what these inputs could plausibly support, not a central estimate. Second, and more fundamentally, the turnover half-life is derived entirely from mouse measurements; to my knowledge, no directly analogous human measurement of colonic mucus turnover exists, and mucus turnover plausibly differs between species, and may itself depend strongly on local microbial activity, diet, and inflammatory status – factors not captured by a single fixed half-life. Third, unlike several other resource estimates in this work, this figure cannot be cross-checked against fecal output: because resident bacteria substantially degrade mucin before excretion, fecal mucin-derived material reflects only what escapes bacterial consumption, not total release, and the fraction consumed is itself unknown – making fecal measurements uninformative as an independent check on this particular estimate, in contrast to their use elsewhere in this work (e.g. for urea-derived ammonium).

**Summary and confidence classification** Given the number of compounding assumptions and the reliance on non-human turnover data, I classify this estimate as low confidence: a genuine order-of-magnitude estimate is possible from first physiological principles, but it is only weakly constrained by direct measurement. A direct, human-specific measurement of colonic mucus turnover – for example via isotope labeling analogous to the approaches already used for urea kinetics elsewhere in this work – would be the most direct way to improve this estimate.

#### 6.5 Epithelial cell release from the large intestine

Beyond mucus (Section 6.4), the colonic epithelium itself constitutes a further, continuously renewed source of host-derived material entering the large intestinal lumen. Colonic epithelial cells are among the most rapidly renewing cell populations in the body, and their turnover and release into the lumen is a well-established, if still incompletely quantified, physiological process. No direct measurement of the resulting mass flux specific to the human large intestine is currently available. I therefore construct a rough, order-of-magnitude estimate from more basic geometric and cell-biological quantities, following the same general approach used for mucus release above.

**Background: epithelial renewal and shedding** The colonic epithelium is a single layer of columnar cells, continuously replenished by stem cells residing at the base of the crypts of Lieberkühn. Daughter cells generated in the proliferative crypt compartment migrate upward, differentiate, and are ultimately shed from the colonic surface into the intestinal lumen [133, 134]. Human colonic epithelial turnover is rapid, occurring on a timescale of approximately 3–5 days [133]. Unlike the small intestine, where physiological shedding occurs predominantly at villus tips, the large intestine lacks villi and epithelial cells are instead shed from the colonic surface [134].

Shed cells are extruded directly into the base of the dense, epithelium-adjacent mucus layer (Section 6.4); because this mucus layer is itself continuously secreted and pushed outward as part of its own turnover, shed cells are plausibly carried along with it toward the lumen, rather than needing to independently cross a static barrier. I am not aware of a study directly tracing this specific transit process, and note it here as a plausible mechanistic picture rather than a directly confirmed one.

The scale of this process is illustrated by a comprehensive whole-body census of human cellular turnover [135], which integrated the ubiquity, characteristic mass, and lifespan of all major human cell types. This study found that gut (intestinal and gastric) epithelial cells account for only $\sim$12% of total whole-body cell turnover by cell number, but $\sim$40% by mass, owing to their comparatively large size relative to the numerically dominant blood cells – corresponding to roughly 32 g/day (24–40 g/day) of the body's total estimated cellular mass turnover of $80 \pm 20$ g/day. This whole-gastrointestinal-tract figure is not colon-specific, and, like the estimate I construct below, reflects tissue-level production/shedding mass rather than the mass of intact material ultimately recovered in feces; I discuss it here as a useful point of reference for the overall scale of the phenomenon, and use it as an independent plausibility check on my own colon-specific estimate below.

**Estimating the amount released** To estimate the resulting mass flux for the large intestine specifically, I combine an estimate of total colonic epithelial cell number with an estimate of epithelial turnover time.

*Surface area.* Approximating the large intestine as a cylindrical tube of length $l_{\mathrm{LI}} = 150$ cm and diameter $d_{\mathrm{LI}} = 4$ cm gives an unamplified surface area of

$$A_{\mathrm{tube}} = \pi\, d_{\mathrm{LI}}\, l_{\mathrm{LI}} \approx 0.19 \text{ m}^2.$$

This substantially underestimates true epithelial surface area, since it does not account for the additional area contributed by crypt invaginations and haustral folding.

Using instead the estimate of total human large-intestinal mucosal surface area of approximately 1.9–2 m$^2$ [136] gives, for a 70 kg adult,

$$A_{\mathrm{colon}} \approx 2.1 \text{ m}^2.$$

The approximately ten-fold difference from the simple cylindrical estimate illustrates the substantial surface-area amplification produced by crypt architecture and mucosal folding. Because the unamplified cylindrical area does not represent the actual epithelial surface available for cell turnover, I use the mucosal surface-area estimate of approximately 2 m$^2$ for the calculation below.

*Cell density and total cell number.* Approximating a colonocyte's apical footprint as a $10 \times 10$ $\mu$m square, typical for columnar epithelial cells, gives a cell density of

$$\rho_{\mathrm{cell}} \approx 10^{10} \text{ cells/m}^2.$$

Combined with the surface area estimates above, this gives a total colonic epithelial cell number of

$$N_{\mathrm{total}} \approx 2 \times 10^{10} \text{ cells.}$$

*Turnover time.* Human colonic epithelial renewal occurs on a timescale of approximately 3–5 days [133]. I therefore use $\tau = 3$–$5$ days as the characteristic cell-renewal timescale. This range is independently consistent with stable-isotope measurements of intestinal epithelial protein turnover in conventionally raised mice, which report protein half-lives of approximately 3.5–4.2 days across intestinal segments [132]. Because protein turnover is not identical to whole-cell turnover, I use the human epithelial-renewal estimate rather than the mouse protein half-life for the calculation below.

Assuming steady-state renewal, the daily number of cells shed follows as

$$\frac{dN}{dt} = \frac{N_{\mathrm{total}}}{\tau} \approx 4 \times 10^9\text{–}7 \times 10^9 \text{ cells/day.}$$

*Conversion to mass.* Approximating cell volume as the apical footprint times a typical columnar cell height of $\sim$20 $\mu$m gives $V_{\mathrm{cell}} \approx 2 \times 10^{-9}$ cm$^3$, and, assuming a cell density close to that of water, a wet mass of $m_{\mathrm{cell}} \approx 2$ ng per cell. This gives a total shed wet mass of

$$\frac{dM^{\mathrm{wet}}}{dt} \approx 8\text{–}14 \text{ g/day,}$$

or, applying a typical cellular dry-matter fraction of $\sim$20–25%,

$$\frac{dM^{\mathrm{dry}}}{dt} \approx 1.5\text{–}3.5 \text{ g/day.}$$

**Consistency check** This range is broadly consistent with, and substantially smaller than, total fecal dry weight (~30 g/day, of which ~55% is bacterial biomass [5]), leaving ample room for this contribution alongside other known fecal dry-matter components. It is also consistent with, and a small fraction of, the ~32 g/day whole-gastrointestinal-tract epithelial turnover estimate discussed above [135], as expected given that estimate spans the stomach and full length of the small intestine in addition to the colon. I note both comparisons as useful plausibility checks rather than independent validation: as with mucin (Section 6.4), the fraction of shed epithelial material that survives bacterial degradation to appear intact in feces is unknown, so fecal measurements cannot directly confirm the shedding rate itself, only its broad order of magnitude.

**Limitations** This estimate rests on several uncertain geometric and cell-biological assumptions, particularly the representative colonocyte footprint and height and the conversion from epithelial renewal time to luminal mass release. Although human measurements constrain the approximate mucosal surface area and epithelial renewal timescale, I am not aware of a direct, colon-specific human measurement of epithelial mass shedding into the lumen. Moreover, shed cells may undergo rapid degradation within the mucus layer, so tissue turnover need not translate quantitatively into intact cellular material available to luminal bacteria.

**Summary and confidence classification** I therefore classify this estimate as **low** confidence and report it strictly as an order-of-magnitude range, $\frac{dM}{dt}^{\text{dry}} \approx 1.5\text{–}3.5$ g/day, rather than a precise flux. Formally, $J^{\text{LI}}_{\text{epithelial},i} = c_{\text{epithelial}} \approx 1.5\text{–}3.5$ g/day, a population-wide constant range independent of $D_i$. A direct measurement of human colonic epithelial shedding, for example using isotope-labeling approaches capable of following epithelial material into the lumen, would substantially improve this estimate.

#### 6.6 Substrates with net uptake

There are also substrates known or thought to be predominantly taken up by the epithelium, with a net flux directed from the large intestine into the body rather than the reverse. The most important category is short-chain fatty acids and other fermentation products. Strictly speaking, these substrates are not resources supplied to the microbiota; their uptake nonetheless represents an important boundary condition shaping the chemical environment bacteria encounter (luminal pH, nutrient concentration gradients, standing metabolite pools), and I discuss it here for completeness.

**Short-chain fatty acids and other fermentation products** Fermentation products, including lactate and the short-chain fatty acids acetate, butyrate, and propionate, are produced in large quantities by fermenting gut bacteria, driven primarily by catabolic carbohydrate utilization. We recently systematically analyzed the fermentation behavior of major gut bacteria and the resulting fermentation product flux, dissecting its dependence on diet and microbiome composition [5]. Importantly, the large majority of fermentation products released, over 90%, is taken up by the host. This uptake occurs predominantly via carrier-mediated transport (MCT1/MCT4, SMCT1/2), much of it coupled to bicarbonate exchange (Section 6.3), with a smaller passive-diffusion contribution from the protonated, uncharged fraction of SCFA present at physiological pH. Uptake capacity is substantial relative to typical luminal concentrations: measured maximal uptake rates are on the order of 20 mmol m$^{-2}$ h$^{-1}$, with a half-saturation concentration around 20 mM [125]. The efficiency of uptake is spatially coupled to fermentation activity: the proximal colon, where carbohydrate availability and fermentation rates are highest, also shows the most acidic luminal pH and the highest SCFA concentrations (commonly 120–540 mM [125]), favoring both a larger protonated (passively diffusible) fraction and saturating carrier-mediated uptake; distally, as fermentable substrate and SCFA concentrations fall, transporter saturation decreases and the relative importance of transport and diffusive limitations changes. Total daily SCFA production is commonly estimated at several hundred mmol/day, with reported contributions to host daily energy supply on the order of several percent with the exact contribution depending strongly on diet [126, 5].

**Ammonium** As discussed in Section 6.2, ammonium is released in substantial quantities in the large intestine via bacterial hydrolysis of host-supplied urea. This produces substantial luminal ammonium concentrations, confirmed directly by fecal measurements (on the order of 15–55 mmol/kg wet feces depending on diet; Section 6.2). Blood ammonia concentrations are, by contrast, very low (on the order of 15–50 $\mu$M), orders of magnitude below the concentrations measured in fecal material. Given that ammonia diffuses readily across the colonic epithelium, this gradient drives substantial net host uptake. Direct evidence for this uptake comes from both the excluded-colon perfusion literature (rapid, near-complete equilibration of luminal and blood ammonia within $\sim$60 minutes) and the isotope-tracer urea studies discussed above, which quantified that a meaningful share of hydrolyzed urea-nitrogen is recovered as host-reabsorbed, resynthesized urea in urine, direct confirmation that hydrolyzed ammonia is absorbed by the host rather than remaining entirely luminal (Section 6.2).

**Sulfate** Unlike SCFA and ammonium, the direction of net sulfate movement across the human colonic epithelium is not well established. Human mass-balance data are consistent with substantial net removal of sulfate somewhere along the colon: an estimated 1–12 mmol/day reaches the large intestine (Section 7.3), while free faecal sulfate loss is trivial ($<$0.5 mmol/day). However, this removal cannot currently be attributed specifically to host absorption, since bacterial reduction of sulfate to hydrogen sulfide by sulfate-reducing bacteria offers an equally plausible microbial fate for the missing sulfate, and the two are not distinguished by mass balance alone. Complicating the mechanistic picture further, a study directly measuring transepithelial sulfate flux in isolated mouse cecum found net *secretion* of sulfate via the anion exchanger DRA (SLC26A3), alongside simultaneous chloride absorption by the same transporter [111] – in apparent tension with earlier human and rabbit colonic perfusion data more consistent with net absorption. Given this mouse-specific, cecum-specific, and apparently contested finding, I do not treat sulfate as a clear example of net host uptake here, and flag its transport direction as a genuinely open question rather than resolving it in either direction.

**Other substrates and notable non-examples** Bile acids are also substantially reabsorbed along the large intestine, though I treat this separately in Section 3.3 given the added complexity of ongoing microbial deconjugation and transformation. Water and the major inorganic ions ($Na^+$, $K^+$) are absorbed extensively along the colon (see Section 5). Notably, free amino acids are not thought to undergo substantial net absorption in the adult human colon, in marked contrast to SCFA and ammonium [137].

### 7 Terminal electron acceptors

In addition to serving as substrates for biomass synthesis and fermentation, resources entering the large intestine can support respiratory energy metabolism by acting as terminal electron acceptors. Gut bacteria encode different aerobic and anaerobic respiratory pathways, using electron acceptors including oxygen, nitrate, sulfate and other sulfur compounds, fumarate, trimethylamine N-oxide (TMAO), dimethyl sulfoxide (DMSO), and numerous additional inorganic and organic compounds [138, 46, 139]. Because respiratory energy generation can substantially increase the energetic yield obtained from organic substrates, even comparatively small electron-acceptor influxes can have important consequences for microbial growth and competition.

I focus on oxygen, nitrate/nitrite, and sulfate because they represent the major terminal electron acceptors for which physiologically relevant delivery to the human large intestine can be quantitatively constrained from dietary and physiological measurements. Other electron acceptors used by particular gut bacteria, including fumarate, TMAO, and DMSO, may be ecologically important in specific contexts but currently lack sufficiently constrained human influx measurements for inclusion in the quantitative reconstruction; I discuss them briefly below for completeness. Electron acceptors generated primarily during inflammation or produced locally by microbial metabolism are not included as independent resource influxes in my reference reconstruction of the healthy adult large intestine.

I consider the influx of these electron acceptors as a distinct resource category because their influxes are typically much smaller than those of major fermentable substrates and arise from both dietary and endogenous sources.

#### 7.1 Oxygen

Among the electron acceptors considered here, oxygen provides the highest redox potential and therefore supports the greatest energetic yield from respiration. However, its quantitative influx into the healthy human cecum is poorly constrained. Here I present a flux estimation based on a systematic literature review and the available measurements (summarized in SI Text Table 3), together with an explicit account of the uncertainties involved.

**Luminal oxygen concentrations** Several capsule-based ingestible sensors have reported oxygen-equivalent gas signals or oxidation-reduction potential along the human gastrointestinal tract [140, 141]. However, the metal-oxide sensor used by Kalantar-Zadeh et al. provides only relative signals and responds to oxidizing species more generally. The newer GISMO platform measures oxidation-reduction potential rather than oxygen itself and resolves only three broad intestinal compartments. Thus, while these capsule measurements do not provide absolute cecal oxygen partial pressures, they consistently demonstrate a sharp transition from the more oxidizing small intestine to the strongly reducing large intestine. This is compatible with oxygen being delivered from the distal small intestine, while direct rectal measurements confirm that the distal large-intestinal lumen itself is extremely anoxic, with luminal $pO_2$ below 0.1% ($< 0.13$ kPa) [142].

To obtain an (upper bound) estimate of oxygen influx, I base my primary estimate on direct human tissue electrode measurements. Lampe et al. (as compiled in [143]) report serosal $pO_2$ of 34 mmHg ($\sim$4.5%, $\sim$4.5 kPa) at the terminal ileum and 30 mmHg ($\sim$3.95%, $\sim$4.0 kPa) at the cecum. I use these direct human tissue measurements as a deliberately generous reference oxygen tension for constructing an upper bound on luminal oxygen delivery. Because the measurements are serosal rather than luminal, they do not establish the oxygen partial pressure of ileal contents directly. Applying the tissue-side value to the luminal fluid or gas therefore biases the resulting estimate upward and should be interpreted as an upper-bound assumption. In the following I use this as my reference value for ileal oxygen. As my measurements are tissue-proximal rather than luminal, and oxygen is expected to diffuse from tissue into the lumen rather than the reverse, I treat my estimate as an upper bound of the luminal gas-phase $O_2$.

| Source | Method | Species | Region | Key value |
|---|---|---|---|---|
| Lampe et al., in [143] | Electrode | Human | Ileum, cecum, sigmoid (serosa) | Ileum 4.5%, cecum 4.0%, sigmoid 5.1% |
| Lind Due et al. 2003 [142] | Polarographic electrode, direct in vivo | Human | Rectal lumen | $<$0.13% |
| Fisher et al. 2013 | EPR oximetry (implanted particulate probe) | Rat | Small intestine, serosal wall (not lumen) | 7.2% (54.5 mmHg) |
| Albenberg et al. 2014 | Phosphorescence quenching (luminal probe) | Mouse | Cecum lumen | $<$0.13% |
| Friedman et al. 2018 | Oxyphor MicroS (lumen) / G4 (tissue) | Mouse (conv. & germ-free) | Terminal ileum, cecum | Lumen near zero in both conventional and germ-free |
| Kalantar-Zadeh et al. 2018 [140] | Non-specific MOS gas sensor ("$O_2$-equivalent") | Human, direct, whole tract | Stomach, jejunum, ileum, colon | Not used as basis; see text |
| Even et al. 2025 [141] | Electrochemical ORP sensor | Human, direct, whole tract | Stomach, small intestine, large intestine (3 compartments only) | Not used quantitatively; qualitative corroboration only |

**SI Text Table 3.** Summary of oxygen-tension measurements in the distal small intestine and cecum/colon, by method and species. The Fisher et al. measurement is from rat small-intestinal *serosal wall* tissue rather than the lumen.

**Estimating oxygen supply via ileal fluid** With the estimate of oxygen levels in the terminal ileum, $p_{O_2} = 34$ mmHg, I next estimate how much of oxygen could plausibly reach the cecum dissolved in the fluid that passes through the ileocecal valve each day. Assuming the gas phase to be in equilibrium with the liquid phase, I calculate the concentration of dissolved oxygen in the luminal fluid,

$$C_{O_2} = \alpha\, p_{O_2} = 1.02 \text{ mL/L}.$$

Here $\alpha = 0.003$ mL $O_2$ dL$^{-1}$mmHg$^{-1}$ is the oxygen solubility coefficient of Henry's law for aqueous solution at a temperature of 37°C. For a daily ileocecal fluid flow of approximately $Q_{cecum} = 2$ L/day, this dissolved oxygen concentration translates into a total daily oxygen flux of

$$J_{O_2} = C_{O_2} \times Q_{cecum} \approx 2.04 \text{ mL/day} \approx 91 \ \mu\text{mol/day} \approx 2.9 \text{ mg/day}.$$

**Estimating oxygen supply via ileal gas** Importantly, this is only a starting point as besides oxygen influx via the luminal fluids, oxygen can also enter the cecum in the gas phase. I am not aware of any direct measurements quantifying gas flux from the small to the large intestine. To roughly estimate the oxygen influx, I thus start from the measured gas-to-liquid volume ratios observed within the small intestine (SI Text Table 4), assuming this same ratio approximates the content passing into the cecum and using the directly measured influx of luminal fluids into the cecum, $Q_{cecum} = 2$ L/day. This assumes that the standing gas:liquid volume ratio measured in the small intestine approximates the ratio transported across the ileocecal valve, an assumption for which direct measurements are not available and which therefore represents a major source of uncertainty. Since the gas:liquid ratio in the small intestine differs between fasting and active digestion (SI Text Table 4), I account for this by separating the daily liquid flow into two temporal components: a larger flow occurring during digestion (approximately 1.5 L/day, associated with the fed-state ratio) and a smaller basal flow occurring during fasting periods between meals (the remainder, approximately 0.5 L/day associated with the fasting-state ratio) [144, 145].

Using the fasting gas:liquid ratio ($16.7\,\text{mL} : 43\,\text{mL} \approx 0.39$) and the fed ratio ($16.7\,\text{mL} : 94\,\text{mL} \approx 0.18$), the accompanying gas volumes for each component are

$$V_{\text{gas}}^{\text{fed}} \approx 0.18 \times 1500 \text{ mL} \approx 267 \text{ mL/day}, \qquad V_{\text{gas}}^{\text{fasting}} \approx 0.39 \times 500 \text{ mL} \approx 194 \text{ mL/day},$$

giving a total gas volume of $V_{\text{gas}} \approx 461$ mL/day. At 4.5% $O_2$ in the gas phase, this contributes an additional oxygen flux of

$$J_{O_2}^{\text{gas}} \approx 0.045 \times 461 \approx 20.7 \text{ mL/day}.$$

Adding the dissolved-liquid contribution from the previous section ($J_{O_2}^{\text{liquid}} \approx 2.0$ mL/day) gives a combined estimate of

$$J_{O_2}^{\text{ileal}} \approx 22.7 \text{ mL/day} \approx 1013 \ \mu\text{mol/day} \approx 32.4 \text{ mg/day}.$$

As a sensitivity check, I also compute the two limiting cases in which the entire 2 L/day liquid flow is assumed to occur under a single digestive state throughout the day. Assuming the fed ratio applies to the full 2 L/day gives $J_{O_2} \approx 18.0$ mL/day $\approx 25.7$ mg/day; assuming the fasting ratio applies throughout gives $J_{O_2} \approx 37.0$ mL/day $\approx 52.8$ mg/day. Together with the liquid-only estimate from the previous section as a floor, this gives

$$J_{O_2}^{\text{ileal}} \approx 2.9\text{–}52.8 \text{ mg/day} \approx 0.09\text{–}1.65 \text{ mmol/day}$$

with the combined fed/fasting split above (32.4 mg/day) as my primary estimate within this range.

| Source | State | Value | Notes |
|---|---|---|---|
| Levitt 1971, *NEJM* | Fasting | 30–200 mL total gut gas | Argon washout technique; $O_2 < 2\%$ of gas |
| Perez, Accarino, Azpiroz et al. 2007, *Am J Gastroenterol* | Fasting | $\sim$100 mL total, $\sim$equal across 6 compartments (SI $\approx$ 15–20 mL) | CT volumetric analysis; |
| Perez et al. 2007 (cont.) | Fed | $+65\%$ total gas, almost entirely in *pelvic colon*; SI gas roughly unchanged | Postprandial increment appears at $99 \pm 22$ min |
| MRI water-ingestion studies | Fasting | SI fluid $\approx 43 \pm 14$ mL | Resting small-bowel water |
| MRI water-ingestion studies (cont.) | Fed (post 240 mL water) | SI fluid $\approx 77$–$94$ mL | |

**SI Text Table 4.** Gas and fluid volume data used to construct the fasting vs. fed gas:fluid ratio in the small intestine.

##### 7.1.1 Oxygen delivery by epithelial diffusion

Oxygen can also diffuse across the epithelium into the cecal lumen, independent of ileal influx. This pathway has been discussed frequently in the microbiome literature, highlighting the steep gradients across the mucus layer from oxygenated tissue to an essentially anaerobic lumen [146, 147, 148]. However, the flux of exchanged oxygen is poorly quantified. To obtain an order-of-magnitude estimate of this flux, I use the same human serosal cecal measurement introduced above (30 mmHg; Lampe et al., in [143]) as the tissue-side oxygen concentration driving diffusion into the lumen. I then apply Fick's law to estimate the diffusive flux as

$$J_{O_2} = D_{O_2}\, A\, \frac{\Delta C}{L},$$

with appropriate values or estimations for the diffusion coefficient $D_{O_2}$, the surface area of the cecum $A$, and the concentration gradient $\frac{\Delta C}{L}$ (based on the diffusion distance L and a concentration difference $\Delta C$).

**Diffusion distance** $L$ spans the physical path from the serosal/submucosal source to the lumen. This path consists of three components. First, the mucus layer overlying the epithelium, measured directly in the human right colon at $107\ \mu$m [149]. Second, the epithelium itself, a single layer of columnar cells; I assume a representative thickness of $20$–$25\ \mu$m, a typical value for columnar epithelium rather than a colon-specific measurement, adopted here as a modeling assumption. Third, a subepithelial gap of approximately $20\ \mu$m separating the epithelium from the capillary plexus that supplies it; this figure is likewise an assumed geometric parameter, not a direct measurement, chosen to be generous toward diffusion (i.e. biased toward a shorter rather than longer distance) since it feeds only the upper-bound flux calculation below. Summing these three contributions gives

$$L \approx 107 + 25 + 20 \approx 150\ \mu\text{m}.$$

##### 7.1.2 Surface area, and why I do not apply a crypt amplification factor

The cecum measures approximately 5–6 cm in length. Diameter is more variable, since the cecum is distensible, frequently partially collapsed or moderately filled. I use a functional range of 4–6.5 cm diameter, spanning collapsed to openly patent (non-pathological) states. Modeling the cecum as a cylinder with a closed hemispherical cap:

$$A_{\text{low}} = \pi(4)(5) + \pi(2)^2 \approx 75\ \text{cm}^2, \qquad A_{\text{high}} = \pi(6.5)(6) + \pi(3.25)^2 \approx 156\ \text{cm}^2$$

Unlike the small intestine, the colon has no villi, so its surface area is amplified only by crypts and haustral folding. I deliberately do not apply an amplification factor for this crypt surface as the epithelial oxygen consumption rate used below (Ussing chamber measurement) was reported per unit of mounted/gross tissue area. Instead, I use the unamplified gross surface area, $A \approx 75\text{–}156$ cm$^2$ ($0.0075\text{–}0.0156$ m$^2$), for the diffusion estimates below.

**Flux calculation** The remaining parameter is the oxygen diffusion coefficient, $D_{O_2} = 2.5 \times 10^{-9}$ m$^2$s$^{-1}$, an assumed representative literature value for oxygen diffusivity in aqueous/mucus-like media rather than a colon-specific measurement. The concentration difference $\Delta C$ follows from the tissue-side oxygen level introduced above (30 mmHg), converted via Henry's law ($\alpha \approx 1.3\ \mu$M/mmHg) to $\Delta C \approx 39\ \mu$M, taken relative to an assumed lumen concentration of zero.

With all these parameters — $D_{O_2} = 2.5 \times 10^{-9}$ m$^2$s$^{-1}$, $\Delta C \approx 39\ \mu$M, $L \approx 150\ \mu$m, and $A \approx 75\text{–}156$ cm$^2$ — I obtain

$$J_{O_2}^{\text{passive}} \approx 0.42 \text{ to } 0.88 \text{ mmol } O_2\text{/day}, \quad \text{central estimate} \approx 0.65 \text{ mmol/day}.$$

This represents a maximal, no-consumption upper bound on diffusive oxygen delivery into the cecal lumen.

**Comparison with epithelial oxygen consumption capacity** Ussing-chamber measurements of isolated human sigmoid colonic mucosa report a basal epithelial oxygen consumption rate of $Q_{O_2} = 8.09 \pm 0.55\ \mu$mol h$^{-1}$cm$^{-2}$, supplied preferentially from the serosal rather than the mucosal side, consistent with a "rectifying" asymmetry in transepithelial oxygen transport demonstrated in rat colon. Applying this rate to the same unamplified surface area used above:

$$J_{O_2}^{\text{consumption}} = 8.09\ \mu\text{mol h}^{-1}\text{cm}^{-2} \times 24 \text{ h} \times (75\text{–}156 \text{ cm}^2) \approx 15 \text{ to } 30 \text{ mmol/day}$$

This consumption capacity exceeds the diffusive supply upper bound by roughly $17\text{–}70\times$, providing a consistency check for why the cecal lumen remains near-anoxic despite a substantially higher tissue-side source: the measured epithelial oxygen-consumption capacity is more than sufficient to account for the calculated diffusive supply, consistent with substantial consumption of tissue-derived oxygen before it reaches the bulk lumen.

**Reaction-diffusion check: how much of this flux is plausible** The comparison above establishes that consumption *capacity* exceeds diffusive supply, but not how much of the upper-bound flux would actually survive the 150 $\mu$m transit. I assess this using a reaction-diffusion comparison.

Diffusion timescale across $L = 150\ \mu$m:

$$\tau_{\text{diff}} = \frac{L^2}{D_{O_2}} = \frac{(1.5 \times 10^{-4})^2}{2.5 \times 10^{-9}} \approx 9 \text{ s}$$

Consumption timescale, converting the areal Ussing-chamber rate to a volumetric rate using an approximate mucosal thickness of 400 $\mu$m:

$$R = \frac{8.09\ \mu\text{mol h}^{-1}\text{cm}^{-2}}{400\ \mu\text{m}} \approx 0.056 \text{ mol m}^{-3}\text{s}^{-1}$$

At $C_0 = 39\ \mu$M ($0.039$ mol/m$^3$):

$$\tau_{\text{react}} = \frac{C_0}{R} \approx \frac{0.039}{0.056} \approx 0.7 \text{ s}$$

Damköhler number:

$$\text{Da} = \frac{\tau_{\text{diff}}}{\tau_{\text{react}}} \approx 13$$

indicating a strongly reaction-dominated regime. Steady-state penetration depth:

$$\delta = \sqrt{\frac{2D_{O_2}C_0}{R}} = \sqrt{\frac{2 \times 2.5 \times 10^{-9} \times 0.039}{0.056}} \approx 59\ \mu\text{m}$$

This penetration depth is well below $L = 150\ \mu$m, suggesting oxygen entering at the tissue boundary is largely consumed before reaching the mucus/lumen interface. This calculation is intended only as an order-of-magnitude consistency check, because it treats oxygen consumption as spatially uniform and approximately zero-order over the relevant concentration range. The true diffusive contribution to the bulk lumen is likely substantially smaller than the Fick's law estimate above.

I note a caveat: the Ussing-chamber rate was measured across the full $\sim$400 $\mu$m mucosal thickness, including the more metabolically active crypt-base/proliferative zone, and consumption is unlikely to be uniform across this depth. Applying the whole-tissue average rate uniformly to the outer 150 $\mu$m likely overstates local consumption there, so $\delta \approx 59\ \mu$m should be read as a lower bound on the true penetration depth. I do not convert this into a corrected point estimate, since that would require a depth-resolved consumption rate that is not available; I present it as a quantitatively grounded indication that the no-consumption flux is an overestimate.

##### 7.1.3 Overall oxygen influx

$$J_{O_2}^{\text{ileal}} \approx 0.09\text{–}1.65 \text{ mmol/day}, \qquad J_{O_2}^{\text{epithelial}} \lesssim 0.42\text{–}0.88 \text{ mmol/day} \quad \text{(no-consumption upper estimate).}$$

The two contributions are therefore of broadly comparable maximal magnitude, but they are not equally well constrained. The ileal estimate depends strongly on the unmeasured gas flux across the ileocecal valve, whereas the epithelial estimate explicitly neglects oxygen consumption during diffusion and is therefore expected to overestimate delivery to the bulk lumen. The actual relative contribution of these two routes remains unknown.

Ileal oxygen delivery is additionally expected to vary strongly over time with postprandial fluid transfer, whereas epithelial diffusion should be more continuous; these temporal dynamics are not quantified here.

In mice, strong oxygen consumption by the cecal epithelium has been confirmed, leading to an anoxic cecal environment even in germ-free mice lacking microbial oxygen consumption [150]. As such, my epithelial diffusion estimate likely represents an upper bound, and real values could be considerably lower. Mice differ from humans in several important quantities, however, including cecal geometry and frequency of food intake, so the extent to which such a process occurs in the human gut remains unknown.

**Summary and confidence classification** Overall, I classify oxygen influx into the large intestine as **unknown** confidence. No direct measurement of total oxygen delivery to the human cecum is available, requiring the flux to be constrained indirectly from oxygen concentrations in ileal contents, gas transport, and epithelial diffusion. Each of these contributions is itself only partially constrained, with the gas-phase contribution particularly uncertain, and no direct measurement exists against which to validate the combined estimate. Despite this uncertainty, the available constraints establish that oxygen delivery to the healthy large intestine is small in absolute terms, on the order of at most approximately millimoles per day. This places a quantitative upper bound on the extent to which aerobic respiration can contribute to bulk microbial carbon catabolism under healthy conditions. Formally, $J^{\mathrm{LI}}_{O_2,i} = c_{O_2} \approx 0.5\text{–}2.5$ mmol/day (combined ileal and epithelial routes), a population-wide constant independent of $D_i$.

More measurements are needed to go beyond these estimations. Oxygen concentrations in the terminal ileum should be more rigorously measured, ideally via direct sampling of luminal fluid or a properly $O_2$-selective (rather than non-specific or ORP-based) ingestible sensor. Gas exchange across the ileocecal junction should ideally be observed directly, resolved across the postprandial cycle. Depth-resolved epithelial oxygen consumption, and a human equivalent of the germ-free chemical oxygen-consumption assay, would help convert the diffusive upper bound into a genuine flux estimate.

#### 7.2 Nitrate and nitrite

Estimating nitrate and nitrite influx into the cecum is more tractable than estimating oxygen influx because both occur as dissolved solutes and therefore lack the gas:liquid partitioning ambiguity described above. Moreover, direct human measurements of terminal ileal contents are available: several studies have measured nitrate and nitrite concentrations in ileostomy effluent. I combine these measurements with dietary pharmacokinetic data to construct two independent estimates of nitrate delivery to the large intestine.

**Approach 1: dietary intake and pharmacokinetics** Typical dietary nitrate intake in adults is approximately $100\ \mathrm{mg\,day^{-1}}$[151], corresponding to

$$J_{\mathrm{diet}} \approx \frac{100\ \mathrm{mg}}{62\ \mathrm{mg\,mmol^{-1}}} \approx 1.6\ \mathrm{mmol\,day^{-1}},$$

although diets rich in leafy vegetables may exceed $5\ \mathrm{mmol\,day^{-1}}$ [152].

Human pharmacokinetic studies demonstrate that nitrate is rapidly and efficiently absorbed from the small intestine, with the large majority entering the systemic circulation and only a small fraction expected to escape absorption [152, 153]. This is corroborated directly by balance studies: Florin, Neale & Cummings (1990) gave six healthy ileostomy subjects diets varying in nitrate content from 0.83 to 5.20 mmol/d and measured nitrate and nitrite in both ileal effluent and urine by anion-exchange chromatography. They found no significant nitrite in ileal effluent at any intake level, and that dietary nitrate was overwhelmingly recovered in urine (1.31–4.25 mmol/d) rather than ileal effluent, consistent with near-complete small-intestinal absorption [154].

The available pharmacokinetic and balance studies indicate that only a small fraction of dietary nitrate escapes small-intestinal absorption. To establish the corresponding scale, I consider an illustrative escape fraction of 1–5%, consistent with near-complete absorption but not intended as a directly measured population range.

If a fraction $f_{\text{escape}}$ of dietary nitrate escapes absorption, the flux to the cecum is

$$J_{\text{cecum}} = f_{\text{escape}} \, J_{\text{diet}}.$$

yielding

$$J_{\text{cecum}} \approx 0.016\text{–}0.08 \text{ mmol day}^{-1}$$

for a typical Western diet, and for nitrate-rich diets ($J_{\text{diet}} \approx 5 \text{ mmol day}^{-1}$),

$$J_{\text{cecum}} \approx 0.05\text{–}0.25 \text{ mmol day}^{-1}.$$

Dietary intake and pharmacokinetics alone thus suggest an approximate upper bound of

$$J_{\text{cecum}} < 0.1\text{–}0.3 \text{ mmol day}^{-1}.$$

**Approach 2: direct ileostomy effluent measurements** A more direct estimate can be obtained from studies measuring nitrate concentration directly in terminal ileal (ileostomy) effluent. Three such studies give a genuinely inconsistent picture, which I present in full rather than selecting only the most convenient value.

Radcliffe et al. (1989) measured nitrate in ileostomy effluent from healthy ileostomy subjects on conventional and nitrate-rich diets, reporting median concentrations of $6.7$ $\mu$M (conventional diet) and $20.5$ $\mu$M (nitrate-rich diet), with maximum measured concentrations approaching $50$ $\mu$M [155]. This is broadly consistent with a more recent review citing ileal effluent nitrate concentrations of 30–60 $\mu$M [156].

In contrast, Saul et al. (1981), using a method the authors specifically designed to avoid interferences and measurement artifacts present in earlier assays, reported substantially lower ileostomy nitrate concentrations of 0–7 $\mu$mol/kg (comparable in order of magnitude to $\mu$M for an aqueous effluent), together with nitrite at 0–15 $\mu$mol/kg [157]. They also demonstrated that nitrate and nitrite can be altered by microbial activity during sample handling, emphasizing the sensitivity of these measurements to collection and analytical procedures. The authors additionally argued that some discrepancies with earlier reports could reflect assay interference. Separately, the Florin/Cummings balance study described above found no significant ileal effluent nitrite at all, and a nitrate mass balance consistent with only a small fraction of dietary nitrate escaping small-intestinal absorption [154].

Taken together, these three studies do not agree on a single ileal effluent nitrate concentration, spanning roughly 0–60 $\mu$M depending on method and study. I do not attempt to adjudicate which is most accurate; instead, I use the upper end of this range to construct a conservative upper-bound flux estimate. Assuming a terminal ileal flow of $Q_{\text{ileum}} = 1.5\text{–}2 \text{ L day}^{-1}$ [144] and the highest reported concentrations ($\sim$50–60

$\mu$M):

$$J_{\text{cecum}} = Q_{\text{ileum}}[\text{NO}_3^-] \approx 0.075\text{–}0.12\ \text{mmol}\,\text{day}^{-1}.$$

Using instead the lower end of the range (Saul et al., $\lesssim$7 $\mu$mol/kg) gives a substantially smaller flux, on the order of $0.01\text{–}0.014\ \text{mmol}\,\text{day}^{-1}$, and the Florin/Cummings balance data would suggest an even smaller contribution still. I therefore treat

$$J_{\text{cecum}} < 0.1\text{–}0.15\ \text{mmol}\,\text{day}^{-1}$$

as a conservative upper bound from direct ileostomy measurements, noting explicitly that this is anchored to the highest reported concentrations in an inconsistent literature, and that the true value may be considerably lower.

**Interpretation** Despite the disagreement within the ileostomy effluent literature itself, both independent approaches — dietary pharmacokinetics and direct ileal effluent measurement — converge on a broadly consistent picture. I distinguish two figures here: a *typical* influx, of order

$$10^{-2}\text{–}10^{-1}\ \text{mmol}\,\text{day}^{-1}$$

(i.e. approximately 0.01–0.1 mmol/day) under ordinary dietary conditions, and a *conservative upper bound* of $0.1\text{–}0.3\ \text{mmol}\,\text{day}^{-1}$, reflecting nitrate-rich diets and the least favorable (highest-concentration) ileostomy measurements. Taking the full span from the typical lower end to the conservative upper bound gives a working range of approximately $0.01\text{–}0.3\ \text{mmol}\,\text{day}^{-1}$ for this resource, used as such in Table S1. Formally, $J^{\text{LI}}_{\text{nitrate},i} = c_{\text{nitrate}} \approx 0.01\text{–}0.3$ mmol/day, a population-wide constant not currently mapped to individual-level NHANES dietary nitrate intake. This agreement across two methodologically independent approaches is reassuring, even though the ileostomy studies do not agree closely with one another in absolute terms.

These estimates apply only to dietary nitrate reaching the cecum. Additional nitrate may enter the lumen via host-derived nitric oxide metabolism and epithelial secretion, particularly during inflammation, but quantitative estimates for this contribution are currently lacking. Nitrite entering the cecum from the ileum is consistently reported at much lower concentrations than nitrate across all three ileostomy studies discussed above, in some cases below the limit of detection; I therefore do not construct a separate flux estimate for nitrite and treat it as a minor contributor relative to nitrate.

Interestingly, despite the absence of a single reliable direct measurement of cecal nitrate flux, numerous studies demonstrate rapid microbial utilization of nitrate by facultative anaerobes. Consequently, even relatively small continuous nitrate fluxes may exert a disproportionate ecological influence, since nitrate is rapidly consumed and rarely accumulates. As for oxygen, ecological relevance is therefore determined by flux and rapid turnover rather than by standing concentration alone.

**Summary and confidence classification** Overall, nitrate influx into the healthy large intestine can be estimated with moderate confidence. Dietary pharmacokinetics and direct ileostomy measurements independently indicate a typical influx on the order of $10^{-2}\text{–}10^{-1}$ mmol/day, with a conservative upper bound of approximately 0.1–0.3 mmol/day. Nitrite influx appears still smaller. These estimates apply to the healthy state; host-derived nitrate production can increase substantially during inflammation.

#### 7.3 Sulfate

Sulfate is a major anaerobic respiratory electron acceptor for sulfate-reducing bacteria. It is important to consider in the gut, because the influx into the large intestine is comparatively large. I distinguish free sulfate and those bound to mucus.

**Free sulfate** Free sulfate influx into the large intestine can be estimated from a controlled dietary study systematically measuring sulfate in the diet and ileal effluent [158]. Across diets ranging from 1.6–16.6 mmol/day sulfate intake, the capacity of the small intestine to absorb sulfate plateaus at about 5 mmol/day, above which additional intake passes through. Endogenous secretion adds a further 0.96–2.6 mmol/day independent of diet. Together, this gives a total sulfate flux from the ileum into the large intestine of approximately

$$J_{\text{cecum}} \approx 1\text{–}12\ \text{mmol}\,\text{day}^{-1}.$$

Free fecal sulfate loss is much lower ($< 0.5$ mmol/day), indicating extensive removal of free sulfate during colonic transit, through microbial sulfate reduction, host absorption, or both [158].

**Mucin-associated sulfate** The same study measuring free sulfate in ileal effluent also quantified total sulfate, reporting 1–2 mmol bound sulfate loss in ileal effluent per day [158]. The majority of this influx likely reflects sulfate in mucins, in line with the 3–4 g mucin entering the large intestine per day (Section 3.1) which contains about 0.6 to 2.7 mmol sulfate. Unlike free sulfate, however, this organically bound sulfur is not directly available as a terminal electron acceptor and must first be released from mucin glycans by microbial sulfatases.

**Summary and confidence classification** Overall, free sulfate influx into the large intestine can be estimated with moderate confidence from direct human ileal measurements, with a representative range of approximately 1–12 mmol/day depending strongly on dietary sulfate intake. An additional $\sim$1–2 mmol/day of sulfate is delivered in bound forms, particularly mucin, but this pool requires microbial liberation before it can participate in sulfate respiration. Formally, $J^{\text{LI}}_{\text{sulfate},i} = c_{\text{sulfate}} \approx 1\text{–}12$ mmol/day, a population-wide constant not currently mapped to individual-level NHANES dietary sulfate intake.

#### 7.4 Other electron acceptors: fumarate, DMSO, and TMAO

Fumarate, dimethyl sulfoxide (DMSO), and trimethylamine N-oxide (TMAO) are well-established terminal electron acceptors for anaerobic respiration by gut bacteria. Their role is particularly well characterized in facultative anaerobes such as *Escherichia coli* and *Salmonella*, while fumarate respiration is also widespread among abundant obligate anaerobes, including *Bacteroides*. However, these compounds differ from oxygen, nitrate, and free sulfate in that their *external influx* into the healthy large intestine is poorly constrained and their influxes are expected to be comparatively low. I therefore discuss them qualitatively rather than assigning quantitative influxes.

Fumarate can reach millimolar concentrations in intestinal contents [159], but luminal abundance does not translate directly into an external resource influx. Many gut bacteria produce fumarate locally as a metabolic intermediate, and some organisms, including *Bacteroides*, generate fumarate specifically to support their own anaerobic electron-transport chains. Thus, fumarate respiration can be physiologically important even when the net dietary or host-derived fumarate influx is small or unknown [160].

TMAO has a clear dietary source, particularly fish and seafood [161], and dietary TMAO is readily absorbed by the host. Consequently, only a fraction of ingested TMAO is expected to reach the large intestine intact. However, direct measurements of TMAO in terminal ileal effluent are sparse, preventing a quantitative influx estimate.

DMSO can arise from dietary sources as well as from oxidation of microbially or host-derived dimethyl sulfide, but quantitative measurements of DMSO delivery to the human large intestine are lacking. I therefore classify its external influx as unknown.

Accordingly, fumarate, TMAO, and DMSO are important components of the gut respiratory repertoire. While their external influxes are expected to be low compared with oxygen, sulfate, or nitrate, current evidence does not support robust estimates of their external influx into the healthy large intestine. I therefore do not include them in the quantitative electron-acceptor total. These three resources are accordingly not currently included in the quantitative $J_{r,i}^{\mathrm{LI}}$ reconstruction (Section 8).

### 8 Mapping diet to the resource landscape of the large intestine

Building on the separate treatment of each major resource above, I formulated a combined mapping quantifying how the influx of all resources together varies across individuals and diets. This section describes this pipeline, from dietary intake through resource-specific mapping, elemental conversion, and accessibility to the growth-limitation comparison used in Figures 4 and 5.

**Model structure** To set this up mathematically, I estimate for each resource $r$ and individual $i$, the flux:

$$J_{r,i}^{\mathrm{LI}} = \mathcal{F}_r(D_i), \tag{18}$$

where $D_i$ describes the dietary intake of individual $i$ and $\mathcal{F}_r$ represents the resource-specific mapping from diet to large-intestinal influx established in the preceding sections. For resources that I treat as diet-independent, $\mathcal{F}_r$ reduces to a constant physiological influx.

The form and complexity of $\mathcal{F}_r$ differ substantially among resources, depending on the complexity of the physiology shaping resource influx and the data available. The construction of the functional forms $\mathcal{F}_r$ and potential dependencies on diet and food types are described for each resource separately in their designated section above. Five resources are mapped directly from an individual's own NHANES dietary data, at varying levels of detail: resistant starch and dietary amino acids/protein use food-item-resolved mappings built on FNDDS ingredient-level starch composition and food-category-specific true ileal digestibility, respectively; dietary fiber uses a direct passthrough of an individual's own total NHANES fiber intake under a uniform escape fraction; vitamin B12 (cobalamin) uses an individual's own NHANES dietary B12 intake directly; and phosphorus uses a calcium-modulated physiological regression on an individual's own NHANES phosphorus and calcium intake (Heaney & Nordin 2002). All remaining resources – host-derived secretions, electron acceptors, ions, dietary lipids, free sugars, polyphenols, and others – are instead assigned a fixed, population-wide physiological constant identical across individuals, using the literature-derived central estimates in

Table S1 (a diet-dependent, protein-intake-linked variant of host-secreted urea is tested separately as a sensitivity analysis, Section 6.2, but is not the primary reported quantity). I integrate these resource-specific mappings to reconstruct the combined resource landscape encountered by the large-intestinal microbiota.

**Diet dependence across the US population** To quantify variation in dietary inputs among adults in the analyzed NHANES sample, I use individual-level dietary data from the 2017–2018 National Health and Nutrition Examination Survey (NHANES). The day-one 24-hour dietary recall provides the dietary input $D_i$ for each individual, for $n = 4{,}741$ adult respondents (age $\geq 20$ years; ages $\geq 80$ years are top-coded as 80 in NHANES). Where total nutrient intake is sufficient for the corresponding resource mapping, I use the reported NHANES nutrient variables directly. Where resource influx depends on the identity and composition of consumed foods, I resolve reported foods into their constituent ingredients using the USDA Food and Nutrient Database for Dietary Studies (FNDDS, 2017–2018 release) and apply the food-specific mappings described above. In this way, NHANES provides the sample distribution of day-one dietary inputs, while the physiological analyses above determine how these dietary inputs are transformed into the resource influxes encountered by the large-intestinal microbiota.

The current implementation uses the day-one 24-hour dietary recall only and reports unweighted summaries and sample fractions. Accordingly, the resulting distributions describe modeled respondent-days in the analyzed NHANES sample, not survey-weighted national estimates or stable classifications of habitual individual intake. As a sensitivity check, Figure S2 recomputes the key dietary and colon-reaching distributions and the population-level binding-resource fractions using the day-one dietary sample weight (`WTDRD1`); applying this weighting shifts these distributions and fractions only modestly and does not change which resource is identified as limiting for the large majority of individuals. Population-wide physiological constants are combined with individual dietary inputs unless a resource mapping is explicitly individual-specific. Matching to the phosphorus mapping's food-composition data resulted in no sample attrition (all food records matched a food code). Missing-data treatment and FNDDS ingredient-match coverage are reported separately within each resource's own subsection above.

The carbon/nitrogen growth-limitation calculations additionally exclude 56 individuals (1.2%) with implausible day-one energy intake ($< 500$ kcal/day), giving an analytic sample of $n = 4{,}685$ for those results specifically; all other summary statistics reported use the full $n = 4{,}741$.

### 9 Conversion to growth-accessible elemental supplies

To assess whether bacterial growth may plausibly be constrained by the availability of specific elements (carbon, nitrogen, phosphorus, sulfur, magnesium, and the major inorganic ions potassium, sodium, and calcium), I first convert the flux estimates of the resources established above (Table S1) into elemental fluxes. For resource $r$, element $E$, and individual $i$, I define the growth-accessible elemental influx as

$$J^{\mathrm{acc}}_{E,r,i} = J^{\mathrm{LI}}_{r,i}\, f_{E,r}\, a_r, \tag{19}$$

where $f_{E,r}$ is the mass fraction of element $E$ in resource $r$ (Table S3) and $a_r$ is the resource-level effective utilization fraction implemented in the current analysis (Table S3). It is applied uniformly to every element in resource $r$: depending on the underlying evidence, this operational factor may reflect resource disappearance, degradation, fermentation, or assumed availability rather than direct element-specific assimilation. The total growth-accessible supply of element $E$ is then

$$J_{E,i}^{\text{acc}} = \sum_r J_{E,r,i}^{\text{acc}}. \tag{20}$$

It is this growth-accessible supply, $J_{E,i}^{\text{acc}}$, rather than total colon-reaching mass, that is carried forward into the biomass-support analysis below. The following subsections establish the substrate-specific elemental mass fractions $f_{E,r}$ used above, organized by element.

#### 9.1 Carbon content of resources and carbon demand

Carbon constitutes both the dominant energy source for bacterial growth and one of the principal anabolic building blocks for biomass synthesis. It is supplied by essentially every organic substrate considered in this work, so establishing the elemental carbon influx requires a substrate-specific carbon mass fraction for each contributing category, derived either from exact molecular stoichiometry (for well-defined small molecules) or from representative compositional data (for heterogeneous polymer classes).

**Carbon from resistant starch and free sugars** From exact molecular formulae, resistant starch (a glucan polymer, $(C_6H_{10}O_5)_n$) is $72/162 \approx 44\%$ carbon by mass and free glucose ($C_6H_{12}O_6$) is $72/180 = 40\%$, applied directly to the resistant starch (Section 2.1.1) and free sugar (Section 2.1.3) fluxes respectively.

**Carbon from dietary fiber** Dietary fiber comprises a heterogeneous mixture of polysaccharides – cellulose, hemicellulose, pectins, arabinoxylans, $\beta$-glucans, and inulin – whose monomer composition varies by polysaccharide class but is uniformly carbohydrate-derived. I adopt a representative carbon content of 40–45% by mass, consistent with the range spanned by these individual polysaccharide classes and closely matching the stoichiometric starch value above, since both are built predominantly from hexose-derived monomer units.

**Carbon from protein and amino acids** I adopt a representative carbon content of 53% by mass, consistent with standard protein elemental composition tables, applied uniformly to dietary protein escaping absorption and to endogenous (small-intestinal) protein and free amino acid losses (Section 2.2).

**Carbon from lipids** I adopt a representative carbon content of approximately 76% by mass, consistent with standard triglyceride/fatty acid composition for typical dietary $C_{16}$–$C_{18}$ chain lengths; I do not further resolve this factor by lipid class (triglyceride vs. phospholipid vs. free fatty acid), given the limited molecular characterization of lipids reaching the large intestine discussed in Section 2.3.

**Carbon from mucin** The intestinal mucus layer is built primarily from the gel-forming mucin MUC2 [162, 163]. MUC2 consists of approximately 80% O-linked glycans and 20% protein backbone by mass [163, 164]. The protein backbone contains about 53% carbon by mass, matching the general protein composition value above. Unlike dietary carbohydrates, the glycan chains contain appreciable nitrogen because they are rich in *N*-acetylgalactosamine, *N*-acetylglucosamine, and sialic acids, and together contain approximately 47% carbon by mass. The mass-weighted carbon content of whole mucin is therefore

$$f_C^{\text{mucin}} = 0.8 \times 0.47 + 0.2 \times 0.53 \approx 0.48,$$

i.e. approximately 48% carbon by mass. This factor is applied to the total mucin mass flux discussed in Section 6.4 and Section 9.2, noting that the underlying mucin mass estimate itself carries substantial uncertainty (see Section 9.2 for discussion of the discrepancy between turnover-based and glycoconjugate-based mucin mass estimates).

**Carbon from polyphenols** Polyphenols (Section 2.6) are aromatic, phenolic-ring-dominated structures with a correspondingly higher carbon content than typical carbohydrates. Based on representative flavonoid and phenolic-acid molecular formulae, I adopt a carbon content of approximately 60–65% by mass. As noted in Section 2.6, this mass-based carbon content should not be read as implying that an equivalent fraction of polyphenol carbon is available for microbial growth: the extent to which polyphenol carbon is actually catabolized or incorporated into biomass, as opposed to being ring-cleaved without assimilation or passing through largely unmodified, cannot currently be inferred from mass alone. The 30% utilizable fraction adopted for this carbon in the growth-limitation model (Table S3) is accordingly a first-pass modeling assumption rather than a directly measured quantity.

**Carbon from bile acids** Bile acids (Section 3.3) are steroid-derived molecules. Unconjugated forms (e.g. cholic acid, $C_{24}H_{40}O_5$, MW 408.6) have a carbon content of approximately 70% by mass, while taurine- or glycine-conjugated forms (e.g. taurocholic acid, $C_{26}H_{45}NO_7S$, MW 515.7) have a somewhat lower carbon content of approximately 60% by mass owing to the added conjugate moiety. I adopt a representative range of 60–70% depending on conjugation state, noting that the dominant human conjugation pattern (glycine:taurine $\approx$ 3:1, Section 3.3) favors values toward the lower end of this range. I note that this carbon-content figure describes the elemental composition of the bile-acid molecule itself and should not be conflated with microbial growth-accessibility: the deconjugation and dehydroxylation reactions discussed in Section 3.3 demonstrate that bacteria chemically transform bile acids, not that the resulting carbon skeleton is assimilated into biomass or catabolized for energy. The growth-limitation model nonetheless treats bile-acid carbon as 100% utilizable (Table S3), an upper-bound, first-pass modeling assumption adopted in the absence of direct evidence for or against bacterial assimilation of bile-acid carbon specifically; bile acids' overall influx is small relative to the major carbon sources (Table S1), so this assumption is unlikely to materially affect the population-level growth-limitation classification.

#### 9.2 Nitrogen content of resources

Nitrogen reaches the large intestine through several chemically distinct routes, including dietary proteins escaping digestion, ammonium, urea, and host-derived mucus. As for carbon, converting the underlying substrate fluxes into an elemental nitrogen flux requires a substrate-specific conversion factor for each contributing route.

**Nitrogen from protein and amino acids** I use the standard nitrogen-to-protein conversion factor of 6.25 (i.e. protein is assumed to be 16% nitrogen by mass), following the conventional Jones factor [165], applied uniformly to dietary protein reaching the large intestine and to endogenous (small-intestinal) protein and free amino acid losses. I note that Chacko & Cummings [30] report that, across three different diets, proteins in the ileum consistently accounted for 45–50% of total ileal nitrogen, while free amino acids accounted for only 5–11%, indicating that a substantial fraction of amino-acid-derived nitrogen exits the ileum in peptide or intact protein form rather than as free amino acids; the single 16%-nitrogen conversion factor applies equally to all three forms and is therefore unaffected by this distinction.

**Nitrogen from mucin** As derived in Section 9.1, mucin glycans contain approximately 4% nitrogen by mass (via *N*-acetylgalactosamine, *N*-acetylglucosamine, and sialic acid residues) and the MUC2 protein backbone contains approximately 16% nitrogen by mass, consistent with standard protein composition. Combined with the 80:20 glycan:protein mass split, the mass-weighted nitrogen content of whole mucin is

$$f_N^{\text{mucin}} = 0.8 \times 0.04 + 0.2 \times 0.16 \approx 0.064,$$

i.e. approximately 6.4% nitrogen by mass – notably higher than typical dietary carbohydrate (negligible N) but lower than pure protein, consistent with a C:N ratio of approximately 7.5:1 by mass (approximately 9:1 molar) for whole mucin. I note that two available estimates of total mucin mass reaching the large intestine differ substantially – an indirect turnover-based estimate of approximately 4.4 g/day combining ileal and local colonic contributions [5], versus a direct glycoconjugate-based measurement of approximately 15 g/day in terminal ileostomy effluent [70] – and this conversion factor should be applied to whichever mass estimate is adopted, rather than resolving that discrepancy here.

**Nitrogen from urea** Urea ($CH_4N_2O$, MW 60.06) contains two nitrogen atoms per molecule, giving a nitrogen content of $28/60.06 \approx 46.7\%$ by mass. This factor applies both to the small ileal urea efflux (Section 3.4) and to the substantially larger flux of urea hydrolyzed within the large intestine itself (Section 6.2); I treat these as two separate line items given their different physiological origin – unabsorbed dietary/ileal urea versus host urea actively secreted and subsequently hydrolyzed in situ – even though the elemental conversion factor is identical for both.

**Nitrogen from ammonium and nitrate** Ammonium and nitrate fluxes are reported in molar units (mmol/day) in the ileal effluent literature (Sections 3.4 and 7.2); since each mole of either ion contains exactly one mole of nitrogen, 1 mmol $NH_4^+$ or $NO_3^-$ corresponds directly to 14 mg N (the atomic mass of nitrogen), with no mass-fraction conversion needed. Given the small absolute magnitude of both fluxes relative to protein- and urea-derived nitrogen, their contribution to total nitrogen influx is expected to be minor.

**Other nitrogen-containing compounds** In addition to the major nitrogen sources above, several other nitrogen-containing compounds may enter the large intestine, including nucleic acids and their degradation products (nucleotides, nucleosides, purines, pyrimidines), amino sugars derived from mucus glycans and bacterial cell walls, and polyamines (e.g. putrescine and spermidine). With the exception of DNA, I am not aware of quantitative measurements of their concentration in terminal ileal contents or their daily flux into the human colon. Total nitrogen efflux measurements together with quantified amino acid, urea, and ammonium content suggest that these components collectively account for less than 5% of total ileal nitrogen efflux [31]. Point measurements of DNA-bound nitrogen (approximately 0.2 mg N per 40 mg total N per gram dry matter) confirm a very small contribution from DNA specifically [31]. I do not assign individual conversion factors to these compounds here, since their combined contribution has been bounded only as an upper limit rather than quantified as a discrete flux; this upper bound should be treated as a check on, rather than an additive component of, the total nitrogen budget.

##### 9.3 Sulfur content of resources

Several chemically distinct sulfur sources contribute to the overall influx of sulfur into the large intestine. As for carbon and nitrogen, each requires its own conversion basis.

**Sulfur from dietary protein** Proteins reaching the large intestine provide sulfur through their methionine and cysteine residues, the two sulfur-containing amino acids. As a first-order approximation, I assume that the amino-acid composition of undigested dietary protein is constant and representative of a mixed diet. Food-composition data indicate that methionine and cysteine typically account for approximately 2.2% and 1.5% of dietary protein mass, respectively, although their abundance varies among individual protein sources [166]. Using central values of 2.2% methionine and 1.5% cysteine, and accounting for the fraction of each amino acid constituted by sulfur (methionine: S is approximately 21.5% of residue mass; cysteine: S is approximately 26.5% of residue mass), gives an elemental sulfur content of approximately 0.9% of protein mass overall. This factor is applied to the same dietary protein flux used for nitrogen (Section 9.2).

**Free sulfate** Each mole of sulfate contains one mole of sulfur, so an ileal sulfate influx of 1–12 mmol/day corresponds directly to 1–12 mmol S/day, or approximately 30–400 mg S/day.

**Sulfur from taurine-conjugated bile acids** Bile acids provide an additional source of sulfur because a fraction is conjugated to taurine before secretion from the liver. For a fully taurine-conjugated bile acid pool, molecular masses of approximately 500–516 g/mol for the major taurine-conjugated bile acids and an atomic sulfur mass of 32 g/mol correspond to an elemental sulfur content of approximately 6.2–6.4% by mass. Because glycine conjugation predominates in humans, with a glycine:taurine conjugation ratio of approximately 3:1 [167], I use a taurine-conjugated fraction of approximately 20–25% to obtain a central estimate of the sulfur associated with the bile-acid pool reaching the terminal ileum. This calculation assumes that taurine- and glycine-conjugated bile acids have comparable fractional recovery at the ileum and should therefore be regarded as an approximation. Importantly, deconjugation does not itself remove sulfur from the intestinal sulfur pool: taurine released by microbial bile-salt hydrolases retains its sulfur, and its contribution to colonic sulfur influx depends on the subsequent absorption and metabolism of free taurine. I therefore retain the fully taurine-conjugated calculation as an upper bound and the 20–25% calculation as a representative central estimate.

**Sulfur from mucus** Mucins supply sulfur through both sulfur-containing amino acids within the MUC2 protein backbone and sulfate groups esterified to mucin O-glycans; I treat these as two separate components. For the protein backbone, cysteine and methionine constitute approximately 5–6% and 1% of MUC2 protein mass, respectively [168]. Using the same per-residue sulfur contents as for dietary protein above gives an estimated elemental sulfur content of approximately 1.5–1.8% by mass of the MUC2 protein backbone. Assuming that protein constitutes approximately 20% of total mucin mass, this corresponds to approximately 0.3–0.36% sulfur by total mucin mass.

For the glycan component, sulfur is present as sulfate ester groups attached to O-linked glycans. Biochemical measurements of purified human intestinal mucin report approximately 0.45–1.6 g $SO_4$ per 100 g mucin, including values around 1.2 g $SO_4$ per 100 g mucin [169, 170]. Because sulfur accounts for one-third of sulfate mass, these measurements correspond to approximately 0.15–0.53% elemental sulfur by total mucin mass, with approximately 0.40% corresponding to the central 1.2 g $SO_4$/100 g mucin estimate. Because these measurements are already reported relative to total mucin mass, no additional multiplication by the assumed glycan mass fraction is applied.

Combining the protein-backbone (0.3–0.36%) and glycan (0.15–0.53%) contributions as a single blended range for total mucin sulfur – rather than simply summing each component's independent low and high extremes, which would assume both simultaneously take their most extreme value – gives the 0.44–0.76% range reported in Table S3, kept as one blended value since sulfur, unlike carbon and nitrogen, is not split by macro-component in the resource-limitation model.

**Additional sulfur sources** Sulfur in the protein portion of sloughed epithelial cells is represented in Table S3 using an elemental sulfur fraction of 0.45%. Additional sulfur sources not assigned explicit conversion factors include glutathione and other thiols and sulfated host metabolites more broadly. Their combined contribution is expected to be small relative to dietary protein and free sulfate, but is not currently quantified.

##### 9.4 Direct elemental measurements of magnesium, phosphorus, and major ions

The aggregate ileal or fecal influxes of magnesium, phosphorus, potassium, sodium, and calcium are already reported as elemental quantities and therefore use a conversion factor of 1.0. Phosphorus contributed through sloughed epithelial cells is an exception: this substrate-specific contribution is converted using the epithelial-cell phosphorus fraction reported in Table S3 rather than a unity factor.

I emphasize that most conversion factors in this section (Sections 9.1–9.3) are based on stoichiometric or representative compositional data rather than on direct measurement of elemental content within ileal or fecal material – partial exceptions being mucin composition and bile acid conjugation state, where more direct biochemical measurements exist. The magnesium/phosphorus/ion group here is the cleanest exception of all: since these influxes are already reported as elemental quantities, no compositional estimation is involved at all.

##### 9.5 Microbial accessibility of incoming resources

Total colon-reaching influx and growth-accessible influx are not generally identical. I use $a_r$ as a community-level effective fraction: the fraction of incoming resource $r$ that becomes available for microbial catabolism or assimilation during large-intestinal transit, applied uniformly to every element contained in that resource

(Section 9). This is not the fraction available to every species, and it does not eliminate spatial, kinetic, or taxon-specific restrictions.

I distinguish three reasons why $a_r$ may be below unity. First, some material is chemically excluded from appreciable microbial degradation under large-intestinal conditions – lignin (not a carbohydrate) and dietary lipids (not a fermentation substrate for gut bacteria) are the clearest examples, both assigned $a_r = 0$ (Table S3). Second, substrates such as cellulose and hemicellulose may be only partly degraded during transit (assigned 30% and 60%, respectively), whereas free sugars and several soluble fiber classes such as pectin and inulin-type fructans are used far more completely (90–100%). Third, degradation or fermentation does not necessarily imply quantitative incorporation of every constituent element into biomass: bile acids, for instance, are assigned a 100% utilizable fraction because they are near-universally deconjugated/dehydroxylated (biotransformed) by gut bacteria, even though this transformation does not itself constitute assimilation of bile-acid carbon into microbial biomass. Together, these considerations span the full observed range of accessibility values, from 0% (lignin, lipids) to 100% (e.g. resistant starch, free sugars, most amino acids, ammonium). The accessibility factor therefore represents an operational supply term for this analysis and must be interpreted according to the evidence available for each resource.

I assign central accessibility fractions and, where possible, uncertainty ranges using compositional exclusion, comparative fermentation measurements, or intestinal mass-balance evidence. These assignments and their evidential bases are reported in Table S3.

For phosphorus, quantitative microbial accessibility remains particularly poorly constrained. I therefore use $a_P = 1$ only as an explicit upper-bound supply scenario in the central calculation, not as a measured best estimate. Consequently, any conclusion about phosphorus sufficiency or limitation is conditional on this assumption and is revisited in a one-parameter (yield-only) sensitivity analysis below (Figure 4F) and, jointly with accessibility, in a two-dimensional sensitivity analysis (Figure S4E).

Multiplying resource influx, elemental content, and accessibility gives the growth-accessible elemental influx in Eq. 19. Only this accessible quantity is carried forward into biomass-support calculations.

### 10 Growth limitations

I next estimate how much bacterial dry-biomass production could in principle be supported by the reconstructed growth-accessible supplies. The calculation does not represent realized production and does not capture spatial gradients, uptake kinetics, taxon-specific access, or competition among organisms.

#### 10.1 Biomass elemental requirements and yields

To start with I consider the biomass which can be supported per influx of resources. This step is commonly done on the elementary level, considering the biomass supported given the influx of a certain growth supporting element. For the major biomass-forming elements C, N, P, and S, the amount of biomass that can be supported by a given elemental supply can be estimated from elemental biomass yields. I define the elemental biomass yield as

$$Y_{X/E} = \frac{\text{g dry biomass}}{\text{g element } E}.$$

For elements that are primarily incorporated into cellular biomass, this yield is approximately the inverse of the mass fraction of the respective element in bacterial dry biomass. Carbon requires additional consideration because organic carbon substrates are used not only for biomass synthesis but also to provide energy for growth. Bacterial elemental composition and biomass yields are also not fixed but can vary among species and with environmental conditions, nutrient limitation, and growth rate. I therefore use representative central values together with ranges reflecting experimentally observed variation. Table S4 summarizes the central yield, physiological range, and confidence level I adopt for each element; the paragraphs below explain the evidence and reasoning behind each entry in turn.

**Carbon** Carbon differs fundamentally from the other elemental requirements considered here because organic carbon substrates serve both as building blocks for cellular biomass and as sources of energy. Consequently, it is useful to distinguish between an *anabolic carbon requirement*, representing the carbon incorporated into newly synthesized biomass, and the larger *total carbon requirement*, which additionally accounts for substrate carbon consumed to provide the energy required for biomass synthesis and cellular maintenance.

The anabolic carbon requirement follows directly from bacterial elemental composition. Carbon typically constitutes approximately $45–55\%$ of bacterial dry biomass [171, 172]. Thus, if carbon were required only as a building block, one gram of carbon could support approximately two grams of bacterial dry biomass. I therefore use

$$Y_{X/C,\text{anabolic}} = 2.0\ \text{g DW g}^{-1}\ \text{C}$$

with a representative physiological range of

$$Y_{X/C,\text{anabolic}} = 1.8\text{–}2.2\ \text{g DW g}^{-1}\ \text{C}.$$

The additional catabolic carbon requirement can be understood from bacterial bioenergetics. Synthesis and assembly of one gram of bacterial dry biomass requires substantial metabolic energy. For *E. coli*, theoretical accounting of biosynthetic reactions gives an ATP requirement of approximately $35\ \text{mmol ATP g}^{-1}\ \text{DW}$ [173].

How much carbon must be consumed to provide this energy depends strongly on the available catabolic pathways and, in particular, on the availability of external electron acceptors [139]. In the healthy large intestine, the influx of carbohydrate carbon is large compared with the influxes of strong external electron acceptors such as oxygen, nitrate, and sulfate. Specifically, a typical carbohydrate influx of approximately 19 g/day corresponds to roughly 106 mmol glucose equivalents and, if completely oxidized, would release approximately 2.5 mol electron equivalents per day. By comparison, even an upper-bound combined influx of approximately 15 mmol/day of oxygen, nitrate, and sulfate provides at most approximately 0.12 mol electron-accepting equivalents per day, assuming the maximal eight-electron reduction of sulfate. Thus, even under this deliberately permissive upper-bound calculation, these external electron acceptors could support complete respiratory oxidation of less than approximately 5% of the incoming carbohydrate. Thus, external oxygen, nitrate, and sulfate can account for only a small fraction of the electron-accepting capacity required for complete carbohydrate oxidation, supporting fermentation as the dominant mode of bulk carbohydrate catabolism in the healthy large intestine.

Under fermentative conditions, major pathways used by gut bacteria typically generate on the order of $2$–$4$ ATP per glucose equivalent, although the exact yield depends on the pathway and fermentation products formed [5]. Combining these values predicts a carbohydrate requirement of approximately

$$\frac{35\ \mathrm{mmol\,ATP\,g^{-1}\,DW}}{2\text{–}4\ \mathrm{mol\,ATP\,mol^{-1}\,glucose}} \simeq 9\text{–}18\ \mathrm{mmol\,glucose\ equivalents\,g^{-1}\,DW}.$$

Thus, energetic considerations alone predict that substantially more organic carbon must be consumed during anaerobic growth than is ultimately retained in newly formed biomass.

This additional carbon requirement is intrinsically linked to redox balance. In the absence of sufficient external terminal electron acceptors, organic metabolic intermediates commonly serve as electron acceptors, regenerating oxidized electron carriers and subsequently being released as fermentation products. These products retain substantial amounts of the carbon originally present in the consumed substrate. Different gut bacteria employ different combinations of fermentative pathways and consequently release different products, but all face the common constraints of retaining sufficient carbon for biomass synthesis while generating sufficient ATP and maintaining redox balance [5, 139].

These bioenergetic considerations provide an independent expectation for carbon demand. For my quantitative analysis, however, I use direct measurements of carbohydrate consumption across a phylogenetically diverse panel of major human gut bacteria [5]. Arnoldini et al. quantified carbohydrate uptake, biomass production, and fermentation-product release, allowing carbohydrate consumption to be expressed as glucose equivalents per gram of newly formed bacterial dry biomass. For the present analysis, I use the strain-averaged measurements obtained in YCA medium. After excluding *Eubacterium siraeum*, whose exceptionally high apparent uptake was associated with its maltose measurement, measurements were available for 21 gut bacterial strains. Across these strains, total carbohydrate uptake averaged

$$12.17\ \mathrm{mmol\,glucose\ equivalents\,g^{-1}\,DW},$$

with individual strain averages ranging from

$$6.00\text{–}24.76\ \mathrm{mmol\,glucose\ equivalents\,g^{-1}\,DW}.$$

Because one mmol of glucose contains $6 \times 12 = 72$ mg carbon, the mean uptake corresponds to

$$12.17 \times 0.072 = 0.876\ \mathrm{g\,carbohydrate\ C\,g^{-1}\,DW}.$$

If all carbon incorporated into new biomass were derived from these carbohydrates, this would correspond directly to a carbon yield of

$$Y_{X/C} = 1/0.876 = 1.14\ \mathrm{g\,DW\,g^{-1}\,C}.$$

However, the growth medium also contains amino acids and other organic compounds that can contribute directly to anabolic biomass carbon. Consequently, the measured carbohydrate uptake does not account for all carbon consumed during growth, and $1.14\ \mathrm{g\,DW\,g^{-1}\,C}$ should be interpreted as a carbohydrate-carbon yield rather than a complete elemental carbon yield. The total carbon requirement can be written as

$$C_{\text{total}} = C_{\text{CH}} + f_{\text{other}} C_{\text{biomass}},$$

where $C_{\text{CH}} = 0.876\,\text{g C g}^{-1}\,\text{DW}$ is the measured carbohydrate-carbon consumption, $C_{\text{biomass}} \simeq 0.50\,\text{g C g}^{-1}\,\text{DW}$ is the carbon content of newly formed biomass, and $f_{\text{other}}$ denotes the fraction of biomass carbon supplied directly by non-carbohydrate substrates.

For the central working scenario, I assume that half of biomass carbon is supplied by non-carbohydrate substrates,

$$f_{\text{other}} = 0.5.$$

The limiting scenarios $f_{\text{other}} = 0$ and $f_{\text{other}} = 1$ are retained as sensitivity bounds below.

This provides a simple intermediate assumption between the two limiting cases and is physiologically plausible given the substantial contribution of proteins to bacterial biomass and the availability of amino acids in the growth medium. The resulting total carbon requirement is

$$C_{\text{total}} = 0.876 + 0.5 \times 0.50 = 1.126\ \text{g C g}^{-1}\,\text{DW},$$

corresponding to

$$Y_{X/C,\text{total}} = 0.89\,\text{g DW g}^{-1}\,\text{C}.$$

To quantify the uncertainty associated specifically with the origin of anabolic carbon, I consider the two limiting cases. If all biomass carbon originates from carbohydrates ($f_{\text{other}} = 0$), the measured carbohydrate uptake already contains both anabolic and catabolic carbon demand, giving

$$Y_{X/C,\text{total}} = 1.14\ \text{g DW g}^{-1}\,\text{C}.$$

At the opposite extreme, if none of the biomass carbon originates from carbohydrates ($f_{\text{other}} = 1$), an additional $0.50\,\text{g C}$ from other substrates must be consumed per gram of biomass, giving

$$Y_{X/C,\text{total}} = \frac{1}{0.876 + 0.50} = 0.73\,\text{g DW g}^{-1}\,\text{C}.$$

I therefore use

$$Y_{X/C,\text{total}} = 0.89\ \text{g DW g}^{-1}\,\text{C}$$

as my central estimate, with

$$Y_{X/C,\text{total}} = 0.73\text{–}1.14\,\text{g DW g}^{-1}\,\text{C}$$

as the range associated with uncertainty in the anabolic origin of cellular carbon.

This accounting range should be distinguished from the observed variation in carbohydrate consumption among bacterial strains. Across the 21 gut bacterial strains considered here, carbohydrate uptake varied from $6.00$ to $24.76\ \text{mmol}$ glucose equivalents per gram dry biomass. This interspecies variation represents genuine physiological variation in carbohydrate utilization and can be considered separately when assessing sensitivity to bacterial community composition. The $0.73\text{–}1.14\,\text{g DW g}^{-1}\,\text{C}$ range instead isolates the uncertainty involved in converting the mean measured carbohydrate requirement into a total elemental carbon requirement.

The aggregate effective yield derived above is used for the population-typical comparison in Figure 3. It is not an input to the individual-level model used for Figures 4–5. That model instead keeps carbohydrate carbon and amino-acid carbon as separate supply pools and allocates each pool between anabolic and catabolic uses. The two calculations therefore address the same carbon-demand problem at different levels of resolution and should not be interpreted as one single parameterization. In particular, the per-individual model at no point invokes $f_{\text{other}}$, so there is no risk of double-counting non-carbohydrate carbon between this population-typical yield and the explicit protein/amino-acid, mucin, lipid, polyphenol, and bile-acid carbon contributions already tracked in the elemental-influx accounting (Section 9).

The experimentally measured carbohydrate requirement is also consistent with independent bioenergetic expectations: the mean uptake of $12.17\ \mathrm{mmol}$ glucose equivalents per gram biomass lies within the approximately $9$–$18\ \mathrm{mmol\,g^{-1}}$ range predicted from the ATP requirement for biomass synthesis and typical fermentative ATP yields. At very slow growth rates, maintenance energy can constitute a larger fraction of cellular energy expenditure and further increase carbon consumption per unit biomass; the values used here instead reflect direct measurements during active growth of representative gut bacteria.

Carbohydrate consumption was measured directly and systematically across major human gut bacterial strains, and its magnitude agrees with independent bioenergetic predictions. Conversion to total elemental carbon demand requires an assumption regarding the fraction of biomass carbon supplied by amino acids and other non-carbohydrate substrates; I explicitly bracket this uncertainty using the two limiting cases.

**Nitrogen** Nitrogen constitutes a comparatively large and relatively stable fraction of bacterial biomass, predominantly because of its incorporation into proteins and nucleic acids. Compared with phosphorus and several inorganic ions, experimentally measured cellular nitrogen fractions vary surprisingly little. The compilation of Rajpurohit and Eiteman reports biomass yields of approximately $6$–$8\ \mathrm{g\,DW\,g^{-1}\,N}$ for *Bacillus* species and approximately $7$–$10\ \mathrm{g\,DW\,g^{-1}\,N}$ for *E. coli* under a range of nutrient limitations [174]. In *E. coli*, nitrogen-limited cultures contain approximately $10$–$11\%$ N by dry mass, whereas cultures limited by carbon or iron contain approximately $12.5$–$13.5\%$ N, demonstrating some physiological adjustment but considerably less variation than observed for phosphorus or iron [174]. Measurements across cultured and environmental bacteria similarly place nitrogen near $\sim 10$–$15\%$ of cellular dry mass and indicate comparatively limited variation in cellular N:C ratios [171].

I therefore use

$$Y_{X/N} = 7.5\ \mathrm{g\,DW\,g^{-1}\,N},$$

corresponding to $13.3\%$ N by dry mass, with a representative range of approximately

$$Y_{X/N} = 6\text{–}10\ \mathrm{g\,DW\,g^{-1}\,N}.$$

This value (Table S4) is used for the population-typical, single-scenario comparison in Figure 3, analogous to $Y_{X/C,\text{total}}$ above. The per-individual coupled carbon/nitrogen optimization used for Figures 4–5 instead draws its nitrogen stoichiometry from an independent source, the biomass formula of Roels (1980) ($CH_{1.8}O_{0.5}N_{0.2}$, $q_N = 8.12$ mmol N g$^{-1}$ biomass, equivalent to $Y_{X/N} \approx 8.8$ g DW g$^{-1}$ N). As with carbon, these are two independently sourced estimates of the same underlying quantity, addressing different parts of the analysis, and should not be read as a single reconciled parameterization.

Among the elemental yields considered here, the nitrogen requirement is consequently one of the better constrained.

**Phosphorus** Phosphorus is required for nucleic acids, phospholipids, phosphorylated metabolites, and numerous cofactors, and can additionally be stored intracellular as polyphosphate. A particularly large fraction of cellular phosphorus is associated with RNA, making the phosphorus content of cells sensitive to ribosome abundance and therefore to growth rate. Measurements of bacterial phosphorus content and phosphorus-limited biomass yields consequently span a considerable range.

A compilation of experimentally determined elemental biomass yields reports phosphorus yields of approximately $28–50\ \mathrm{g\,DW\,g^{-1}\,P}$ for exponentially growing or nutrient-replete *Escherichia coli* and *Pseudomonas* strains [174]. Direct measurements by Bauer and Ziv similarly yielded values of approximately $35$, $50$, and $42\ \mathrm{g\,DW\,g^{-1}\,P}$ for *E. coli* W, B, and K-12, respectively [175]. Continuous-culture experiments with *Bacillus subtilis* demonstrate that the phosphorus requirement can decrease substantially when phosphorus itself becomes limiting. Dauner et al. measured biomass compositions corresponding to yields of approximately $38–42\ \mathrm{g\,DW\,g^{-1}\,P}$ under nitrogen limitation but approximately $65–96\ \mathrm{g\,DW\,g^{-1}\,P}$ under phosphorus limitation [176, 174].

Systematic measurements further demonstrate that cellular phosphorus content depends on physiological state. In *E. coli*, Makino et al. found substantial variation in cellular C:P stoichiometry with both phosphorus availability and growth rate [177]. Much of this growth-rate dependence can be understood from variation in cellular RNA abundance: rapidly growing cells allocate a substantially larger fraction of their biomass to ribosomes and ribosomal RNA, thereby increasing their phosphorus requirement. Variation in cellular phosphorus and its association with RNA accumulation was already observed in early studies of *E. coli* growth [178]. Across bacterial species, variation can be still larger. Measurements of 24 heterotrophic bacterial isolates revealed substantial strain-to-strain differences in cellular phosphorus quotas and in the extent to which these quotas could be reduced under phosphorus scarcity [179]. Measurements across cultured and environmental bacteria similarly demonstrate substantial variation in cellular P content [171].

Based on the clustering of measurements from *E. coli*, *Bacillus*, and *Pseudomonas* under non-extreme growth conditions, I use

$$Y_{X/P} = 40\ \mathrm{g\,DW\,g^{-1}\,P}$$

as the representative phosphorus yield. This corresponds to a phosphorus content of $2.5\%$ of bacterial dry biomass. To account for variation in species identity and physiological state, I consider a broader range of

$$Y_{X/P} = 20–100\ \mathrm{g\,DW\,g^{-1}\,P},$$

corresponding to approximately $1–5\%$ phosphorus by dry mass. The upper end of this yield range is particularly relevant under phosphorus limitation, where cells can decrease their phosphorus quota, whereas rapidly growing, ribosome-rich cells generally have greater phosphorus requirements.

**Sulfur** Sulfur is incorporated primarily into the sulfur-containing amino acids cysteine and methionine, as well as into iron–sulfur clusters and several cofactors. Direct measurements of sulfur-limited bacterial growth are less numerous than measurements for nitrogen or phosphorus. Rajpurohit and Eiteman compile sulfur biomass yields of 163 and 278 $\mathrm{g\,DW\,g^{-1}\,S}$ for *E. coli* and approximately 244 $\mathrm{g\,DW\,g^{-1}\,S}$ for *Pseudomonas putida* [174]. These values correspond to approximately 0.36–0.61% S by dry mass. Independent measurements of elemental composition across cultured and environmental bacteria also identify sulfur as a consistently present cellular element, although its abundance varies among taxa and growth conditions [180, 171].

I therefore use a representative value of

$$Y_{X/S} = 200\,\mathrm{g\,DW\,g^{-1}\,S}$$

(0.5% S by dry mass), with a working range of approximately

$$Y_{X/S} = 100\text{–}300\,\mathrm{g\,DW\,g^{-1}\,S}.$$

The lower end encompasses commonly reported bacterial compositions approaching $\sim 1\%$ S by dry mass, whereas the direct sulfur-limitation measurements generally imply lower cellular sulfur fractions.

##### 10.1.1 Inorganic ion requirements

The interpretation of Mg, Ca, K, and Na differs from that of C, N, P, and S. Whereas the latter elements are predominantly incorporated into cellular macromolecules and therefore impose relatively direct stoichiometric constraints on biomass production, Mg, Ca, K, and Na occur largely as intracellular ions. Their cellular abundances reflect both specific biochemical requirements and physiologically regulated ionic pools and can therefore vary substantially with growth rate, osmolarity, pH, external ion concentrations, and physiological state.

I nevertheless compare their influxes with representative cellular quotas to assess whether ion availability could plausibly constrain bacterial growth. The resulting biomass-support estimates should not, however, be interpreted as strict stoichiometric upper bounds analogous to those obtained for C, N, P, and S. Instead, they indicate the amount of biomass that could be supported while maintaining representative cellular ion inventories. This distinction is particularly important for K and Na, whose intracellular pools are highly adjustable and whose bulk ionic functions can in part be compensated by other ions or osmolytes.

**Magnesium** Magnesium is required as a counterion for nucleic acids and ribosomes, contributes to membrane stability, and acts as a cofactor for numerous enzymes. Unlike nitrogen or sulfur, a substantial fraction of cellular Mg is ionically associated with macromolecules rather than covalently incorporated into biomass. Accordingly, cellular Mg content changes with physiological state and particularly with ribosome abundance. Chemostat measurements summarized by Rajpurohit and Eiteman give $Y_{X/Mg}$ values ranging from approximately 400–830 $\mathrm{g\,DW\,g^{-1}\,Mg}$ for *B. subtilis* and *Klebsiella aerogenes*, with Mg requirements increasing at higher growth rates [174]. Other measurements are lower, including approximately 278 $\mathrm{g\,DW\,g^{-1}\,Mg}$ for an *E. coli* strain and 128–236 $\mathrm{g\,DW\,g^{-1}\,Mg}$ for *Pseudomonas* strains [174]. Direct elemental measurements of *E. coli* have reported Mg fractions extending to approximately 0.5–0.9% of dry mass, corresponding to yields of approximately 110–200 $\mathrm{g\,DW\,g^{-1}\,Mg}$ [172]. Measurements across bacterial species likewise demonstrate appreciable variation in cellular Mg content [180].

Thus, reported Mg requirements span several-fold, in part because cellular Mg covaries with RNA content and growth rate and because analytical approaches differ in their treatment of loosely associated cellular Mg. For the present calculations I use

$$Y_{X/Mg} = 200\,\mathrm{g\,DW\,g^{-1}\,Mg}$$

as a conservative representative value, while considering approximately

$$Y_{X/Mg} = 150\text{–}800\,\mathrm{g\,DW\,g^{-1}\,Mg}$$

as the broader experimentally supported range. Because estimated Mg supply substantially exceeds bacterial requirements in my analysis, conclusions regarding Mg limitation are insensitive to the precise value chosen.

**Potassium** Potassium is the predominant intracellular monovalent cation in many bacteria and contributes to osmotic homeostasis, maintenance of membrane potential, enzyme activity, protein synthesis, and regulation of intracellular pH. Unlike the major biomass-forming elements, however, much of cellular K belongs to a regulated intracellular ion pool rather than being stoichiometrically incorporated into macromolecules. Cellular K content can therefore vary substantially with growth conditions and extracellular osmolarity.

Direct elemental measurements of bacterial cells place potassium among the most abundant intracellular inorganic ions, while also revealing substantial variation among species and physiological states [172, 180]. Based on these measurements, I use

$$Y_{X/K} = 40\,\mathrm{g\,DW\,g^{-1}\,K}$$

as a representative apparent biomass yield, corresponding to approximately $2.5\%$ K by dry mass, and consider a physiological range of

$$Y_{X/K} = 20\text{–}60\,\mathrm{g\,DW\,g^{-1}\,K}.$$

These values should be interpreted as representative cellular inventories rather than fixed anabolic requirements. In particular, bacteria regulate intracellular K accumulation in response to osmotic and ionic conditions, and some bulk ionic functions can be partly compensated by other ions and compatible solutes. The resulting biomass-support estimate therefore provides a test of whether K influx is plausibly sufficient to maintain typical cellular K pools, rather than a strict stoichiometric limit on biomass production.

**Calcium** Calcium is present in bacterial cells and contributes to processes including envelope stability, signaling, and enzyme activity, but unlike Mg it is generally not required as a large stoichiometric component of vegetative biomass. Quantitative measurements of cellular Ca are comparatively sparse and variable. Direct X-ray microanalysis of exponentially growing *E. coli* reported approximately $0.22\%$ Ca by dry mass, corresponding to $Y_{X/Ca} \sim 450\,\mathrm{g\,DW\,g^{-1}\,Ca}$, whereas Ca was frequently below the detection limit in stationary-phase cells [172]. Measurements across bacterial species further indicate substantial variation in Ca abundance and generally place Ca among the less abundant and less consistently detected cellular elements [180].

Because the cellular Ca quota is poorly constrained and can vary by several-fold, I use

$$Y_{X/Ca} = 1{,}000\,\mathrm{g\,DW\,g^{-1}\,Ca}$$

as an approximate representative value, with a broad range of

$$Y_{X/Ca} = 400\text{–}2{,}000\,\mathrm{g\,DW\,g^{-1}\,Ca}.$$

Given the large estimated intestinal Ca supply relative to these requirements, uncertainty in this value does not materially affect my conclusions.

**Sodium** Sodium differs from K and Mg in that many bacteria maintain intracellular Na well below its extracellular concentration, and no large, universal stoichiometric Na requirement exists. Sodium nevertheless participates in membrane energetics, transport, pH regulation, and osmotic homeostasis in specific taxa and physiological conditions. Its cellular abundance therefore depends strongly on extracellular salinity, the balance of Na and K transport, and the use of sodium-motive versus proton-motive bioenergetic systems.

For this reason, an apparent biomass yield derived from a single cellular Na fraction is considerably less meaningful than the corresponding quantities for C, N, P, or S. Sodium is nevertheless included in the present ion-sufficiency comparison because it is plotted in Figure 3. The current implementation uses

$$Y_{X/Na} = 42.638\,\mathrm{g\,DW\,g^{-1}\,Na}.$$

I treat this as a provisional representative cellular inventory rather than a fixed stoichiometric biomass requirement. This value is now included in Table S4, traced to the same X-ray microanalytic elemental-composition measurements used for Mg, K, and Ca above [172]; unlike those elements, no independent literature range could be identified for cellular Na content, so this remains a single point estimate rather than a range-bounded value.

In the large intestine, sodium influx is substantially larger than the cellular amounts plausibly required for microbial biomass, so this uncertainty does not affect the conclusion that sodium supply is unlikely to directly constrain community biomass production.

### 11 Community biomass-support model

#### 11.1 Purpose and interpretation

The model estimates the maximum daily bacterial dry-biomass production that could be supported by the reconstructed resource supplies under the stated accessibility and biomass-demand assumptions. A resource is described as “binding” when its constraint determines this modeled ceiling. This does not by itself establish in vivo ecological limitation.

#### 11.2 Decision variables and parameters

Let $M$ denote supported bacterial dry-biomass production. The implementation also allocates carbon and nitrogen among catabolic and anabolic uses. The symbols appearing in the existing analysis include $C_{CH}$, $C_{AA}^{ana}$, $N_{other}$, and $N_{other}^{assim}$.

The six nonnegative decision variables are $M$, $C_{CH}^{ana}$, $C_{CH}^{cat}$, $C_{AA}^{ana}$, $C_{AA}^{cat}$, and $N_{other}^{assim}$. Here $M$ is supported bacterial dry-biomass production (g/day); the four carbon variables are carbohydrate- or amino-acid-carbon allocated to anabolic or catabolic use (mmol C/day); and $N_{other}^{assim}$ is assimilated non-amino-acid nitrogen (mmol N/day). The individual-specific inputs are total available $C_{CH}$, $C_{AA}$, and $N_{other}$. Population-wide parameters are, unless noted otherwise, derived from the biomass formula of Roels (1980), $CH_{1.8}O_{0.5}N_{0.2}$: $q_C = 40.62$ and $q_N = 8.12$ mmol g$^{-1}$ biomass (giving the formula's C:N ratio of 5); $m_{ATP} = 34.8$ mmol ATP g$^{-1}$ biomass [173, 5]; $y_{ATP,CH} = 0.534$ and $y_{ATP,AA} = 0.103$ mol ATP mol$^{-1}$ C, the per-carbon ATP yields from carbohydrate and amino-acid catabolism [5]; $\eta_C = 1.0$, i.e. no loss assumed when amino-acid carbon is used anabolically; $\phi_{AA} = 1.0$, i.e. no cap on catabolic amino-acid use beyond the LP's own energy balance; and $r_{NC,AA} = 0.259$ mol N mol$^{-1}$ C, the amino-acid pool's nitrogen:carbon ratio – obtained from the protein nitrogen and carbon mass fractions already introduced above (Sections 9.1, 9.2), $0.16/0.53 = 0.302$ g N g$^{-1}$ C, converted from a mass to a molar ratio via the carbon:nitrogen atomic-mass ratio ($\times 12.011/14.007$). I denote the optimum by $M_{max}$. Phosphorus and sulfur are not constraints inside this LP; their elemental ceilings are calculated separately and compared post hoc with $M_{max}$.

#### 11.3 Optimization problem

The optimization objective is

$$\max M. \tag{21}$$

Subject to nonnegativity, the implemented constraints are

$$
\begin{aligned}
q_C M &\le C_{CH}^{ana} + \eta_C C_{AA}^{ana}, && \text{anabolic carbon balance,} && (22)\\
q_N M &\le r_{NC,AA} C_{AA}^{ana} + N_{other}^{assim}, && \text{biomass nitrogen balance,} && (23)\\
m_{ATP} M &\le y_{ATP,CH} C_{CH}^{cat} + y_{ATP,AA} C_{AA}^{cat}, && \text{ATP balance,} && (24)\\
C_{CH}^{ana} + C_{CH}^{cat} &\le C_{CH}, && \text{carbohydrate-carbon budget,} && (25)\\
C_{AA}^{ana} + C_{AA}^{cat} &\le C_{AA}, && \text{amino-acid-carbon budget,} && (26)\\
C_{AA}^{cat} &\le \phi_{AA} C_{AA}^{cap}, && \text{catabolic amino-acid cap,} && (27)\\
N_{other}^{assim} &\le N_{other}, && \text{non-amino-acid nitrogen budget.} && (28)
\end{aligned}
$$

In the baseline solve, $C_{AA}^{cap} = C_{AA}$ and $\phi_{AA} = 1$, so the catabolic amino-acid cap adds no restriction beyond the total amino-acid-carbon budget. The separate cap is retained for counterfactual relaxed-resource solves, where it prevents an unlimited amino-acid supply from also creating unlimited catabolic capacity.

The optimization is lexicographic. I first maximize $M$. Holding $M = M_{max}$, I minimize amino-acid carbon catabolism; holding both preceding optima fixed, I minimize total resource use. These tie-breaks do not change $M_{max}$ but select a reproducible allocation among multiple biomass-maximizing solutions. They therefore matter for allocation-derived quantities such as $N_{other}^{assim}$.

##### 11.4 Individual-level implementation

For each NHANES individual $i$, the model receives that individual's mapped diet-dependent supplies together with the population-wide physiological inputs defined above and returns $M_{max,i}$ and the associated resource allocations.

The LP is solved with `scipy.optimize.linprog` (SciPy 1.15.2) using the `highs` method and its default feasibility tolerances. The all-zero solution is feasible for nonnegative inputs, but the implementation does not currently assert successful termination or explicitly report solver failures. The current C–N analysis contains 4,685 adult respondents; joining this cohort to the separately computed phosphorus mapping (Section 12) resulted in 0 of 4,685 individuals dropped for lacking a matching phosphorus estimate.

#### 12 Modeled biomass-supporting constraints

The aggregate calculation used for Figure 3 applies the elemental yields in Table S4 to population-typical accessible supplies. The individual-level Figures 4–5 instead use the substrate-resolved C–N LP above; P is compared post hoc, while S and the inorganic ions remain population-typical sufficiency checks.

Within this framework, the elemental ceiling associated with element $E$ is

$$M_{\text{supported},E} = J_E^{\text{acc}} Y_{X/E} = \frac{J_E^{\text{acc}}}{q_E}, \tag{29}$$

where $q_E$ is the amount of element $E$ required per gram bacterial dry biomass. For carbon and coupled carbon–nitrogen allocation, the full model specified above is used rather than this simple quotient.

As a concrete illustration of Equation 29, for carbon the population-typical growth-accessible influx is $J_C^{\text{acc}} = 15.77$ g C/day (Table S1) and the biomass yield is $Y_{X/C} = 0.89$ g dw/g C (Table S4), giving a supported biomass of $M_{\text{supported},C} = 15.77 \times 0.89 \approx 14.0$ g dry biomass/day, matching the value reported in the main text and shown in Figure 3D.

**Nitrogen versus carbon limitation across individuals** I perform these calculations for typical resource influxes (Table S1), presented in Figure 3. Furthermore, I analyze the variation of major influx values with variation in diet, using NHANES dietary data (Section 8).

To account for food-specific variation in amino-acid digestibility and carbohydrate content reaching the large intestine, I use the food-specific digestibility values described in the corresponding sections above. Colon-reaching protein is also converted to amino-acid nitrogen assuming all amino acids including those entering as intact proteins are fully utilized by the microbes in the gut (the same lower-bound assumption discussed in the main text's analysis of amino acid auxotrophies). Free (non-amino-acid) nitrogen, $N_{other}$, combines a population-wide constant baseline (ileal urea carryover, ileal ammonium, and endogenous glycan-derived nitrogen) with a diet-dependent urea contribution: urea is actively secreted and hydrolyzed across the large-intestinal epithelium, modeled as a linear function of the individual's own dietary protein intake (interpolated between two measured anchor points, Meakins & Jackson 1996, and floored at the low-protein measured value rather than extrapolated toward zero). Solving the resource-limitation LP for each individual's own $(C_{CH}, C_{AA}^{ana}, N_{other})$ gives that individual's LP-derived joint C–N biomass-support ceiling, $M_{max}$ (the same quantity underlying Figure 3D, now computed per individual rather than at the population-typical influx).

To isolate nitrogen's own contribution, I additionally compute a nitrogen-only ceiling, $M_N = N_{tot}/q_N$: the biomass that the individual's total nitrogen supply ($N_{tot}$, amino-acid nitrogen plus free nitrogen) could support if carbon and ATP were never limiting. I compute $M_N$ twice per individual – including and excluding the diet-dependent urea contribution to $N_{other}$ – and compare each to that individual's own $M_{max}$. Without urea, $M_N$ falls below the individual's LP-derived joint C–N ceiling for 18.8% of the NHANES cohort (i.e., nitrogen would be the tighter constraint for close to a fifth of individuals absent host urea recycling); with urea included, $M_N$ exceeds $M_{max}$ for every individual in the cohort (0%), so nitrogen is never the binding constraint once urea recycling is accounted for (Figure 4B,C).

**Carbon versus phosphorus limitation across individuals** Colon-reaching phosphorus is estimated for each individual from dietary phosphorus and calcium intake using a calcium-dependent regression fit to metabolic-ward balance-study data (Heaney & Nordin 2002), chosen as the most mechanistically grounded of three candidate mappings considered (a flat population-wide absorption fraction and a plant/non-plant-tiered fraction both agree closely with it; Figure S3). This colon-reaching phosphorus flux is converted to a phosphorus-only biomass ceiling, $M_P = P_{colon} \times Y_P$, using a fixed biomass phosphorus yield $Y_P = 40$ g dry weight per g P (equivalent to 2.5% phosphorus content of bacterial dry mass; Table S4). For each individual, I compare $M_P$ against that same individual's LP-derived joint C–N ceiling $M_{max}$ (Section 11 above) and classify the individual as phosphorus-limited if $M_P < M_{max}$, and carbon-limited otherwise (amino-acid limitation, the LP's third possible binding resource, occurs in $< 0.1\%$ of individuals and is not shown separately). Across the NHANES cohort, this yields 92.4% of individuals classified as carbon-limited and 7.6% as phosphorus-limited (Figure 4D,E), conditional on the assumption of 100% phosphorus accessibility ($a_P = 1$; Section 9.5). Figure S4E extends this to a joint sensitivity analysis over both the assumed yield and accessibility simultaneously.

Because the assumed phosphorus content of bacterial dry mass is itself a physiologically variable quantity (cellular phosphorus scales with RNA content, which in turn scales with growth rate), I repeated this classification while sweeping $Y_P$ across the full range of literature- reported biomass phosphorus yields (20–100 g dry weight per g P, i.e. 1–5% phosphorus content; Table S4), recomputing $M_P$ and the resulting phosphorus-limited population fraction at each value while holding each individual's own phosphorus and carbon influx fixed. The phosphorus-limited fraction is highly sensitive to this single parameter, ranging from $\sim$69% at the low-yield/high-phosphorus-content end of the swept range down to effectively 0% at the high-yield/low-phosphorus-content end; Figure 4F displays this sweep over the sub-range 20–65 for visual clarity (the curve is already flat near zero well before the full range's upper end), while the percentages above describe the complete computed 20–100 sweep.

More generally, the phosphorus-supported ceiling is

$$M_{P,i} = P_{\mathrm{colon},i}\, a_P\, Y_{X/P}. \tag{30}$$

### 13 Functional requirements implied by the resource model

#### 13.1 Requirements for de novo amino-acid synthesis and urease activity

For each NHANES individual, I use that individual's own carbon-supported biomass ceiling, $M_{max}$ (Section 12), to define a fixed anabolic nitrogen demand, $q_N M_{max}$: the total nitrogen required to build that much biomass, regardless of where the nitrogen ultimately comes from. This demand can be met by two routes – directly importing pre-formed dietary/endogenous amino acids (requiring no biosynthetic capability beyond uptake and, where relevant, extracellular proteolysis), or synthesizing amino acids de novo from free nitrogen (urea- and ammonium-derived) plus a carbon skeleton. The resource-limitation LP's own optimal allocation for each individual reports $N_{other}^{assim}$, the amount of free nitrogen actually drawn upon to meet that individual's nitrogen demand alongside directly-supplied amino-acid nitrogen. Because every unit of biomass nitrogen in this model is amino-acid nitrogen either way, the fraction

$$f_{AA} = \frac{N_{other}^{assim}}{q_N M_{max}}$$

is equivalently the fraction of an individual's total nitrogen demand that must come via free-nitrogen assimilation (i.e., de novo amino-acid synthesis) and the fraction of that individual's cellular amino-acid pool that cannot simply be taken up preformed. An individual requires de novo amino-acid synthesis at all whenever $f_{AA} > 0$; this occurs in 20.1% of the NHANES cohort (Figure 5B,C). Among individuals for whom synthesis is required, $f_{AA}$ has a median of 10.2% (interquartile range 5.0–18.6%; Figure 5D).

For each individual, urease necessity is assessed by two otherwise identical solves. The complete solve includes nitrogen from both host-secreted urea and ileal urea carryover. In the counterfactual solve, all urea-derived nitrogen is removed while preformed amino-acid nitrogen, ileal ammonium, and accessible non-urea nitrogen are retained. Urease-dependent nitrogen is required to attain the complete-model ceiling whenever

$$M_{max,i}^{\text{no urea}} < M_{max,i}^{\text{with urea}}. \tag{31}$$

This counterfactual definition tests whether urea availability changes the supported biomass ceiling and avoids assigning an arbitrary source composition to the pooled variable $N_{other}^{assim}$.

Urease is required in 18.7% of the NHANES cohort (Figure 5C); among these individuals, $f_{urease}$ has a median of 10.2% (interquartile range 4.8–18.1%; Figure 5E). Because urease-derived nitrogen is one specific source within the broader free-nitrogen pool that also supports de novo amino-acid synthesis, an individual who requires urease activity always also requires de novo amino-acid synthesis (though the converse need not hold): the population requiring urease (18.7%) is accordingly a subset of the population requiring de novo amino-acid synthesis (20.1%).

#### 13.2 Illustrative micronutrient case: corrinoid requirements

Over the last decade, vitamin $B_{12}$ (cobalamin) has been studied as an important factor shaping gut microbial communities. Comparative genomic analyses have shown that complete de novo corrinoid biosynthesis is restricted to a minority of gut bacteria, whereas many more species encode corrinoid-dependent enzymes together with transport and remodeling pathways. Work by Degnan and colleagues demonstrated that different bacterial species distinguish among structurally distinct corrinoids using multiple transporter systems, and that transporter specificity directly affects bacterial fitness and competition in the mammalian gut [48, 49]. Together, these studies establish corrinoids as important mediators of microbial interactions, cross-feeding, and community assembly.

**Lower-bound estimate of the cobalamin requirement for bacterial biomass production** I focus on cobalamin among the many micronutrients and cofactors gut bacteria require because it is the one case where a quantitative per-cell demand has been directly measured in pure culture (25–500 molecules/cell, below) and where the comparative-genomics and cross-feeding literature above already establishes its ecological importance; comparable quantitative demand estimates are not readily available for other vitamin/cofactor auxotrophies, which is why they are not quantified here. Unlike the C/N/P/S/Mg calculation above, I present this estimate as a fully worked numeric example rather than deferring the final value to the main text, since it serves primarily to illustrate the scale of cobalamin demand relative to the dietary influx estimated in Section 2.5.1 rather than to feed directly into the multi-element supply-demand figure. The underlying parameters (molecules required per cell, representative cell dry mass, assumed daily bacterial biomass production) are listed in `data/growth_demand_parameters.csv` alongside the other demand parameters, so that this calculation can be recomputed consistently if any of these inputs are revised.

To estimate the minimum amount of cobalamin required to support bacterial growth, I use classical measurements reporting that approximately 25–500 cobalamin molecules per cell are sufficient to support maximal growth of auxotrophic bacteria in pure culture. Using a representative bacterial dry mass of 285 fg/cell (the same X-ray microanalytic measurement, early exponential phase, used for the Mg/K/Ca/Na biomass yields above [172]), one gram of bacterial dry biomass contains approximately $3.51 \times 10^{12}$ cells. Multiplying by the per-cell requirement and converting from molecule count to mass via Avogadro's number and the molecular mass of cobalamin ($M_{\mathrm{Cbl}} = 1355$ g/mol) gives a cobalamin requirement of 0.20–3.95 $\mu$g per gram bacterial dry biomass across the 25–500 molecule/cell range. Multiplying by a representative daily community bacterial dry-biomass production of 12.31 g/day (the median $M_{max}$ across the NHANES cohort from the carbon/nitrogen growth-limitation model above) gives an implied daily cobalamin requirement of 2.4–48.6 $\mu$g/day.

Comparing this against median dietary vitamin $B_{12}$ intake (3.37 $\mu$g/day; Section 2.5.1) – used here as an upper-bound proxy for colon-reaching supply, since a resolved colon-reaching/accessibility mapping for B12 (accounting for ileal absorption, host secretion, microbial production, corrinoid identity, and accessibility) has not been developed – dietary supply modestly exceeds the lower-bound demand at the low end of the per-cell requirement range (supply/demand $\approx 1.4$), but covers only around 7% of the demand implied by the high end of that range (supply/demand $\approx 0.07$, i.e. demand $\approx 14\times$ supply).

This calculation should be interpreted as a lower-bound estimate of the cobalamin-equivalent requirement for bacterial biomass production. The underlying estimate of 25–500 molecules per cell originates from classical studies of cobalamin uptake in pure culture and may not represent the total intracellular corrinoid content of bacterial cells under gut conditions. Gut bacteria frequently utilize a diversity of cobamides, may store or remodel corrinoids, and often contain additional corrinoid intermediates or analogues; consequently, the total corrinoid requirement of the gut microbiota could be substantially higher than this cobalamin-specific lower bound. Revisiting the original pure-culture studies and extending such measurements to representative gut auxotrophs will be important for establishing more accurate quantitative estimates of corrinoid demand. Notably, at the high end of the classical per-cell requirement range, dietary B12 alone could not plausibly account for community-wide biomass-linked cobalamin demand – consistent with the independent evidence (Figure 5G) that fecal non-B12 corrinoid analogues substantially exceed dietary and fecal B12 itself, pointing to substantial de novo microbial corrinoid synthesis.

### 14 Summary of robust and assumption-dependent conclusions

**Conclusions expected to be comparatively robust** Fermentative energy generation substantially increases carbon consumption relative to the carbon incorporated into biomass. Host urea recycling can materially expand the nitrogen supply available to the microbial community. The reconstructed bulk supplies of Na, K, Ca, and Mg appear large relative to representative cellular ion inventories, although these comparisons are sufficiency checks rather than strict stoichiometric growth limits.

**Conclusions conditional on model assumptions** The fraction of the population assigned a phosphorus-binding constraint depends strongly on both cellular phosphorus quota and the unknown accessibility of colon-reaching phosphorus (Figure S4E quantifies this joint sensitivity directly). The fraction requiring urease activity (18.7%) reflects the corrected classification above, in which both host-secreted and ileal- carry-over urea are treated as urease-dependent. The inferred need for de novo amino-acid synthesis depends on the modeled accessibility of preformed amino acids and the carbon–nitrogen allocation constraints.

**Important unresolved quantities** The principal unresolved quantities include microbial phosphorus accessibility, allocation of anabolic carbon among substrate classes, taxon-specific and spatial access to resources, and quantitative corrinoid requirements under gut conditions.

### 15 Supplementary Figures

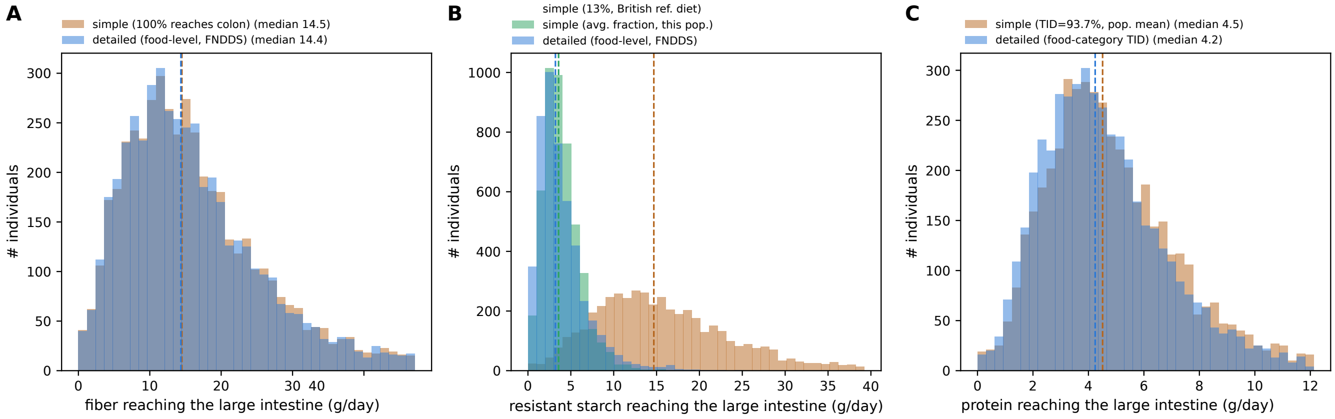

**Figure S1: Food-level mapping from dietary intake to large-intestinal resource influx, validated against simple flat-fraction approaches for three dietary resources.** In this work I introduce an estimate of resource influxes into the large intestine using a detailed, FNDDS-resolved mapping that accounts for food-specific composition and/or digestibility at the level of individual foods consumed, rather than applying a single population-average escape or digestibility fraction to total reported intake. This approach allows influx estimates to reflect not only how much of a given macronutrient class an individual consumes, but which specific foods it comes from and how efficiently each is digested or absorbed. To assess how this food-level resolution compares to simpler approach, I compare my detailed mapping against a flat-fraction mapping, applied uniformly to total reported intake, for three representative dietary resources. (A) Dietary fiber reaching the colon: both mappings assume complete escape from small-intestinal digestion (Section 2.1.2), so this isolates whether food-level resolution adds information beyond total reported fiber intake. The two agree closely (person-level $r = 1.00$), confirming that it does not, as expected given the shared assumption of complete escape. (B) Resistant starch reaching the colon: my detailed, food-level mapping (Section 2.1.1) is compared against two simple constant-fraction approaches – a historically calibrated 13% escape fraction and a constant fraction recalibrated to this population's own detailed-mapping-implied average. The fixed 13% comparator overestimates typical delivery roughly 4-fold (median 14.7 vs. 3.2 g/day); the recalibrated constant tracks the median closely (3.6 vs. 3.2 g/day) but correlates only moderately at the individual level ($r = 0.72$), since a single population-wide fraction cannot capture person-to-person variation in which starch-rich foods are consumed. (C) Dietary amino acids/protein reaching the colon: a simple mapping using a constant, population-mean true ileal digestibility (TID $= 93.7\%$) is compared against my detailed, food-category-specific mapping (Section 2.2). The two distributions are similar in central tendency (median 4.5 vs. 4.2 g/day) but only moderately correlated person-to-person ($r = 0.75$), indicating that individual protein-source composition materially affects colon-reaching amino-acid supply. Together, these comparisons show that food-level resolution is essential for capturing realistic person-to-person variation in resource influx, even when it adds little beyond a population-average estimate for resources, such as fiber, where digestion assumptions are already uniform across foods. Dashed vertical lines mark each distribution's median. Numeric summary statistics: medians, 5th–95th percentiles.

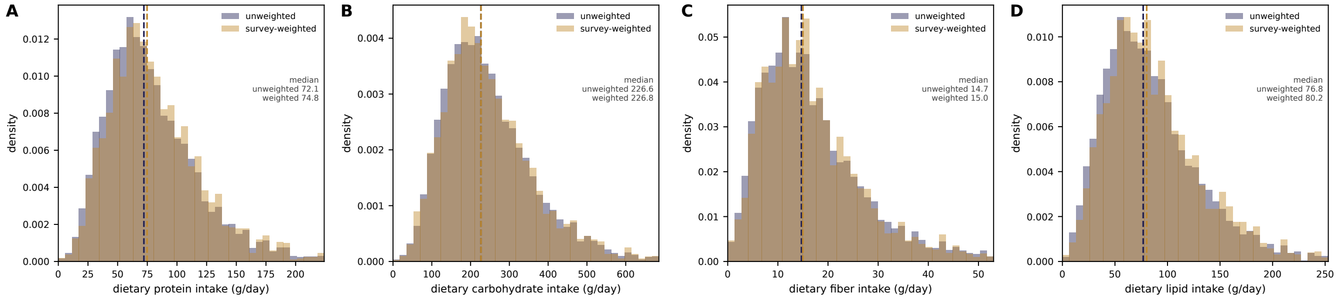

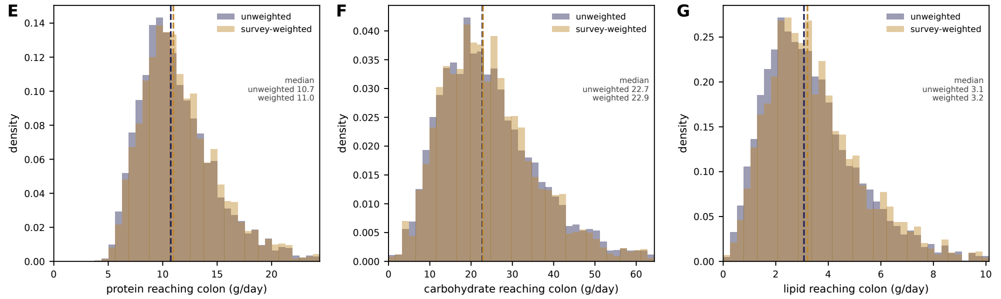

**H**

*mean ± SD, by quantity:*

| Quantity | Unweighted | Survey-weighted |
|---|---|---|
| Dietary protein (g/day) | 80.3±43.1 | 82.7±42.1 |
| Dietary carbohydrate (g/day) | 250.1±128.4 | 250.4±126.3 |
| Dietary fiber (g/day) | 17.0±11.0 | 17.0±10.6 |
| Dietary lipid (g/day) | 86.0±49.9 | 89.1±49.5 |
| Protein reaching colon (g/day) | 11.5±3.9 | 11.7±3.8 |
| Carbohydrate reaching colon (g/day) | 24.9±12.9 | 24.8±12.6 |
| Lipid reaching colon (g/day) | 3.4±2.0 | 3.6±2.0 |

**Figure S2: Sensitivity to NHANES dietary sample weighting.** NHANES uses a complex, multistage sampling design that deliberately oversamples specific population subgroups (e.g. by age, income, or race/ethnicity) to obtain adequate statistical power for analyzing them individually; NHANES therefore assigns each respondent a sample weight (for the day-1 data I use, the weight `WTDRD1`), roughly inverse to that respondent's probability of selection, so that a *survey-weighted* statistic approximates a nationally representative estimate of the U.S. population. All analyses elsewhere in this work are instead unweighted: every individual in the analyzed NHANES sample is given equal weight regardless of their selection probability, so the resulting distributions describe the analyzed sample itself rather than a nationally representative projection. This distinction could be important for my conclusions if differences in sampling probabilities are associated with systematic differences in diet or reconstructed resource influxes. Recomputing the key dietary and colon-reaching distributions from the same sample using the day-1 dietary sample weight shows that weighting has little effect on the dietary intake and reconstructed resource influx distributions considered here: applying survey weights shifts these distributions only modestly. (A–D) Dietary protein, carbohydrate, fiber, and lipid intake, unweighted vs. survey-weighted, each with its own median marked. (E–G) The corresponding colon-reaching quantities for protein, carbohydrate, and lipid, unweighted vs. survey-weighted. (H) Mean $\pm$ SD, unweighted vs. survey-weighted, for each of the seven quantities shown in (A–G), demonstrating that these summary statistics are also only modestly affected by reweighting. Given the modest effects of survey weighting observed here, I use the unweighted NHANES sample for the other analyses in this work.

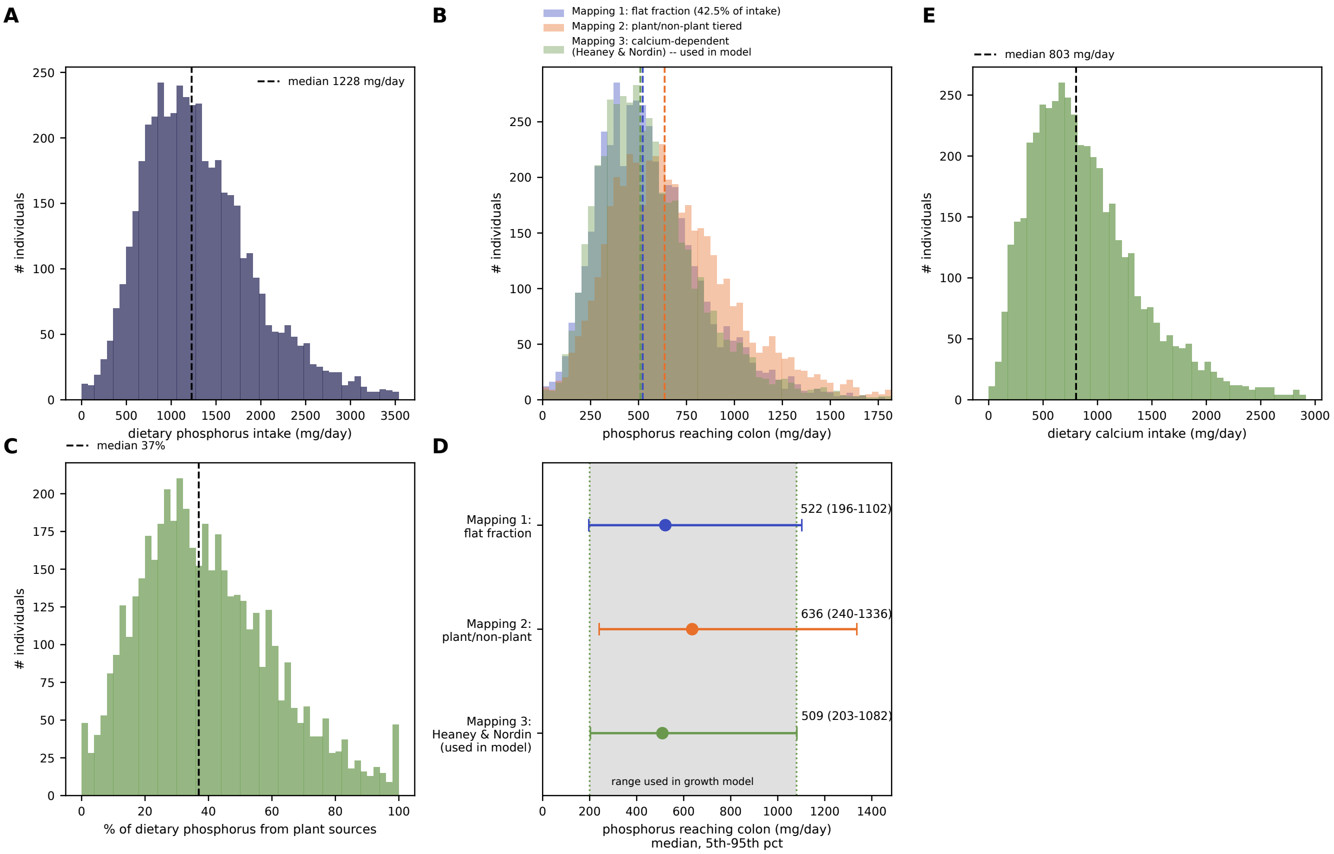

**Figure S3: Dietary phosphorus intake and three literature-grounded mappings to colon-reaching phosphorus (Section 4.3).** To estimate phosphorus influx into the large intestine, I build on measurements of total phosphorus in the diet and ileal effluent rather than estimating and summing the phosphorus content of individual resources. Phosphorus sources are diverse, including nucleic acids and plant-based components for which resource-specific influxes are poorly constrained, whereas total phosphorus intake and ileal delivery can be constrained more directly from available measurements (SI Text 4.3). Here, I show the distributions obtained by applying the mapping I derived to the NHANES dietary data. (A) Dietary phosphorus intake across NHANES adults (median 1228 mg/day). (B) Colon-reaching phosphorus under the three candidate mappings: a flat absorption fraction (Mapping 1), a plant/non-plant-tiered fraction (Mapping 2), and a calcium-dependent regression fit to metabolic-ward balance-study data (Mapping 3, Heaney & Nordin 2002) – the standard mapping I used throughout this work. The shaded band marks Mapping 3's 5th–95th percentile range (200–1080 mg/day), the range used in the growth limitation model. (C) Percent of dietary phosphorus from plant sources (median 36.9%), the tiering variable behind Mapping 2. (D) The three mappings' medians (5th–95th percentile) for direct comparison; the medians agree to within about 25% of one another despite relying on different absorption assumptions. (E) Dietary calcium intake (median 803 mg/day), the second input to Mapping 3's regression.

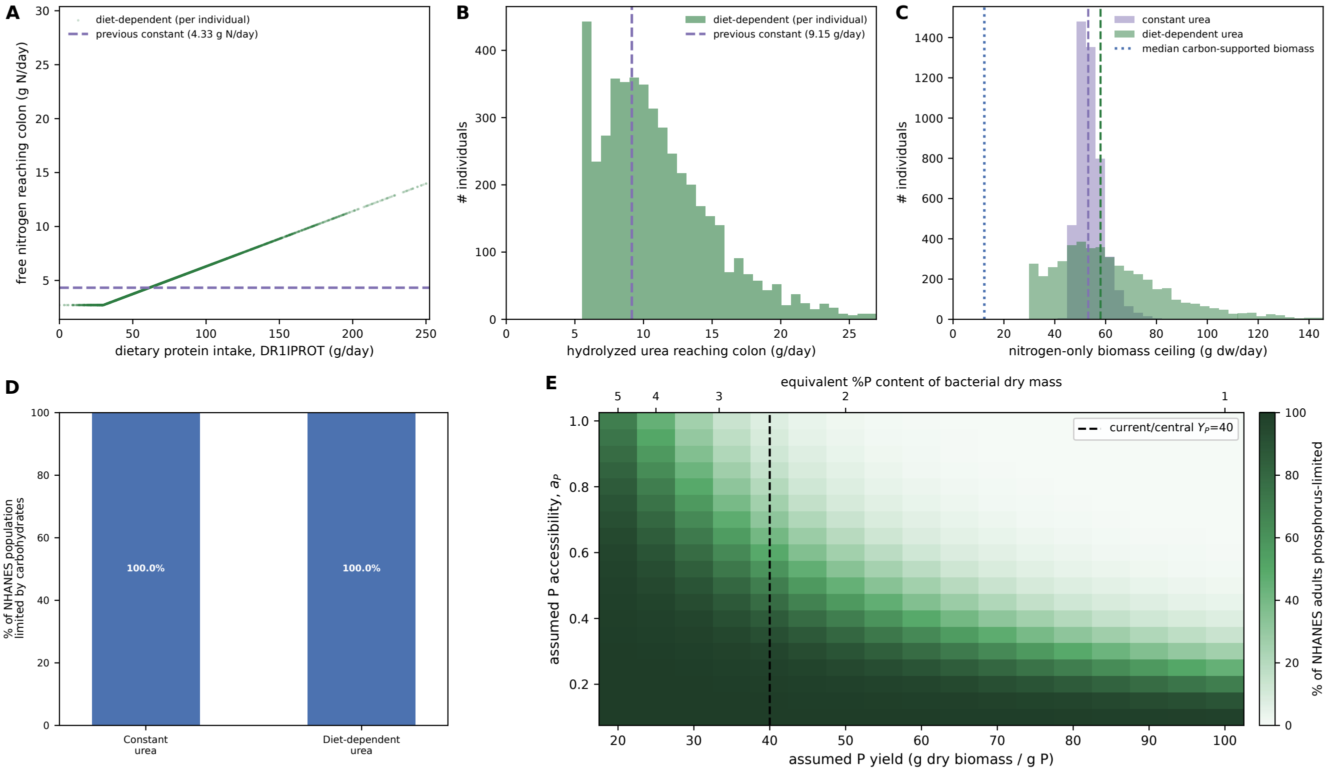

**Figure S4: Sensitivity of the nitrogen and phosphorus limitation conclusions to modeling assumptions.** (A–D) Sensitivity of the nitrogen-limitation result to diet-dependent urea supply. Because hepatic urea production depends in part on the disposal of excess amino-acid nitrogen, urea delivery to the large intestine can strongly depend on dietary protein intake. To probe whether this dependence on diet affects the overall picture of nitrogen supply and supported growth in the gut, I also considered a mapping in which urea influx into the large intestine is coupled to protein consumption, as described in SI Text 6.2 (paragraph "Modeling urea influx into the large intestine"). Here, I show the modeling results and compare them with a model assuming a constant urea influx. (A) Free nitrogen reaching the colon under the diet-dependent model as a function of individual protein intake (`DR1IPROT`), compared with the population-wide constant used elsewhere (dashed line). For most individuals, the diet-dependent model predicts greater urea nitrogen delivery than the population-wide constant. (B) The corresponding population-level distribution of hydrolyzed urea flux, showing the same comparison as in (A) as a distribution rather than resolved against protein intake. (C) Population distribution of the nitrogen-only biomass ceiling considering the total nitrogen amount reaching the large intestine, comparing the constant-urea and diet-dependent-urea models. Every individual receives the same urea contribution under the constant model, so the distribution's spread reflects only variation in amino-acid nitrogen supply; the diet-dependent model additionally varies the urea contribution with individual protein intake, widening the population-level spread. The median carbon-supported biomass ceiling (dotted line) lies well below both nitrogen-ceiling distributions. (D) The growth-limiting resource, re-solved for each individual with the constant vs. diet-dependent urea supply. In both scenarios carbohydrates are growth limiting, showing that the conclusion that nitrogen is not a primary growth constraint is robust to diet-dependent urea supply, at least for the typical protein consumed rates of the US population probed here. (E) Joint sensitivity of phosphorus limitation to the assumed phosphorus biomass yield $Y_P$ and accessibility $a_P$ (Section 9.5), extending Figure 4F's one-dimensional $Y_P$ sweep (at $a_P = 1$) to the full $Y_P$ (20–100 g dry biomass/g P) $\times$ $a_P$ (0.1–1.0) grid, with each individual's phosphorus and carbon influx held fixed. Here, $a_P$ rescales the phosphorus-supported biomass ceiling but not the carbon- or nitrogen-supported biomass ceiling. The reference scenario ($Y_P = 40$, dashed line; $a_P = 1$) sits at the edge of the near-zero region, so a lower yield or lower accessibility alone can substantially increase the phosphorus-limited fraction, reaching effectively the whole cohort at the lowest yield and accessibility considered ($Y_P = 20$, $a_P = 0.1$). Overall, this analysis emphasizes that a systematic investigation of phosphorus demand by gut bacteria is an important next step.

### 16 Supplementary Tables

**Table S1.** Summary of estimated resource influxes into the human large intestine. For each resource discussed in this Supplementary Information, I report the route of entry (dietary, host-derived via the ileum, host-derived across the large-intestinal epithelium, terminal electron acceptor, or major ion/metal), a central (single-value) estimate and, where available, a range of the daily influx, its qualitative dependence on diet, the assigned confidence level and the type(s) of evidence underlying the estimate (Table S2), and the corresponding Supplementary Text section in which the estimate is derived and justified. The central estimate is the single working value used in the model wherever influx is not modeled with population variation; the range reflects the estimated spread of the influx, either across the US population or across the literature source studies, depending on the resource. The additional *central-value basis* column indicates which of these applies. Where a resource is mapped from NHANES dietary recall data using the mapping I developed for this work (SI Text 8), the central estimate is the population median and the range is the 5th–95th percentile of the resulting population distribution.

| Category | Resource | Route | Central estimate | Range | Diet dependence | Confidence | Central-value basis | Evidence type | SI section |
|---|---|---|---|---|---|---|---|---|---|
| Dietary carbohydrate | Resistant starch | Ileum | 3.2 | 0.8-9.4 g/day | Strong | Good | NHANES population median | IL, FC, DV | Sec. 2.1.1 |
| Dietary carbohydrate | Dietary fiber | Ileum | 14.4 | 4-37 g/day | Strong | Good | NHANES population median | IL, FC, DV | Sec. 2.1.2 |
| Dietary carbohydrate | Free glucose and other soluble sugars | Ileum | 2.5 | 1.0-5.0 g/day | Strong (via starch and added-sugar/fruit intake) | Low | Combined literature estimate: starch-derived glucose/maltose plus an inferred fructose contribution | IL, DV, PI | Sec. 2.1.3 |
| Dietary protein | Dietary amino acids/ protein escaping absorption | Ileum | 4.2 | 1.5-9.3 g/day | Strong | High | NHANES population median | IL, FC, DV, TR, PI | Sec. 2.2 |
| Dietary lipid | Dietary lipids | Ileum | 3.1 | 1-7 g/day | Strong (fat/ fiber intake) | Moderate | Literature central estimate (point value, not distribution-derived) | IL, DV, FO, PI | Sec. 2.3 |
| Dietary nucleic acid | Nucleic acids and nucleotides | Ileum | | Not quantified g/ day | Unknown | Unknown | N/A (not quantified) | | Sec. 2.4 |
| Dietary vitamin | Cobalamin (vitamin B12) | Ileum | 3.4 | 0.5-12.5 ug/day | Strong (diet/ supplementation) | Moderate | NHANES population median | PI, DV | Sec. 2.5.1 |
| Dietary vitamin | Non-cobalamin corrinoids | Ileum | | Not quantified ug/ day | | Unknown | N/A (not quantified) | | Sec. 2.5.1 |
| Dietary vitamin | Other B-vitamins and heme | Ileum | | Not quantified ug/ day | | Unknown | N/A (not quantified) | | Sec. 2.5.2 |

| Category | Resource | Route | Central estimate | Range | Diet dependence | Confidence | Central-value basis | Evidence type | SI section |
|---|---|---|---|---|---|---|---|---|---|
| Dietary polyphenol | Polyphenols | Ileum | 0.8 | 0.3-0.9 g/day | Strong (intake) but escape fraction poorly constrained | Low | Literature central estimate (point value, not distribution-derived) | IL, DV, PI | Sec. 2.6 |
| Host-derived (ileum) | Mucin entering from ileum | Ileum | 4.0 | 1.9-18.8 g/day | Not established | Low | Literature central estimate (point value, not distribution-derived) | IL, PI | Sec. 3.1 |
| Host-derived (ileum) | Endogenous amino acids | Ileum | 3.0 | 3 g/day | Not established | Moderate | Literature central estimate (point value, not distribution-derived) | IL, DV | Sec. 3.2 |
| Host-derived (ileum) | Bile acids | Ileum | 0.7 | 0.4-1 g/day | Strong (dietary fat) | Moderate | Literature central estimate (point value, not distribution-derived) | IL, FO, DV, PI | Sec. 3.3 |
| Host-derived (ileum) | Urea (ileal efflux) | Ileum | 1.2 | 0.5-1.9 mmol/day | Not established | Moderate | Literature central estimate (point value, not distribution-derived) | IL, DV | Sec. 3.4 |
| Host-derived (ileum) | Ammonium (ileal efflux) | Ileum | 0.9 | 0.6-1.2 mmol/day | Not established | Moderate | Literature central estimate (point value, not distribution-derived) | IL, DV | Sec. 3.4 |

| Category | Resource | Route | Central estimate | Range | Diet dependence | Confidence | Central-value basis | Evidence type | SI section |
|---|---|---|---|---|---|---|---|---|---|
| Host-derived (epithelium) | Urea (secreted and hydrolyzed in situ) | Large-intestinal epithelium | 9.15 | 5.4-12.9 g/day | Strong (dietary protein; 30-70 g/day range) | Good | Literature central estimate (point value, not distribution-derived) | TR, DV, (FO), PI | Sec. 6.2 |
| Host-derived (epithelium) | Bicarbonate (host secretion) | Large-intestinal epithelium | | 10-300 mmol/day | Coupled to SCFA/ fermentation activity | Unknown | N/A (no central value; intentionally broad order-of-magnitude range only) | DI, PI | Sec. 6.3 |
| Host-derived (epithelium) | Mucus/epithelial mucin release (local) | Large-intestinal epithelium | 0.4 | 0.4 g/day | Not modeled | Low | Literature central estimate (point value, not distribution-derived) | PI, TR | Sec. 6.4 |
| Host-derived (epithelium) | Epithelial cell shedding | Large-intestinal epithelium | | 1.5-3.5 g/day | Not modeled | Low | Literature central estimate (point value, not distribution-derived) | PI, TR | Sec. 6.5 |
| Terminal electron acceptor | Oxygen | Ileum/ epithelium | 1.5 | 0.5-2.5 combined mmol/day | Not modeled | Unknown | Range midpoint (no independent central estimate reported) | DI, PI | Sec. 7.1 |
| Terminal electron acceptor | Nitrate | Ileum | 0.155 | 0.01-0.3 mmol/day | Strong (leafy vegetable intake) | Moderate | Range midpoint (no independent central estimate reported) | IL, DV, PI | Sec. 7.2 |
| Terminal electron acceptor | Sulfate (free) | Ileum | 6.5 | 1-12 mmol/day | Strong (intake up to absorption plateau ~5 mmol/day) | Moderate | Range midpoint (no independent central estimate reported) | IL, DV, (FO) | Sec. 7.3 |

| Category | Resource | Route | Central estimate | Range | Diet dependence | Confidence | Central-value basis | Evidence type | SI section |
|---|---|---|---|---|---|---|---|---|---|
| Terminal electron acceptor | Fumarate | Large intestine (largely microbial) | | Not quantified - | Not established | Unknown | N/A (not quantified) | | Sec. 7.4 |
| Terminal electron acceptor | TMAO | Ileum | | Not quantified - | Dietary (fish/ seafood) | Unknown | N/A (not quantified) | | Sec. 7.4 |
| Terminal electron acceptor | DMSO | Ileum | | Not quantified - | Unknown | Unknown | N/A (not quantified) | | Sec. 7.4 |
| Major ion | Potassium | Ileum | 0.55 | 0.2-0.9 g/day (6-22 mmol/day) | Not established | Moderate | Literature central estimate (point value, not distribution-derived) | IL, FO | Sec. 5.1 |
| Major ion | Sodium | Ileum | 3.15 | 2.8-3.5 g/day (120-150 mmol/ day) | Not established | Moderate | Literature central estimate (point value, not distribution-derived) | IL, FO | Sec. 5.2 |
| Major ion | Calcium | Ileum | 0.8 | 0.8 g/day (20 mmol/day) | Diet (fiber/ pectin) may modulate | Moderate | Literature central estimate (point value, not distribution-derived) | IL, FO | Sec. 5.3 |
| Major ion | Magnesium | Ileum | 0.12 | 0.12 g/day (5 mmol/day) | Not established | Moderate | Literature central estimate (point value, not distribution-derived) | IL, FO | Sec. 5.4 |
| Metal/mineral | Iron | Ileum | 8.0 | 8 mg/day | Not established | Low | Literature central estimate (single direct ileostomy measurement) | IL | Sec. 4.1 |

| Category | Resource | Route | Central estimate | Range | Diet dependence | Confidence | Central-value basis | Evidence type | SI section |
|---|---|---|---|---|---|---|---|---|---|
| Metal/mineral | Zinc | Ileum | 10.0 | 5-13 mg/day | Strong (phytate reduces bioavailability) | Moderate | Literature central estimate (point value, not distribution-derived) | IL, FO | Sec. 4.2 |
| Metal/mineral | Phosphorus | Ileum | 0.51 | 0.20-1.08 g/day | Moderate (plant/ animal/ additive composition) | Moderate | NHANES population median | IL, FO, FC, DV, PI | Sec. 4.3 |

**Table S2.** Evidence types and quantitative confidence levels. Evidence types: I characterize the available studies to support resource flux estimations into several major evidence types defined here. Confidence levels: To assign to each resource influx a quantitative confidence level. The confidence level reflects the strength of the quantitative evidence supporting the estimated influx into the large intestine rather than the biological importance of the resource itself. Evidence types and confidence levels for all major resources analyzed in this study are provided in Table S1 and Figure 1A. The choice of confidence levels is motivated in the corresponding supplementary text section of each resource together with the studies and evidence supporting the estimate (SI Text 2– 7).

| **Evidence types** | | |
|---|---|---|
| DI | Direct intestinal measurements | Resource influx directly quantified or constrained from measurements in the intact human intestinal tract. |
| IL | Ileostomy measurements | Resource output measured in ileostomy effluent, providing a direct estimate of material escaping small-intestinal digestion and absorption. |
| TR | Isotope/tracer measurements | Resource absorption, secretion, turnover, or recovery quantified using isotope or other tracer approaches. |
| FO | Fecal output / mass balance | Daily fecal output or physiological mass-balance measurements used to constrain resource influx. |
| DV | Dietary variation/ intervention | Resource delivery or output quantified across different dietary intakes or controlled dietary interventions. |
| FC | Food-specific characterization | Resource composition, digestibility, or absorption characterized across different foods or nutrient sources. |
| PI | Physiological inference | Resource influx inferred by quantitatively combining physiological measurements or established physiological parameters. |
| **Confidence levels** | | |
| High | | Direct quantitative measurements are available from multiple independent studies, the underlying physiology is well understood, and major sources of variation (e.g. diet, food processing, or physiology) have been systematically characterized. |
| Good | | Direct measurements exist and are supported by a good mechanistic understanding, but systematic characterization of variation or independent validation remains less systematic. Overall quantitative estimates are considered reliable, although uncertainty remains regarding specific conditions and dependence on diet and digestion. |

| | |
|---|---|
| Moderate | Direct measurements are available to support the estimation. Mechanistic is partly understood. The overall magnitude is very likely correct, but uncertainty remains substantial owing to incomplete measurements or simplifying assumptions. |
| Low | Few or no direct measurements exist. Quantitative estimates rely primarily on indirect evidence, related physiological measurements, or biophysical constraints. Order-of-magnitude estimates are possible, but remain weakly constrained. |
| Unknown | No meaningful quantitative estimate currently exists. Resource influx has not been measured directly and cannot presently be constrained beyond qualitative reasoning or rough theoretical bounds. |

**Table S3.** Conversion factors used to translate substrate-level influx estimates into elemental fluxes of carbon, nitrogen, phosphorus, and sulfur, together with the microbial utilization fraction assumed for each. Conversion factor: For each contributing substrate, I list the assumed elemental mass fraction (or molar conversion, where applicable), the basis for this value (exact molecular stoichiometry, representative compositional data, or direct biochemical measurement), and the Supplementary Text section in which this conversion factor is discussed (Section 9). Utilizable fraction: The utilizable fraction is the fraction of each resource (or resource portion) reaching the large intestine that is assumed accessible to microbial fermentation/assimilation in the growth-limitation model; it is applied uniformly across every element of a given resource, since it reflects assumed availability of the entire resource, rather than a directly measured, element-specific assimilation efficiency. For the breakdown of fiber into individual fiber-type breakdown (cellulose, hemicellulose, beta-glucan, pectin, inulin/FOS, GOS, and lignin), I take the same elemental conversion factor as overall fiber. Some resources have 0 utilizable fraction assigned outright as these resources are of very limited use to support growth in the conditions bacteria encounter in the gut, see SI Text 9.5.

| Element | Substrate | Conversion factor | Conversion type / basis | Utilizable fraction | Utilization basis |
|---|---|---|---|---|---|
| Carbon | Resistant starch | 0.44 | mass fraction | 100% | preferred, essentially completely fermented substrate |
| Carbon | Free glucose and other soluble sugars | 0.4 | mass fraction | 100% | readily and essentially completely fermented |
| Carbon | Dietary fiber | 0.40-0.45 | mass fraction (range reflects general uncertainty in mixed-polysaccharide C content, not a subtype-weighted average – see note) | 50% | single working value – mixed fiber types ferment very differently, see below |
| Carbon | Protein and amino acids (dietary + endogenous) | 0.53 | mass fraction | 100% | assumed all protein/AA-derived material is microbially utilizable |
| Carbon | Lipids | 0.76 | mass fraction | 0% | not a bacterial fermentation substrate in the colon |
| Carbon | Mucin (whole) | 0.482 (= 0.376 glycan portion, 80% of mass + 0.106 protein-backbone portion, 20% of mass) | mass fraction, split by macro-component in the model | 100% | 100% for both portions: protein/AA rule (protein-backbone portion); specialized mucin-degrading taxa present, though not all mucin turned over microbially (glycan portion) |
| Carbon | Polyphenols | 0.60-0.65 | mass fraction (range reflects general uncertainty across flavonoid/phenolic-acid formulae) | 30% | partially ring-cleaved/co-metabolized; much passes through largely intact |
| Carbon | Bile acids (unconjugated) | 0.70 | mass fraction | 100% | deconjugated/dehydroxylated (biotransformed) rather than assimilated as a carbon source |
| Carbon | Bile acids (taurine-conjugated) | 0.60 | mass fraction | 100% | deconjugated/dehydroxylated (biotransformed) rather than assimilated as a carbon source |
| Carbon | Epithelial cell shedding - protein portion | 0.265 | mass fraction (50% of dry mass x 53% C) | 100% | protein/AA rule |
| Carbon | Epithelial cell shedding - glycan/glycoprotein portion | 0.0705 | mass fraction (15% of dry mass x 47% C, mucin-glycan approximation) | 40-70% | first-pass estimate, not literature-sourced |
| Carbon | Epithelial cell shedding - nucleic-acid portion | 0.036 | mass fraction (10% of dry mass x 36% C) | 40-70% | first-pass estimate, not literature-sourced |

| Element | Substrate | Conversion factor | Conversion type / basis | Utilizable fraction | Utilization basis |
|---|---|---|---|---|---|
| Carbon | Epithelial cell shedding - membrane-lipid portion | 0.114 | mass fraction (15% of dry mass x 76% C) – tracked for completeness but not currently routed into the resource-limitation model, which has no lipid-carbon pool | 0% | not a colonic fermentation substrate |
| Nitrogen | Protein and amino acids (dietary + endogenous) | 0.16 | mass fraction | 100% | assumed all protein/AA-derived material is microbially utilizable |
| Nitrogen | Mucin (whole) | 0.064 (= 0.032 glycan portion, 80% of mass x 4% N + 0.032 protein-backbone portion, 20% of mass x 16% N) | mass fraction, split by macro-component in the model | 100% | 100% for both portions: protein/AA rule (protein-backbone portion); specialized mucin-degrading taxa present, though not all mucin turned over microbially (glycan portion) |
| Nitrogen | Urea | 0.47 | mass fraction | 100% | readily hydrolyzed by ureolytic bacteria; treated as fully microbially accessible once hydrolyzed (host reabsorption of the hydrolyzed nitrogen is modest, ~15%) |
| Nitrogen | Ammonium ($NH_4^+$) | 14 | mg N per mmol | 100% | directly assimilable nitrogen source for many taxa |
| Nitrogen | Nitrate ($NO_3^-$) | 14 | mg N per mmol | 70-100% | substrate for a minor but present nitrate-reducing niche |
| Nitrogen | Epithelial cell shedding - protein portion | 0.08 | mass fraction (50% of dry mass x 16% N) | 100% | protein/AA rule |
| Nitrogen | Epithelial cell shedding - glycan/glycoprotein portion | 0.006 | mass fraction (15% of dry mass x 4% N, mucin-glycan approximation) | 40-70% | first-pass estimate, not literature-sourced |
| Nitrogen | Epithelial cell shedding - nucleic-acid portion | 0.0155 | mass fraction (10% of dry mass x 15.5% N) | 40-70% | first-pass estimate, not literature-sourced |
| Nitrogen | Other N compounds (nucleic acids, amino sugars, polyamines) | | <5% of ileal influx. Not considered as an independent dietary/ileal source – NOTE: epithelial cell shedding's own nucleic-acid-N portion (row above) IS tracked, as it enters via a different resource, not this one | | not routed through the microbial-utilization/ accessibility step (see conversion_type note) |

| Element | Substrate | Conversion factor | Conversion type / basis | Utilizable fraction | Utilization basis |
|---|---|---|---|---|---|
| Sulfur | Dietary protein (Met + Cys residues) | 0.009 | mass fraction | 100% | assumed all protein/AA-derived material is microbially utilizable |
| Sulfur | Free sulfate ($SO_4^{2-}$) | 1 | mol S per mol sulfate (= 32 mg S per mmol substrate, atomic mass basis, as actually stored in the model) | 100% | assimilatory uptake for cysteine/methionine synthesis, a near-universal bacterial capability |
| Sulfur | Bile acids | 0.44-1.94 | mmol S/g bile acid, from taurocholic acid's S content (6.22% S by mass when taurine-conjugated) scaled by the estimated taurine-conjugated fraction of the human bile-acid pool (~20-25%; upper bound assumes 100% conjugated) | 80% | bile-salt hydrolase, a broadly conserved gut-bacterial enzyme |
| Sulfur | Mucin (whole) | 0.0044-0.0076 | mass fraction of TOTAL mucin mass, kept as one blended range (not split into protein-backbone vs. glycan sulfate esters) – this is how the model actually computes it, since S isn't routed through the C_CH/C_AA/ N_other allocation the way C and N are | 100% | 100% for both portions: protein/AA rule (protein-backbone portion); specialized mucin-degrading taxa present, though not all mucin turned over microbially (glycan portion) |
| Sulfur | Epithelial cell shedding - protein portion | 0.0045 | mass fraction (50% protein mass x 0.9% S/g protein, same basis as the dietary-protein S row above); other portions (glycan, nucleic-acid, lipid) not separately quantified for S | 100% | protein/AA rule |
| Magnesium | All sources (ileal/fecal flux) | 1 | based on measurement of total influx measurement (not distinguishing free and bound) | 100% | inorganic nutrients; fermentation-utilization framing doesn't apply – treated as fully bioavailable |
| Phosphorus | All sources (ileal/fecal flux) | 1 | based on measurement of total influx measurement (not distinguishing free and bound) | 100% | inorganic nutrients; fermentation-utilization framing doesn't apply – treated as fully bioavailable |
| Phosphorus | Epithelial cell shedding | 0.0095 | mass fraction, nucleic-acid fraction only (~10% of dry mass, ~9.5% P by mass); phospholipid P from the ~15% lipid fraction neglected as a simplification | 40-70% | first-pass estimate, not literature-sourced |
| K | All sources (ileal/fecal flux) | 1 | based on measurement of total influx measurement (not distinguishing free and bound) | 100% | inorganic nutrients; fermentation-utilization framing doesn't apply – treated as fully bioavailable |

| Element | Substrate | Conversion factor | Conversion type / basis | Utilizable fraction | Utilization basis |
|---|---|---|---|---|---|
| Na | All sources (ileal/fecal flux) | 1 | based on measurement of total influx measurement (not distinguishing free and bound) | 100% | inorganic nutrients; fermentation-utilization framing doesn't apply – treated as fully bioavailable |
| Ca | All sources (ileal/fecal flux) | 1 | based on measurement of total influx measurement (not distinguishing free and bound) | 100% | inorganic nutrients; fermentation-utilization framing doesn't apply – treated as fully bioavailable |
| Carbon | Dietary fiber - cellulose | | uses the blended Dietary fiber carbon factor above (0.40-0.45); listed here only for its fermentability-derived utilization fraction | 30% | least-fermented major fiber type in comparative in vitro fecal-fermentation studies, due to its crystalline structure (data/carbohydrate_fermentability_parameters.csv) |
| Carbon | Dietary fiber - hemicellulose/arabinoxylan | | uses the blended Dietary fiber carbon factor above (0.40-0.45); listed here only for its fermentability-derived utilization fraction | 60% | intermediate fermentability in comparative in vitro studies – fermented, but less completely/rapidly than pectin, beta-glucan, or inulin-type fibers |
| Carbon | Dietary fiber - beta-glucan | | uses the blended Dietary fiber carbon factor above (0.40-0.45); listed here only for its fermentability-derived utilization fraction | 90% | rapidly and almost completely fermented in comparative in vitro studies (e.g. oat beta-glucan vs. inulin/xylooligosaccharide comparisons) |
| Carbon | Dietary fiber - pectin | | uses the blended Dietary fiber carbon factor above (0.40-0.45); listed here only for its fermentability-derived utilization fraction | 95% | most rapidly/completely fermented purified fiber across in vitro fecal-fermentation comparisons |
| Carbon | Dietary fiber - inulin/FOS | | uses the blended Dietary fiber carbon factor above (0.40-0.45); listed here only for its fermentability-derived utilization fraction | 97% | the reference 'highly fermentable prebiotic fiber' in essentially every comparative in vitro fermentation study |
| Carbon | Dietary fiber - GOS | | uses the blended Dietary fiber carbon factor above (0.40-0.45); listed here only for its fermentability-derived utilization fraction | 97% | no GOS-specific in vitro fermentation figure located; assumed similar to inulin/FOS by analogy (both well-established, rapidly-fermented prebiotic oligosaccharides of similar chain length/structure) |
| Carbon | Dietary fiber - lignin | | uses the blended Dietary fiber carbon factor above (0.40-0.45); listed here only for its fermentability-derived utilization fraction | 0% | not a carbohydrate and not degraded by human colonic microbiota under normal conditions – a structural/chemical fact (an aromatic polymer), not a fitted or estimated number |

| Element | Substrate | Conversion factor | Conversion type / basis | Utilizable fraction | Utilization basis |
| --- | --- | --- | --- | --- | --- |
| — | Bicarbonate | | not converted to any tracked element | 0% | contributes no carbon, nitrogen, phosphorus, or sulfur toward biomass; assigned 0% utilization outright (see the Bicarbonate section) |

**Table S4.** Elemental biomass yields: grams of dry bacterial biomass producible per gram of each element. Central value and plausible physiological range based on the supporting literature. Elemental mass fraction indicates the inverse yield. See SI Text 10.1.

| Element | Central yield (g DW/g element) | Plausible range (g DW/g element) | Elemental mass fraction | Confidence | Key references |
|---|---|---|---|---|---|
| C, anabolic | 2.0 | 1.8–2.2 | 50% | High | [172, 171] |
| C, total | 0.89 | 0.73–1.14 | — | High; direct gut-bacterial carbohydrate uptake plus anabolic C accounting | [5, 172, 171] |
| N | 7.5 | 6–10 | 13.3% | High | [174, 171] |
| P | 40.0 | 20–100 | 2.5% | High; substantial physiological variation | [175, 177, 176, 179, 174, 178] |
| S | 200.0 | 100–300 | 0.50% | Moderate–high | [174, 171, 180] |
| Mg | 200.0 | 150–800 | 0.50% | Moderate | [174, 172, 180] |
| K | 40.0 | 20–60 | 2.5% | Moderate | [172, 180] |
| Ca | 1000.0 | 400–2,000 | 0.10% | Low–moderate | [172, 180] |
| Na | 42.6 | — (single point estimate; no independent range reported) | 2.3% | Low; single-source point estimate with no universal stoichiometric requirement | [172] |

[23] Shahneela Mazhar, Annie Simon, Joan Colom, Ekaterina Khokhlova, Martin Buckley, Christopher Phipps, John Deaton, and Kieran Rea. Acute physiological effects on macromolecule digestion following oral ingestion of the enzyme blend Elevase® in individuals that had undergone an ileostomy, but were otherwise healthy—a randomized, double blinded, placebo-controlled exploratory study. *Frontiers in Nutrition*, 11, July 2024.

[50] Vandana Sharma, Dmitry A. Rodionov, Semen A. Leyn, David Tran, Stanislav N. Iablokov, Hua Ding, Daniel A. Peterson, Andrei L. Osterman, and Scott N. Peterson. B-Vitamin Sharing Promotes Stability of Gut Microbial Communities. *Frontiers in Microbiology*, 10, July 2019.

[51] Palni Kundra, Annelies Geirnaert, Benoit Pugin, Paola Morales Martinez, Christophe Lacroix, and Anna Greppi. Healthy adult gut microbiota sustains its own vitamin B12 requirement in an in vitro batch fermentation model. *Frontiers in Nutrition*, 9, December 2022.

[59] Stefanía Magnúsdóttir, Dmitry Ravcheev, Valérie de Crécy-Lagard, and Ines Thiele. Systematic genome assessment of B-vitamin biosynthesis suggests co-operation among gut microbes. *Frontiers in Genetics*, 6, April 2015.

[60] Chiara Tarracchini, Francesca Bottacini, Leonardo Mancabelli, Gabriele Andrea Lugli, Francesca Turroni, Douwe van Sinderen, Marco Ventura, and Christian Milani. Approaches to dissect the vitamin biosynthetic network of the gut microbiota. *Microbiome Research Reports*, 4(4):N/A–N/A, October 2025.

[61] Qiushi Huang, Barbara H. Braffett, Samuel J. Simmens, Heather A. Young, and Cynthia L. Ogden. Dietary Polyphenol Intake in US Adults and 10-Year Trends: 2007-2016. *Journal of the Academy of Nutrition and Dietetics*, 120(11):1821–1833, November 2020.

[62] Raul Zamora-Ros, Viktoria Knaze, Joseph A. Rothwell, Bertrand Hémon, Aurelie Moskal, Kim Overvad, Anne Tjønneland, Cecilie Kyrø, Guy Fagherazzi, Marie-Christine Boutron-Ruault, Marina Touillaud, Verena Katzke, Tilman Kühn, Heiner Boeing, Jana Förster, Antonia Trichopoulou, Elissavet

Valanou, Eleni Peppa, Domenico Palli, Claudia Agnoli, Fulvio Ricceri, Rosario Tumino, Maria Santucci de Magistris, Petra H.M. Peeters, H. Bas Bueno-de-Mesquita, Dagrun Engeset, Guri Skeie, Anette Hjartåker, Virginia Menéndez, Antonio Agudo, Esther Molina-Montes, José María Huerta, Aurelio Barricarte, Pilar Amiano, Emily Sonestedt, Lena Maria Nilsson, Rikard Landberg, Timothy J. Key, Kay-Thee Khaw, Nicholas J. Wareham, Yunxia Lu, Nadia Slimani, Isabelle Romieu, Elio Riboli, and Augustin Scalbert. Dietary polyphenol intake in Europe: The European Prospective Investigation into Cancer and Nutrition (EPIC) study. *European journal of nutrition*, 55(4):1359–1375, June 2016.

[92] Kanapathipillai Wignarajah, Suresh Pisharody, and John W. Fisher. Investigating the Partitioning of Inorganic Elements Consumed by Humans between the Various Fractions of Human Wastes: An Alternative Approach, January 2003.

[99] Institute of Medicine. *Dietary Reference Intakes for Calcium, Phosphorus, Magnesium, Vitamin D, and Fluoride*. The National Academies Press, Washington, DC, 1997.

[134] Angela M. Patterson and Alastair J. M. Watson. Deciphering the Complex Signaling Systems That Regulate Intestinal Epithelial Cell Death Processes and Shedding. *Frontiers in Immunology*, 8, July 2017.

[135] Ron Sender and Ron Milo. The distribution of cellular turnover in the human body. *Nature Medicine*, 27(1):45–48, January 2021.

[136] Herbert F Helander and Lars Fändriks. Surface area of the digestive tract – revisited. *Scandinavian Journal of Gastroenterology*, 49(6):681–689, June 2014.

[137] Paulus G. M. Jochems, Johan Garssen, Antonius M. van Keulen, Rosalinde Masereeuw, and Prescilla V. Jeurink. Evaluating Human Intestinal Cell Lines for Studying Dietary Protein Absorption. *Nutrients*, 10(3):322, March 2018.

[138] Dmitry A. Ravcheev and Ines Thiele. Systematic genomic analysis reveals the complementary aerobic and anaerobic respiration capacities of the human gut microbiota. *Frontiers in Microbiology*, 5, December 2014.

[165] D. Breese (David Breese) Jones. *Factors for Converting Percentages of Nitrogen in Foods and Feeds into Percentages of Proteins*. Washington, D.C. : U.S. Dept. of Agriculture, 1931.
